# Netrin G1 Ligand is a new stromal immunomodulator that promotes pancreatic cancer

**DOI:** 10.1101/2024.05.15.594354

**Authors:** Débora B. Vendramini-Costa, Ralph Francescone, Janusz Franco-Barraza, Tiffany Luong, Myree Graves, Ariana Musa de Aquino, Nina Steele, Jaye C. Gardiner, Sérgio Alexandre Alcantara dos Santos, Charline Ogier, Emily Malloy, Leila Borghaei, Esteban Martinez, Dmitry I. Zhigarev, Yinfei Tan, Hayan Lee, Yan Zhou, Kathy Q. Cai, Andres J. Klein-Szanto, Huamin Wang, Mark Andrake, Roland L. Dunbrack, Kerry Campbell, Edna Cukierman

**Affiliations:** Cancer Signaling and Microenvironment; Marvin and Concetta Greenberg Pancreatic Cancer Institute; Nuclear Dynamics in Cancer and Microenvironment, Fox Chace Cancer Center, Temple Health, Philadelphia, PA; Henry Ford Pancreatic Cancer Center, Henry Ford Health; Henry Ford Health + Michigan State University Health Sciences, Detroit, MI; Department of Anatomical Pathology, The University of Texas MD Anderson Cancer Center, Houston, TX

**Author notes:** Corresponding author Edna Cukierman, Fox Chase Cancer Center. 333 Cottman Avenue, Philadelphia, PA. 19111. Equal contribution (Débora B. Vendramini-Costa, Ralph Francescone). **Present address** Charline Ogier: Centre de recherches en cancérologie de Toulouse, Toulouse, France. Dmitry Zhigarev: The Wistar Institute. Philadelphia, PA.

**Keywords:** NGL-1, *LRRC4C*, pancreatic cancer, cancer-associated fibroblasts, tumor microenvironment, Immunosuppression

## Abstract

Understanding pancreatic cancer biology is fundamental for identifying new targets and for developing more effective therapies. In particular, the contribution of the stromal microenvironment to pancreatic cancer tumorigenesis requires further exploration. Here, we report the stromal roles of the synaptic protein Netrin G1 Ligand (NGL-1) in pancreatic cancer, uncovering its pro-tumor functions in cancer-associated fibroblasts and in immune cells. We observed that the stromal expression of NGL-1 inversely correlated with patients’ overall survival. Moreover, germline knockout (KO) mice for NGL-1 presented decreased tumor burden, with a microenvironment that is less supportive of tumor growth. Of note, tumors from NGL-1 KO mice produced less immunosuppressive cytokines and displayed an increased percentage of CD8^+^ T cells than those from control mice, while preserving the physical structure of the tumor microenvironment. These effects were shown to be mediated by NGL-1 in both immune cells and in the local stroma, in a TGF-β-dependent manner. While myeloid cells lacking NGL-1 decreased the production of immunosuppressive cytokines, NGL-1 KO T cells showed increased proliferation rates and overall polyfunctionality compared to control T cells. CAFs lacking NGL-1 were less immunosuppressive than controls, with overall decreased production of pro-tumor cytokines and compromised ability to inhibit CD8^+^ T cells activation. Mechanistically, these CAFs downregulated components of the TGF-β pathway, AP-1 and NFAT transcription factor families, resulting in a less tumor-supportive phenotype. Finally, targeting NGL-1 genetically or using a functionally antagonistic small peptide phenocopied the effects of chemotherapy, while modulating the immunosuppressive tumor microenvironment (TME), rather than eliminating it. We propose NGL-1 as a new local stroma and immunomodulatory molecule, with pro-tumor roles in pancreatic cancer.

**Statement of Significance:** Here we uncovered the pro-tumor roles of the synaptic protein NGL-1 in the tumor microenvironment of pancreatic cancer, defining a new target that simultaneously modulates tumor cell, fibroblast, and immune cell functions. This study reports a new pathway where NGL-1 controls TGF-β, AP-1 transcription factor members and NFAT1, modulating the immunosuppressive microenvironment in pancreatic cancer. Our findings highlight NGL-1 as a new stromal immunomodulator in pancreatic cancer.

## Introduction

Pancreatic ductal adenocarcinoma (PDAC) bears one of the most devastating 5-year overall survival (OS) rates among all types of cancer, at only 13% (1). Unsurprisingly, it is projected to become the second leading cause of cancer-related death in the United States by 2026 (2). Often termed as a “silent disease”, due to lack of early detection markers and manifestation of symptoms, most of the patients (∼80%) are diagnosed with locally advanced and/or metastatic disease at the time of diagnosis (3). Contributing to the dismal statistics, its anatomical location impedes ease of diagnosis, as well as the lack of effective therapies and other interventions for pancreatic cancer (4).

PDAC is a unique cancer; in addition to lacking markers for early detection, it features a highly fibrotic stroma and low mutational burden (5–8). These features culminate in a very immunosuppressive environment, poor accessibility of drugs and nutrients from the blood stream, and lack of immunogenicity, which contributes to resistance to chemotherapy and immunotherapy and is susceptible to developing chemoresistance (9,10). One of the hallmarks of PDAC is the fibrotic stroma that arises from activated fibroblasts and their deposited extracellular matrix (ECM) (11–13). This stromal expansion, known as desmoplasia, shapes the PDAC microenvironment (8,14). Together with immunosuppressive immune cells and limited cytotoxic cells (e.g., T and NK cells), the desmoplastic stroma can comprise up to 90% of PDAC tissue (15). While desmoplasia is easily identified histologically due to the significant ECM deposition, the biology behind pro-tumoral fibroblastic activation and expansion, and how this unique tumor microenvironment (TME) is formed, are not well understood (16–18). Paradoxically, attempts to eliminate the PDAC TME lead to more aggressive tumors in both mice and patients; the dense stroma represents a barrier for immune cell infiltration and drug penetration, but it is also a barrier for the prompt spread of cancer cells (19–23). Recognizing the inherent tumor suppressive potential of the TME and means to leverage its function through normalization is predicted to be a more effective approach (24). Therefore, uncovering novel stromal targets and their modulation strategies towards normalization deserves immediate attention for future PDAC therapies.

As part of this effort, our group has been focused on understanding the complex biology of cancer associated fibroblasts (CAFs) and their ECM (defined as CAF/ECM units) (25,26). We reported a pro-tumor CAF/ECM unit signature that involves the activation of αvβ5-integrin, sustaining constitutive expression of the tyrosine 397 phosphorylated focal adhesion kinase (p-FAK), and the upregulation of the actin-binding protein Palladin, in a canonical TGF-β dependent manner (12,27,28). Moreover, comparing functional tumor adjacent fibroblasts and CAF/ECM units from PDAC patients, we uncovered the ectopic expression of the glutamatergic synaptic protein Netrin G1 (NetG1) in CAFs, and its sole binding partner, the receptor Netrin G1 Ligand (NGL-1) in pancreatic cancer cells (29). NetG1 in CAFs controlled their immunosuppressive functions and their ability to provide metabolic support for cancer cells, while NGL-1 in cancer cells regulated their capacity for up-taking nutrients from the TME via macropinocytosis. Moreover, ablation of NetG1 in CAFs, or NGL-1 in cancer cells, reduced tumor burden in mice, and fibroblastic expression of NetG1 inversely correlated with overall patient survival (29,30). Our group also reported that extracellular vesicles produced by NetG1-deficient CAFs hindered PDAC survival compared to control CAFs. Blocking NetG1/NGL-1 interaction further limited extracellular vesicle-mediated PDAC survival (31), solidifying the role for these synaptic proteins in PDAC.

In view of the importance of neuronal signatures in PDAC (32–35) and specifically of the functional and translational impact of the discovery of NetG1 and NGL-1 in pancreatic cancer, we sought to further explore the roles of NGL-1. Here we demonstrated that the stromal expression of NGL-1 inversely correlates with PDAC patient’s overall survival. Moreover, NGL-1 KO mice develop smaller tumors, with a less immunosuppressive environment, where NGL-1 expression in both the local stroma and recruited immune cells were important for tumor development. Functionally, NGL-1 modulates the pro-tumor phenotype of CAFs and macrophages, while inhibiting the antitumor phenotype of cytotoxic T cells. Mechanistically, we report a signaling circuit in CAFs where NGL-1 modulates the expression of members of the TGF-β pathway, AP-1 transcription factor family, and the nuclear factor of activated T cells (NFAT1). NGL-1 ablation results in decreased production of immunosuppressive cytokines, and decreased support for cancer cell survival, while maintaining the physical structure of the TME. Moreover, the genetic deletion of NGL-1 phenocopied the effect of chemotherapy *in vivo*, without side effects, and a small peptide that inhibits NGL-1 signaling also led to smaller tumors *in vivo*. Collectively, this underscores the potential of NGL-1 to be a global target in PDAC due to its expression in different cell compartments (fibroblasts, immune cells, and cancer cells). Therefore, this study reports the novel roles of a synaptic protein, NGL-1, in the pro-tumoral immunomodulation of the PDAC TME, offering an innovative new target for such a devastating disease.

## Results

### NGL-1 expression in CAFs inversely correlates with PDAC patients’ overall survival

In order to verify the expression of NGL-1 in the TME in PDAC, we performed simultaneous multichannel immunofluorescence in human pancreatic tissues (6 normal, 4 tumor-adjacent and 15 tumor samples). These were the same human samples analyzed in our previous study evaluating the expression patterns of NetG1 in fibroblasts/CAFs and NGL-1 in normal/tumor epithelial cells (29). NGL-1 expression was upregulated in the TME (stroma, defined by pan-cytokeratin negative cells) of PDAC patients’ samples compared to normal pancreatic tissue from healthy donors (Fig. 1A and B). The levels of NGL-1 expression in stromal cells were higher in the tumor adjacent (TA) and tumoral tissue (Tumor) compared to pancreatic tissue from healthy donors (Normal), suggesting that its expression in the stroma correlates with the presence of disease in the pancreas (Fig. 1A and B). The expression of NGL-1 in the stroma correlated weakly with the previously reported expression of NGL-1 (29) in the epithelial compartment (R^2^ = 0.3312 p < 0.001) (Supplementary Fig. 1A). Similarly, the expression of the tyrosine 397 phosphorylated focal adhesion kinase p-FAK and NetG1, both found to be upregulated in fibroblasts from PDAC patients compared to samples from healthy donors (12,29,36), moderately correlated with the stromal expression of NGL-1 (R^2^ = 0.4301, p < 0.001 and 0.4690 p < 0.001 respectively) (Sup Figs. 1B-C). These results suggest that NGL-1 expression in the TME is increased in PDAC and correlates with known markers of a pro-tumoral microenvironment.

**Figure 1:**
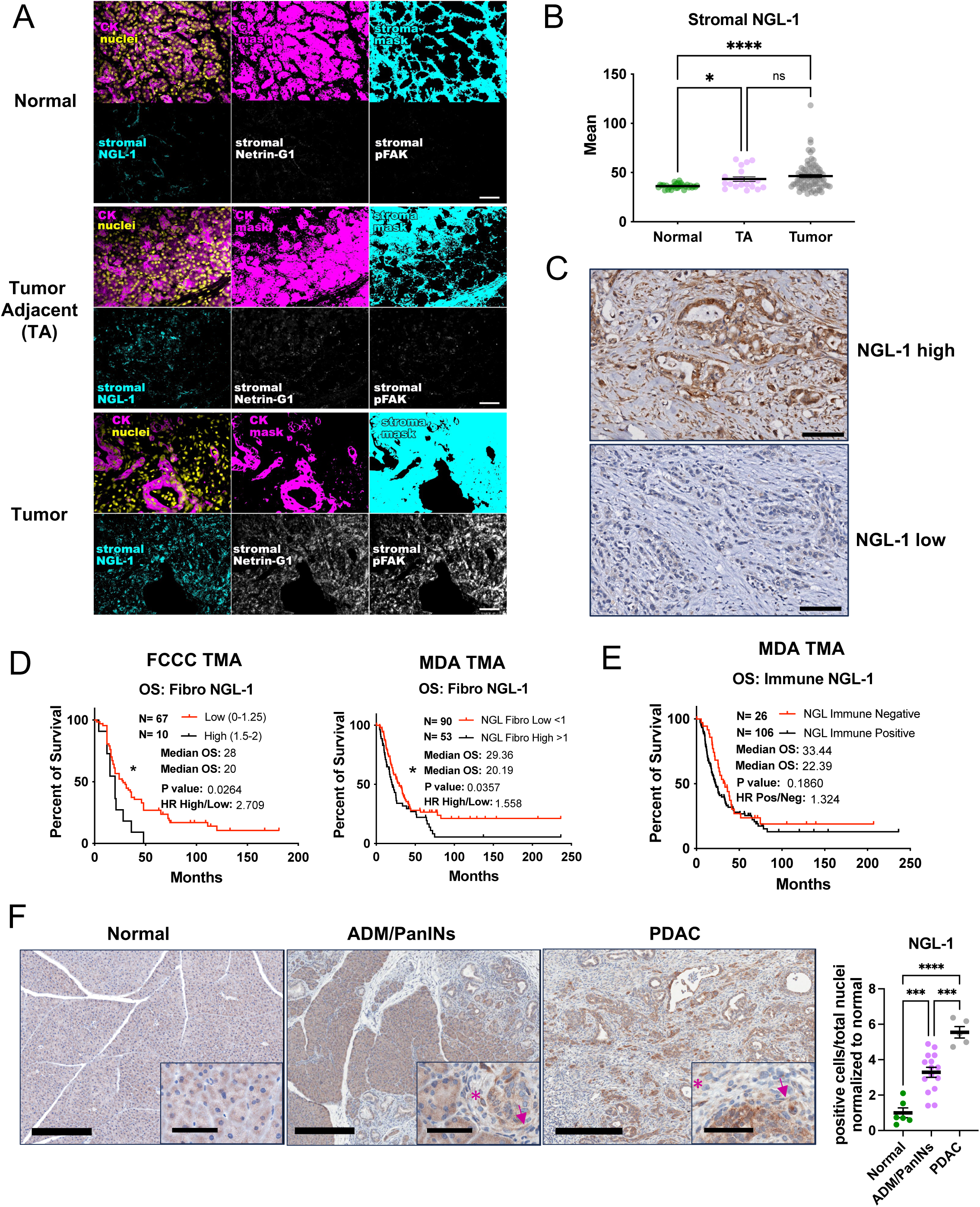
Stromal expression of NGL-1 correlates with disease progression and inversely correlates with patient’s overall survival. **A)** Representative SMI images for NGL-1, Netrin G1 and p-FAK in normal human pancreas, tumor adjacent (TA) and tumoral tissue of PDAC patient surgical samples. The top rows show pseudo-colored images that display the following masks: pan-cytokeratin positive for epithelial compartment (CK, magenta), DRAQ5 positive for nuclei (nuclei, yellow) and vimentin positive for stromal compartment (stromal, cyan). The lower rows show the expression of markers of interest in the stroma (defined as areas negative for CK). NGL-1 in cyan, Netrin G1 and p-FAK in gray. Scale Bar: 50 µm. **B)** Quantification of the expression (mean fluorescence intensity) of NGL-1 in the stroma (defined as pan-cytokeratin negative areas) of normal pancreas (n = 6, green), TA (n = 4, violet) and tumor areas (n = 15, gray). One-way ANOVA, Dunnett’s T3 multiple comparisons test. **C)** Immunohistochemistry for NGL-1 in a representative core from a PDAC patient tumor microarray, comparing a patient with high NGL-1 expression (NGL-1 high) and low expression (NGL-1 low). Scale bar: 100 µm. **D)** Kaplan Meier curves depicting the percentage of overall survival in months of PDAC patients from two independent cohorts: from Fox Chase Cancer Center (FCCC, n = 77) and from MD Anderson (MDA, n = 153). Patients were stratified as expressing low or high levels of NGL-1 in fibroblasts, according to histopathological scores of immunohistochemistry for NGL-1 in tissue microarrays. OS: overall survival in months. HR: hazard ratio. Log-rank test was used to determine statistical significance. **E)** Kaplan Meier curves depicting the percentage of overall survival in months of PDAC patients from MD Anderson (MDA, n = 132). Patients were stratified as negative or positive for NGL-1 in immune cells, according to histopathological scores of immunohistochemistry for NGL-1 in tissue microarrays. OS: overall survival in months. HR: hazard ratio. Log-rank test was used to determine statistical significance. **F)** Representative images and quantification of the expression of NGL-1 in normal pancreatic tissue (n = 6, age 2 months old), in early pancreatic lesions (n = 16, age 3 months old) and in PDAC tissue (n = 5, age 4 months old) of KPC mice (Kras^LSL-G12D/+^; p53^flox/WT^;Pdx-Cre/^+^). One-Way ANOVA, Dunnett’s T3 multiple comparisons test. ADM: acinar to ductal metaplasia. PanINs: pancreatic intraepithelial neoplasia. Scale bar: 300 µm. Insert: 60 µm. Arrow: fibroblast. Asterisk: immune cell. All graphs depict mean ± standard error of the mean (SEM). * p < 0.05, ** p < 0.01, *** p < 0.001, **** p < 0.0001. ns: not significant.

To evaluate the clinical relevance of NGL-1 expression in the TME of PDAC, immunohistochemistry (IHC) was performed for NGL-1 using tissue microarrays generated from two independent cohorts of PDAC patients; a cohort of 80 patients from Fox Chase Cancer Center (FCCC) and a second cohort of 153 patients from MD Anderson (MDA), totaling 233 patients (Fig. 1C, Sup. Table 1). Of note, these were the same cohorts used in our previous study (29). Interestingly, NGL-1 expression in fibroblasts inversely correlated with patients’ overall survival (OS) across these two independent cohorts of PDAC patients (Fig. 1D). In the FCCC cohort, patients with low NGL-1 had a median OS of 28 months, compared to 20.2 months for those with high levels (p = 0.0264, HR 2.701). In the MDA cohort, patients with low NGL-1 had a median OS of 29.36 months, compared to 20.19 months for those with high NGL-1 levels (p = 0.0357, HR 1.558) (Fig. 1D). Similar trends were noted for NGL-1 expression in immune cells (p = 0.1860, median OS NGL-1 positive vs NGL-1 negative 22.39 vs 33.44 months, HR 1.324) (Fig. 1E). Most patients (83.5%) had positive expression of NGL-1 in immune cells (26 negative and 132 positive), suggesting that NGL-1 upregulation in immune cells might be an early event during PDAC progression.

Next, to confirm our findings from the human PDAC tissue, we analyzed pancreatic tissue from the murine pancreatic cancer models KC (Pdx1-Cre; Kras^LSL-G12D/+^) and KPC (Kras^LSL-G12D/+^; p53^flox/WT^;Pdx-Cre/^+^) (37,38) collected at different time points, representing different lesion stages. KC animals were harvested between 1.5 and 12 months of age, while KPC animals were harvested at 15 weeks and 20-28 weeks of age. The tissues were reviewed by a pathologist, and the samples were stained for NGL-1 by IHC. Samples were divided according to the presence and stage of the lesions: absence of lesions (i.e., normal), presence of the pre-cancerous lesions acinar to ductal metaplasia ADM, and pancreatic intraepithelial neoplasms (PanINs), but absence of PDAC lesions (i.e., ADM/PanINs); and presence of PDAC (PDAC). As shown in Sup. Fig. 1D (KC mice) and Fig. 1F (KPC mice), NGL-1 expression increases according to the disease state, with higher expression associated with pancreatic lesions (about 3.5-fold increase), and most highly expressed in tissue bearing PDAC (about 5.5-fold) (Fig 1F), when compared to normal pancreas controls. Note the expression of NGL-1 in fibroblasts (arrow, Fig. 1F and Sup. Fig. 1D) and in immune cells (asterisks, Fig. 1F and Sup. Fig. 1D). We also confirmed these observations using a probe designed to detect a short RNA sequence of the *Lrrc4c* gene (encoding NGL-1) by BaseScope assay, where the expression of NGL-1 is higher in PDAC tissue compared to tissue presenting ADM (Sup. Fig. 2A). Our results show that NGL-1 is expressed in the pathologic PDAC TME, correlating with disease progression.

To further confirm that fibroblasts and immune cells express NGL-1 in the context of PDAC, we isolated CAFs from four different patients, and probed NGL-1 expression via western blotting. All CAF/ECM units tested demonstrated robust expression of NGL-1 (Sup. Fig. 2B). Next, we sought to identify which immune cells were expressing NGL-1 in mice bearing pancreatic tumors. We sorted neutrophils (CD11b^+^ and F4/80 negative (neg), then Ly6C^low/neg^ and Ly6G^+^), monocytes (CD11b^+^ and F4/80^neg^, then Ly6C^+high^), macrophages (F4/80^+^), CD4^+^ and CD8^+^ T cells (CD3^+^ and CD4^+^ or CD8^+^) and NK cells (CD3^neg^ and NK1.1^+^) (Sup. Fig. 2C for sorting strategy) from the spleens of naïve (without pancreatic tumors, 5 months old) and tumor bearing mice (KPC mice, 5.5 months old) and determined NGL-1 expression by real time PCR (RT-PCR). Interestingly, nearly all immune cells from tumor bearing mice tended to overexpress NGL-1 compared to those from naïve mice (Sup. Fig. 2D). We also confirmed this at the protein level, as western blotting of the lysates obtained from sorted neutrophils and CD8^+^ T cells from the spleens of 6- and 9-months old KC and 3.5- and 5.5-months old KPC mice showed expression of NGL-1 in all samples (Sup. Fig. 2E), suggesting a role for NGL-1 in immune cells.

### Stromal NGL-1 shapes a pro-tumor pancreatic microenvironment

Next, we questioned if ablation of NGL-1 in the TME could restrain pancreatic cancer tumorigenesis *in vivo*. We performed syngeneic orthotopic injections of pancreatic cancer cells (29) in wild type mice, germline NGL-1 heterozygous mice (NGL-1 het) and germline NGL-1 knockout mice (NGL-1 KO), ages 3 to 6 months, and assessed tumor burden. The genetic deletion of NGL-1 was confirmed by DNA genotyping (Sup. Fig. 3A) and mRNA expression by BaseScope assay in the brains (tissue where NGL-1 expression is reported) of WT and NGL-1 KO mice (Sup. Fig. 3B)(39,40). We used a BaseScope assay probe that targets the specific RNA sequence that is deleted in the NGL-1 KO mice, therefore the signal is absent in the NGL-1 KO tissue (Sup. Fig. 3B).

Remarkably, we observed that the NGL-1 het and KO tumor-bearing mice presented diminished pancreata weight compared to WT littermates (Fig. 2A; normalized pancreas weight WT: 1 ± 0.23; NGL-1 het: 0.78 ± 0.08 p < 0.05; NGL-1 KO: 0.69 ± 0.05 p < 0.01), indicative of less tumor burden in these animals (Fig. 2A). Because pancreatic cancer is typically more frequently diagnosed in elderly patients, we performed the same orthotopic syngeneic injections in WT, NGL-1 het and KO mice, 14 months old, and assessed tumor burden. The NGL-1 het and KO mice presented decreased tumor weight compared to WT animals (Sup. Fig. 3C; normalized pancreas weight WT: 1 ± 0.33; NGL-1 het: 0.53 ± 0.18 p < 0.05; NGL-1 KO: 0.62 ± 0.29 p < 0.05), and tumors were mostly poorly differentiated in all the groups, independent of the age (Sup. Fig. 3D and 3E). These results indicate that targeting stromal NGL-1 in patients of different ages would result in an antitumor benefit, despite the increased susceptibility of the aged pancreas for the development of tumors (10,41,42).

**Figure 2:**
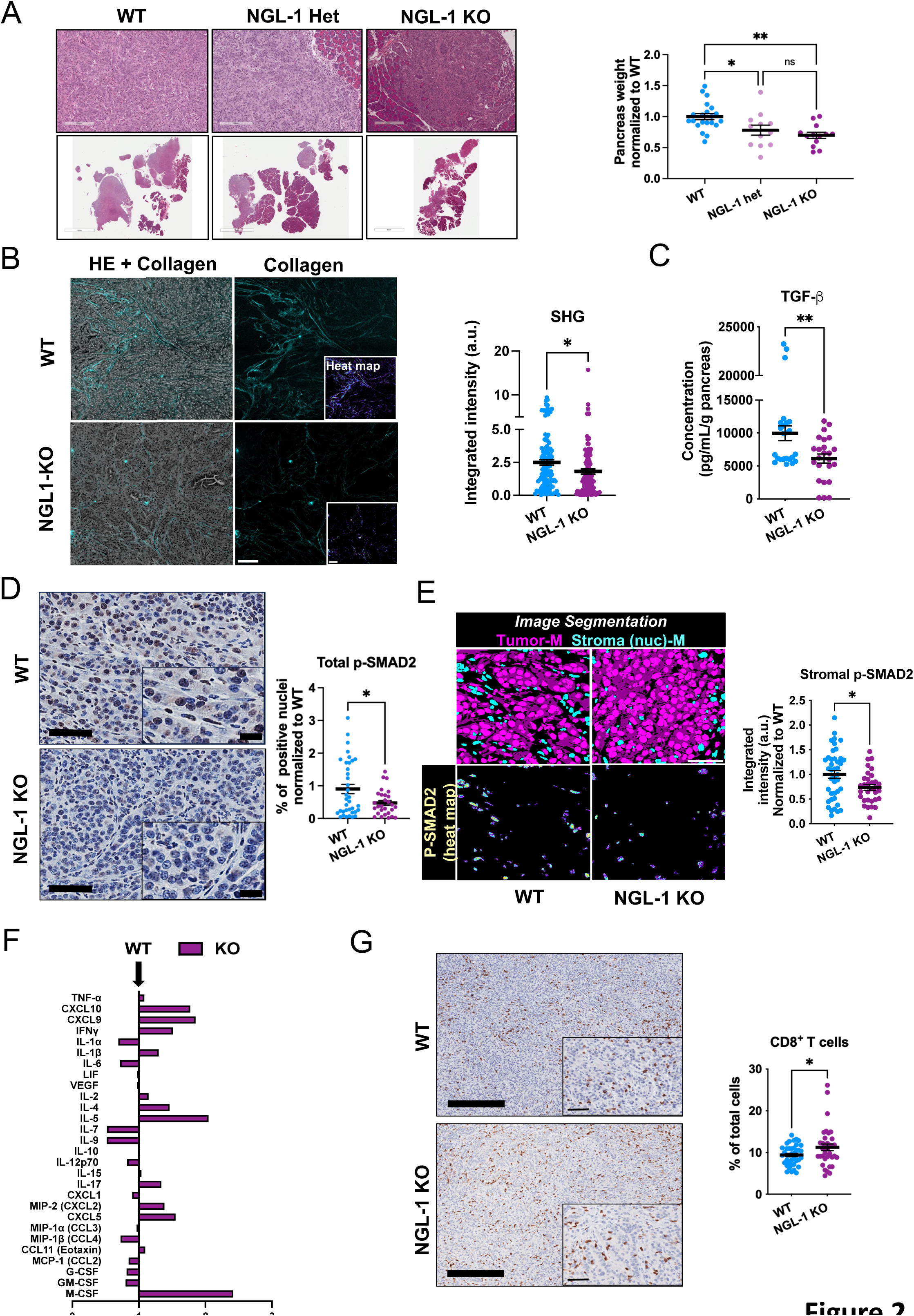
Germline deletion of NGL-1 results in reduced tumor burden and inhibition of pro-tumor microenvironmental features in orthotopic murine models of PDAC. **A)** Representative images of the pancreas of mice germline wild type (WT), heterozygous (NGL-1 Het) and knockout (NGL-1 KO) orthotopically allografted with a murine PDAC cell line (KPC3), stained with H&E. Scale bars: 6mm and 200 µm. Graph showing the pancreas weight as a measurement of tumor burden in the different groups. WT in blue (n = 21), NGL-1 Het in violet (n = 12) and NGL-1 KO in purple (n = 13). The pancreas weight was normalized to the average weight of the WT group. One-Way ANOVA, Tukey’s multiple comparisons test. **B)** Representative images of PDAC tissue from WT and NGL-1 KO mice (H&E) overlayed with corresponding second harmonic generation (SHG) of the collagen signatures, to detect fibrillar collagen, pseudo colored in cyan (right panel). The insert represents the heatmap of the pixel intensity derived from the SHG signal, with warmer tones representing higher and cooler tones representing lower pixel intensity. Scale bars: 100 µm. The quantification of the integrated intensity of fibrillar collagen is presented in the graph. Unpaired t test, N = 7 mice/group. **C)** ELISA detecting the production of the cytokine TGF-β by fragments of pancreatic tissue from WT (n = 8) and NGL-1 KO mice (n = 8). The fragments of pancreas were weighed and cultured for 24 hours in DMEM media without serum. TGF-β concentration was calculated by pg/mL, normalized to the weight of each fragment. Unpaired t test. **D)** Representative images of PDAC tissue from WT and NGL-1 KO mice stained for p-Smad2 (in brown) and counterstained with hematoxylin. Scale bars: 60 µm. Inserts: 34 µm. The quantification of the percentage of positive nuclei normalized to the average of the WT group is presented in the graph. WT (n = 9 mice) and NGL-1 KO (n = 7 mice). Unpaired t test. **E)** Representative SMI images of PDAC tissue from WT and NGL-1 KO mice stained for p-SMAD2 (bottom panel, heat map of the integrated intensity), under the masks (-M, top panel) tumor nuclei (-nuc, DRAQ5^+^ in cytokeratin^+^ cells, magenta) and stroma nuclei (DRAQ5^+^ in cytokeratin negative cells, cyan). Scale bar: 50 µm. The quantification of the integrated intensity of p-Smad2 normalized to the average of the WT group is presented in the graph. N = 3 mice/group. Unpaired t test. **F)** Multiplex cytokine array displayed as bar graphs of the fold change relative to the average of the WT group (levels in the WT group were set as “1”, arrow). Bars to the right (values higher than 1) represent cytokines that are upregulated by the NGL-1 KO group, while bars to the left (values smaller than 1) represent cytokines that are downregulated in the NGL-1 KO group compared to WT group. N = 6 mice per group. **G)** Representative images of PDAC tissue from WT and NGL-1 KO mice stained for CD8 (to detect CD8^+^ T cells, in brown) and counterstained with hematoxylin. Scale bars: 300 µm. Insert: 60 µm. The quantification of the percentage of CD8^+^ T cells is presented in the graph. N = at least 7 mice/group. Unpaired t test. All graphs depict mean ± SEM. * p < 0.05, ** p < 0.01. ns: not significant.

Given that NGL-1 het and KO mice presented statistically similar pancreata weights, we further characterized tumors growing in the KO mice compared to those growing in the WT mice, from the younger cohorts. There were no differences in the frequency of cell death, measured by the terminal deoxynucleotidyl transferase-mediated dUTP nick end labeling (TUNEL) staining and cleaved caspase 3 expression, between WT and NGL-1 KO tumors (Sup. Fig. 3F and G). Nonetheless, we noted a tendency for decreased staining for Ki67 in tumors from NGL-1 KO mice, suggesting a trend for less proliferative cells than their WT counterparts (Sup. Fig. 3H).

Since PDAC is known for its desmoplastic expansion, we sought to characterize the fibroblastic stroma of tumors from WT and NGL-1 KO mice. Analysis of the collagen architecture by second harmonic generation (SHG) of polarized light showed that NGL-1 KO tumors presented with reduced collagen bundling, suggesting less interstitial ECM maturation compared to WT tumors (Fig. 2B). Because one of the major pathways involved in pro-tumoral ECM remodeling and pancreatic cancer development is the canonical TGF-β pathway, we analyzed the levels of TGF-β secreted by WT and NGL-1 KO tumors, as well as the expression levels of phosphorylated Smad2 (p-Smad2). Accordingly, NGL-1 KO tumors secreted less TGF-β compared to WT tumors (Fig. 2C) and presented reduced expression of p-Smad2 in general (Fig. 2D), and specifically in the stroma (Fig. 2E). Conversely, the levels of other known pro-tumor stromal markers such as p-FAK and NetG1 remained unchanged (Sup. Fig. 3I), which is in accordance with the PDAC patient data where we observed a weak to moderate correlation between stromal NGL-1 levels and p-FAK/NetG1 expressions (Sup. Fig. 1C and D).

Another hallmark of pancreatic cancer is its immunosuppressive microenvironment (9), characterized by the presence of immunosuppressive myeloid cells, CAFs and T regulatory cells (Tregs) and decreased frequencies and dysfunction of cytotoxic lymphocytes (43). To assess if NGL-1 modulates this highly suppressive environment, we first analyzed an array of cytokines produced by the tumors of the WT and NGL-1 KO mice, using Luminex assay (Sup. Table 2). Evaluation of the differential production of cytokines showed that the tumor microenvironment in NGL-1 KO mice produced more IFNγ, CXCL9 and CXCL10 than in WT mice (Fig. 2F), revealing a proinflammatory environment with improved immunosurveillance through recruitment and stimulation of cytotoxic T cells and NK cells, an indication of better response to immunotherapy (44). These changes were also accompanied by an increase in TNFα and IL-2 production by tumors growing in NGL-1 KO hosts compared to WT. TNFα is involved in the activation and function of cytotoxic T cells and inflammatory macrophages (45–47), while IL-2 promotes the expansion of T cells (48). Moreover, we observed increased production of Th2/Th17 related cytokines by the NGL-1 KO tumors (IL-4, IL-5, IL-17), which typically correlates with worse outcomes (49,50). Interestingly, while the tumors in the NGL-1 KO mice produced decreased levels of the chemokines CCL2, CCL3, CCL4 and CCL11, and of the growth factors G-CSF and GM-CSF, these same tumors displayed increased production of M-CSF and of the chemokines CXCL5 and CXCL2. M-CSF stimulates differentiation, proliferation and survival of monocytes and macrophages, while the chemokines play a role in the recruitment, expansion and functionality of polymorphonuclear granulocytes (neutrophils and myeloid derived suppressor cells) (51–54). These data suggest that tumors developing in NGL-1 KO mice did not lose the ability to recruit myeloid cells. However, levels of IL-6 (Fig 2F and Sup. Fig. 3J) and TGF-β (Fig 2C) (important pro-tumor cytokines in pancreatic cancer) were decreased in the tumors found in NGL-1 KO mice, compared to those in WT littermates (55). These data, together with the low tumor burden in the NGL-1 KO mice, suggest that despite the number of myeloid cells being potentially similar between the two groups, their pro-tumor functions might be attenuated in NGL-1 KO mice.

In support of this premise, staining tumors from WT and NGL-1 KO mice for assorted immune cell subsets revealed that the percentages of CD4^+^ T cells, Tregs (FoxP3^+^ cells), NK cells (NK1.1^+^) and macrophages (F4/80^+^) tended to be similar between these hosts (Sup. Fig 4A). Yet, the tumors from NGL-1 KO mice presented higher percentages of the cytotoxic CD8^+^ T cells (Fig. 2G), suggesting a shift towards a less tumor-permissive TME. Flow cytometry analysis of the dissociated tumors from WT and NGL-1 KO hosts showed the same trends, including similar percentages of total myeloid cells (CD11b^+^), total and antigen presenting dendritic cells (CD11c^+^ and CD11c^+^, MHCII^+^), total and antigen presenting macrophages (F4/80^+^ and F4/80^+^, MHCII^+^) and podoplanin (PDPN)^+^ fibroblasts (CD45^neg^, then EpCAM^neg^ and e-cadherin^neg^, PDPN^+^) (Sup. Fig 4B and C). There was a tendency for higher percentages of monocytes (CD11b^+^, Ly6C^high^) in the NGL-1 KO tumors compared to the WT, which indicates increased recruitment of these cells to the tumor. This is in accordance with the increased production of M-CSF by these tumors (Fig. 2F). The percentages of the different immune cells isolated from the bone marrow and spleens of WT and NGL-1 KO tumor bearing mice were similar (Sup. Fig. 5A-B). Taken together, our data suggest that NGL-1 modulates the immunosuppressive PDAC TME to promote tumor growth, likely by affecting the functionality of the stromal cell types, rather than preventing influx of pro-tumor immune cells.

### Expression of NGL-1 in both recruited immune cells and in the local stroma is needed for its TGF-β dependent pro-tumor effects

To determine the contribution of immune cells versus other stromal cells for the pro-tumor effects of NGL-1, we generated bone marrow (BM) chimeric mice, transplanting WT BM into NGL-1 KO mice and vice versa. As a control, WT mice were transplanted with WT BM (see scheme in Fig. 3A). The efficiency of the chimerism was confirmed by flow cytometry analyzes of the blood of the transplanted mice 10 weeks after transplantation, where over 97% of the total CD45^+^ immune cells were originated from the donor BM, signifying the success of the transplantation (Sup. Fig. 6A). We performed orthotopic PDAC cell allografts in all mice, 12 weeks post-transplant of BM. The efficiency of the chimerism was again confirmed to be over 97% after 2.5 weeks following the injection of tumor cells (Sup. Fig. 6B).

**Figure 3:**
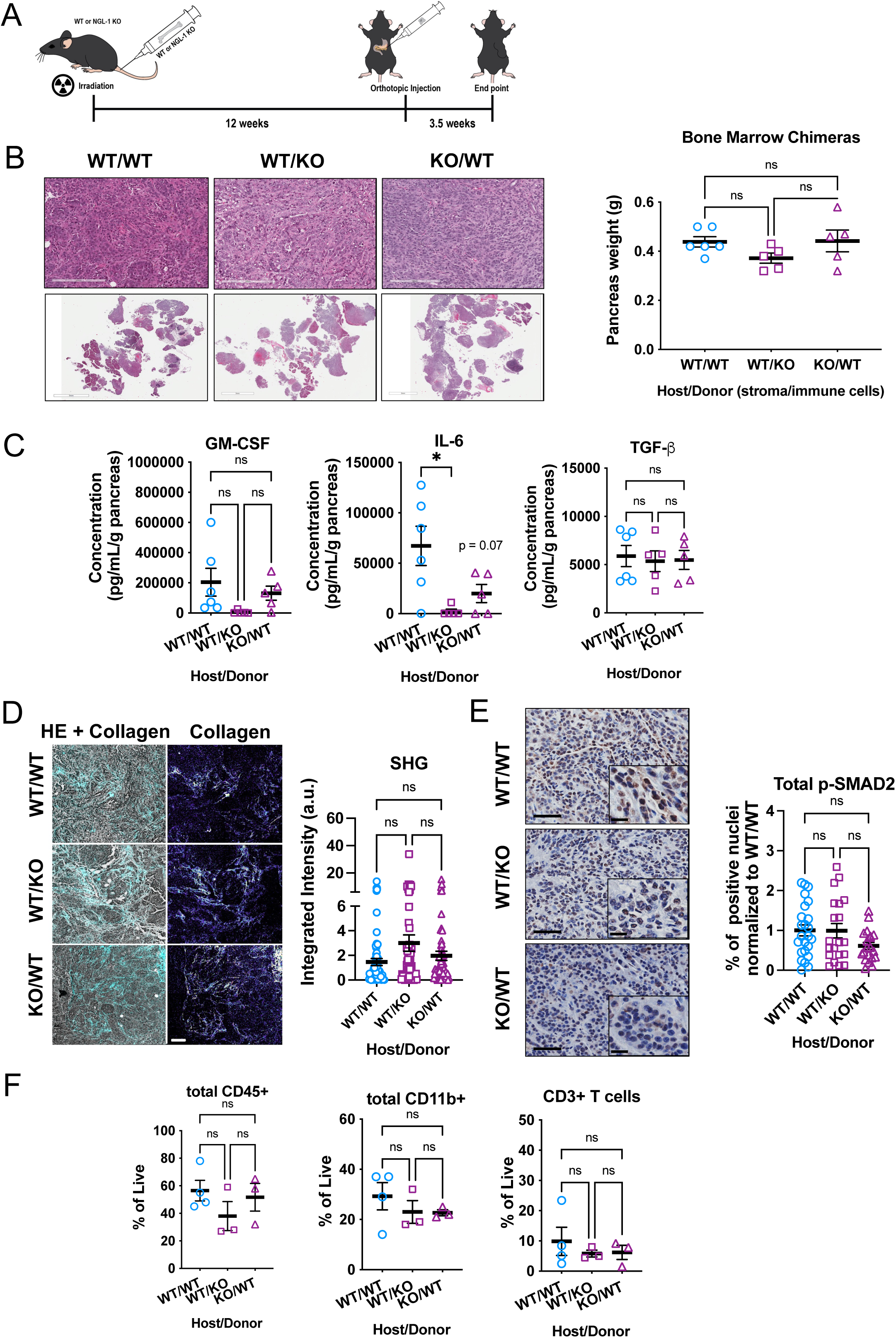
Bone marrow chimeras demonstrate that both stromal and immune expression of NGL-1 promotes PDAC tumorigenesis and correlates with TGF-β expression. **A)** Scheme of the experimental design for the bone marrow chimera experiment. WT, NGL-1 KO and B6 CD45.1 mice were subjected to lethal irradiation, and after 24 hours, they were either transplanted with or served as donors of bone marrow from NGL-1 WT or KO mice. After 12 weeks and confirmed chimerization, the mice were orthotopically allografted with PDAC cells, and 3.5 weeks later, the mice were euthanized and the tumors removed. **B)** Representative images of the pancreas of WT mice transplanted with either NGL-1 WT bone marrows (WT/WT) or NGL-1 KO bone marrow (WT/KO) and NGL-1 KO mice transplanted with WT bone marrow (KO/WT), stained with H&E. Scale bars: 200 µm. Inserts: 7 mm. The assessment of pancreas weight (g) shows that the expression of NGL-1 in either compartment (local stroma and recruited immune cells) promotes the development of pancreatic tumors. N = 5 - 6 mice/group. One-Way ANOVA, Tukey’s multiple comparisons test. **C)** ELISA detecting the production of the cytokines GM-CSF, IL-6 and TGF-β by fragments of pancreatic tissue from mice of the different groups, with TGF-β levels being recovered when NGL-1 is present in one of the compartments (local stroma or recruited immune cells). The fragments of pancreas were weighed and cultured for 24 hours in DMEM media without serum. The cytokine concentration was calculated (pg/mL), normalized to the weight of each fragment. One-Way ANOVA, Tukey’s multiple comparisons test. **D)** Representative images of PDAC tissue from WT/WT, WT/KO and KO/WT mice (H&E) overlayed with the SHG of the collagen, to detect fibrillar collagen, pseudo colored in cyan (left panel) or fibrillar collagen (right panel), with warmer tones representing higher pixel intensity and cooler tones representing lower pixel intensity. Scale bars: 100 µm. The quantification of the integrated intensity of fibrillar collagen is presented in the graph. Note that now the levels of fibrillar collagen are similar between all groups when NGL-1 is present in any cellular compartment. One-Way ANOVA, Tukey’s multiple comparisons test. **E)** Representative images of PDAC tissue from WT/WT, WT/KO and KO/WT mice stained for p-SMAD2 (in brown) and counterstained with hematoxylin. Scale bars: 60 µm. Inserts: 30 µm. The quantification of the percentage of positive nuclei normalized to the average of the WT/WT group is presented in the graph. N = 5 - 6 mice/group. One-Way ANOVA, Tukey’s multiple comparisons test. ns = not significant. **F)** Percentages of CD45^+^, CD45^+^/CD11b^+^ and CD45^+^/CD3^+^ cells out of total live cells in the tumors of WT/WT, WT/KO and KO/WT mice (n = 3 to 4 mice/group). Note the tendency for decreased amounts of these immune cells when NGL-1 is absent in one of the cellular compartments. One-Way ANOVA, Tukey’s multiple comparisons test. All graphs depict mean ± SEM. * p < 0.05. ns: not significant.

Surprisingly, BM chimeras with NGL-1 loss in either immune cells (WT host/KO donor) or stromal cells (KO host/WT donor) did not phenocopy the tumor protection seen in germline NGL-1 KO mice, as assessed by pancreas weight (Fig. 3B). This suggests that NGL-1 expression in either compartment (immune or local stroma) is sufficient for its pro-tumor TME role. Similarly to the intact WT controls, shown in Sup. Fig. 3, these tumors were poorly differentiated and presented similar percentages of cleaved caspase 3, with decreased percentages of positive Ki67 nuclei in the NGL-1 KO host/WT donor group (Sup. Fig. 6C-E). Interestingly, despite having similar tumor burden, the tumors of the mice lacking NGL-1 in either recruited immune cells, or in the local stroma, produced less IL-6 and a trend towards less GM-CSF, but produced the same levels of TGF-β compared to the WT controls (Fig. 3C). This suggests that the pro-tumor effects of NGL-1 might be dependent on TGF-β. Accordingly, different from the mice lacking NGL-1 in all compartments, all chimeric mice presented the same levels of collagen bundling and maturation, as illustrated by the SHG analysis of the tumors (Fig. 3D), and similar levels of p-Smad2 (Fig. 3E). Further, there were no differences between the assorted groups in the percentages of myeloid and T cells, despite the decreased production of IL-6 and GM-CSF by the mice lacking NGL-1 in either the recruited immune cells or in the local stroma (Fig. 3F). These data suggest that a single WT compartment (e.g., BM-derived or local stroma), capable of generating TGF-β, suffices to overcome the antitumor effects observed in germline NGL-1 KO mice.

### Single-cell RNA sequencing of tumors isolated from NGL-1 KO mice reveals decreased expression of TGF-β and AP-1 family members

To further explore how NGL-1 modulates the phenotype of different subsets of cells in the PDAC TME, we performed single-cell RNA sequencing (scRNAseq) in tumors isolated from WT or NGL-1 KO mice. We identified multiple subsets of cells, including endothelial, epithelial, acinar cells, fibroblasts, T cells/NK cells, B cells, and myeloid cells (Sup. Fig. 7A and B, Sup. Table 3). The tumors from NGL-1 KO mice presented equal to relatively decreased abundances of every cell subset identified compared to tumors from WT hosts, except for B cells (Sup. Fig. 7C).

Because of the decreased production of TGF-β and expression of p-Smad2 in the NGL-1 KO tumors, and due to the rescue of these phenotypes by the chimeric mice, we queried the expression of genes involved in the TGF-β signaling. We observed that many of the genes were downregulated in different cell types derived from tumors growing in NGL-1 KO mice, such as *Tgfb1*, *Smad1*, *Smad3*, *Smad4*, *Smad5* and other related genes such as *Thbs1* (Sup. Fig. 7D). This result is in accordance with the observed lower TGF-β and expression levels of p-Smad2 in these tumors compared to tumors from WT mice (Fig. 2C-E). These results further support our previous observations, where a functional NGL-1 compartment in the chimeric mice (e.g., BM or local stroma) could trigger a TGF-β feed forward loop that overcomes NGL-1 deficiency.

We also performed an unbiased analysis using Ingenuity (Qiagen) to uncover the pathways that were enriched in T cells, neutrophils, myeloid cells, fibroblasts, epithelial cells, and B cells from the tumors growing in NGL-1 KO mice (Sup. Fig 7E, Sup. Table 4). The top pathways that were predicted to be activated across all cell types were “RHOGDI signaling”, “antioxidant action of vitamin C” and “PPAR signaling”. Most of the pathways identified were predicted to be inhibited in the different cell types from tumors in NGL-1 KO mice, and encompassed numerous immunomodulatory and cell organization pathways, including “coronavirus pathogenesis pathway”, “G-protein-coupled receptor signaling”, “actin cytoskeleton signaling”, “hepatic fibrosis signaling pathway”, “CXCR4 signaling”, “IL-6 signaling”, and more (Sup. Fig 7E, Sup. Table 4). Because many of the enriched pathways in fibroblasts contained the genes “*Fos*”, “*Jun*”, “*Nfatc1*”, “*Nfatc2*” (Sup. Table 5) and as observed in the word cloud analysis for enriched terms in the canonical pathways represented in fibroblast (Sup. Fig. 7F), we queried the expression of genes pertinent to the AP-1 and NFAT transcription factor families in fibroblast and in other cell subsets (Sup. Fig. 7G-H). Accordingly, we detected downregulation of genes such as *Fos*, *Fosb*, *Fosl1*, *Maf* in different subsets of NGL-1 KO cells (Sup. Fig. 7G), as well as downregulation of *Nfatc1* and *Nfatc2*, when compared to WT cells (Sup. Fig. 7H). Altogether, these results suggest that NGL-1 modulates the expression of key targets related to immunity, ECM organization, and development, prompting us to further investigate the effects of targeting NGL-1 in different subsets of cells, with a focus on myeloid cells, T cells, and fibroblastic populations.

### NGL-1 promotes pro-PDAC phenotypes in stromal and epithelial cells

To take a deeper look into different subsets of cells in the TME, we performed supervised sub-clustering of the myeloid, T cells, fibroblastic, and epithelial cells (Sup. Table 6). The myeloid cell population was subdivided into 4 clusters (Sup. Fig. 8A and B, Sup. Table 6). Cluster 1 (myeloid 1) retained genes corresponding to antigen presentation and dendritic cells, such as *H2-Eb1*, *H2-Ab1*, *H2-Aa*, *Cd74*, *Itgax* and downregulation of classic macrophages markers such as *Chil3*. Interestingly, the tumors of the NGL-1 KO mice presented a higher percentage of these cells compared to WT hosts, suggesting improved capabilities to present antigens to T cells and potentially elicit an anti-tumor immune response. Cluster 2 (myeloid 2) was also enriched in tumors from NGL-1 KO mouse and presented macrophages markers, such as *Chil3*, *Cd14*, *Clec4e*, *Clec4d*. Moreover, this cluster presented upregulation of inflammatory markers, such as *Cxcl2*, *Il1b*, *Ptgs2* and upregulation of *Ccr2*, suggesting that these might be inflammatory monocyte-derived macrophages (56). Cluster 3 (myeloid 3) retained expression of *C1qa*, *C1qb*, *C1qc* and *Mrc1* (mannose receptor c-Type 1), which indicates a tumor-associated macrophage population (57) (Sup. Fig. 8A-C). Finally, cluster 4 (myeloid 4) was enriched in cells expressing markers of immunosuppression, such as *Arg1*, *C1qb*, *Saa3*, *Apoe*, *Trem2* and found to be less represented in the myeloid cell population in tumors from NGL-1 KO mouse (57,58) (Sup. Fig. 8A-C). Additionally, NGL-1 KO myeloid cells downregulated *Arg1*, *C1qa, C1qb, C1qc,* and *Trem2* (Sup. Fig. 8D), further indicating that these cells are less immunosuppressive. These data suggest that NGL-1 could modulate the function of myeloid cells, possibly polarizing them towards a pro-PDAC immunosuppressive phenotype. In turn, loss of NGL-1 could trigger a shift towards an inflammatory/antigen processing and presentation phenotype, which would stimulate the anti-tumor activity of T cells. In fact, one of the top pathways predicted to be upregulated in the NGL-1 KO myeloid cells subset was the “TREM1 pathway”, which is involved in the establishment of an inflammatory response to microbials, eliciting innate immunity (59) (Sup. Fig. 8E, Sup. Table 5). Pathways involved in immunosuppression, such as “Endoplasmic reticulum stress”, “Complement System” and “STAT3 pathway” were predicted to be downregulated, further supporting the notion of a shift in myeloid cell phenotype upon NGL-1 loss (Sup. Fig. 8E, Sup. Table 5).

Since results showed increased percentages of T cells in the tumors from NGL-1 KO mice (Fig. 2G), we next queried RNA expression in these cells. Six T/NK cells subclusters were identified, based on differential gene expression profile (Sup. Fig. 9A-C, Sup. Table 6). The tumor from NGL-1 KO mouse had cells overrepresented in clusters 0 and 1 compared to the tumor from WT mouse. Cluster 0 presented cells upregulating genes related to naïve/resting states, such as *Klf2* and *Lef1*, and commitment to T cell lineage (*Bcl11b),* while markers of activation, such as *Icos, Tnfrsf4* (encoding OX40) and *Tnfrsf9* (encoding 4-1BB) were downregulated (60) (61,62) (Sup. Fig. 9B). Cluster 1 seemed to comprise naïve/resting CD8 T cells, with upregulation of *Cd8b*, *Cd8a*, *Ccr7*, *Sell*, *Lef1* (62,63 (Sup. Fig. 9A-D) (Sup. Fig. 9A-D). These suggest that NGL-1 KO tumors present T cells that are naïve-like or stem cell-like, as indicated in the feature and violin plots (Sup. Fig. 9D and F). Cells in Cluster 2 upregulated *Fcer1g*, *Tyrobp*, *Id2* and *Tbx21*, a signature resembling innate lymphoid cells, NK cells and/or NKT cells (64,65)(Sup. Fig. 9A-C). These cells also expressed markers of activation and exhaustion, such as *Gzma*, *Gzmb*, *Gzmc*, *Xcl1*, *Tgfb1* (*66,67*). Among the most upregulated genes in cluster 3 were *Foxp3, Ctla4 and Il2ra*, indicating that this is likely the Treg cluster (62) (Sup. Fig. 9A-C). Cluster 4 involved cells expressing *Cd4*, *Cblb*, *Adam17, Camk2d* (Sup. Fig. 9A-C), suggestive of engagement and modulation of TCR signaling in CD4 T cells, (68,69) (Sup. Fig. 9C). Cluster 5 included cells expressing genes commonly upregulated by γδT cells, such as *Tcrg-C1*, *Trdc*, and *Tcrg-C2* (Sup. Fig. 9C) (70–72). Notably, tumors in WT mice showcased T cell signatures of immune evasion, denoted by cells in clusters 3 and 5, underrepresented in NGL-1 KO host (Sup. Fig. 9A and E), indicative of better PDAC patient outcomes (73). In accordance, T cells from tumors in NGL-1 KO mouse upregulated *Cd8a* (upregulated in CD8 T cells), *Lef1*, *Sell*, *Tcf7* and other markers typical of naïve T cells, downregulating *Ctla4* and *Icos* (T cell activation/exhaustion) (Sup. Fig. 9F). Pathways related to modulation of innate immunity were predicted to be activated in the T cells from the tumors in NGL-1 KO mice (Sup. Fig. 9G, Sup. Table 5), while pathways modulating IL-17 signaling, inflammatory cell death, and general T cell activation were predicted to be inactivated in this tissue (Sup. Fig. 9G, Sup. Table 5).

Fibroblasts, one of the most abundant cell types in human pancreatic cancer, also displayed significant shifts in gene signature profiles. We subclustered the fibroblastic population, identifying four populations based on distinct gene profiles (Sup. Fig. 10A and B, Sup. Table 6). Cluster 1 resembled mesothelial-like cells, based on the expression of *Slpi*, *Crip1*, *Upk3b*, *Msln* and others (74) (Sup. Fig. 10A and B). Clusters 2 and 4 expressed a mixture of genes indicative of ECM remodeling and immunomodulation, such as *Apod*, *Ccl11*, *Col14a1*, and *Ifngr1* (cluster 2), as well as *Col1a1*, *Col3a1*, *Mmp2*, *C4b*, *Ly6c1*, *Col1a2*, *Ly6a*, *Pi16*, *Cxcl12* (cluster 4) (Sup. Fig 10A and B). Cluster 3 was enriched in genes related to immunomodulation and antigen presentation, such as *Cxcl13*, *Saa3*, *Cxcl1*, *Cxcl10*, *Clu*, *H2-Q7*, *Nfkbia*, *H2-Eb1*, and *H2-Ab1* (75)(Sup. Fig 10A and B). A decrease in the percentages of cells in cluster 1 and 4 was noted in tumors from NGL-1 KO mice, compared to WT. Further, a higher percentage of cells from cluster 3 was noted in tumors from KO mice (Sup. Fig. 10C). In fact, we observed upregulation of genes involved in antigen presentation, such as *H2-Q7, H2-Eb1, H2-Ab1, CD74* in NGL-1 KO CAFs (Sup. Fig. 10D). This indicates that in the tumors of NGL-1 KO mice, fibroblasts could foster a TME that is increasingly conducive to anti-tumor immunity. Accordingly, there was downregulation of genes involved in pro-tumoral immunosuppression, such as *C4b*, and members of the TGF-β family, such as *Tgfb1*, *Smad1*, *Smad3*, *Smad4*, *Thbs1*, and *Maf* (Sup. Fig. 10E). Of note, the expression of genes such as *Col1a1*, *Col1a2*, *Col3a1,* and the CAF marker podoplanin (*Pdnp*), remained unchanged (Sup. Fig. 10F), suggesting that fibroblasts prevail in both experimental models. Pathway analyses indicated that “oxidative phosphorylation”, “VDR/RXR Activation”, “Inhibition of Matrix Metalloproteases”, and “Interferon Signaling”, were upregulated by NGL-1 KO CAFs, while “IL-6 Signaling”, “Senescence Pathway”, “Integrin Signaling”, and “Pancreatic Adenocarcinoma Signaling” were predicted to be downregulated, with these being involved in pro-tumoral CAF activity (Sup. Fig. 10G, Sup. Table 5).

Importantly, NGL-1 KO stroma modulated the epithelial cell transcriptome of KPC3 cells injected in the pancreas, suggesting a broader stroma-driven rewiring of cancer cell function. For instance, sub-clustering of the epithelial compartment revealed 5 different clusters, from which cluster 2 “Epi_proliferating” contains proliferating cells expressing *Ube2c*, *Top2a*, and *Tuba1b*, while cluster 3 is mainly composed by cells with a mesenchymal phenotype, expressing *Fn*, *Ccn1*, and *Ccn2* (Sup. Fig. 11A and C, Sup. Table 6). The percentage of epithelial cells with both proliferating and mesenchymal features was lower in tumors collected from NGL-1 KO mice compared to WT (Sup. Fig. 11A and B). Pathways such as “SPINK1 pancreatic cancer pathway” and “unfolded protein response”, related to pancreatic secretion/function, were predicted to be upregulated in the tumors from NGL-1 KO mice (Sup. Fig. 11D, Sup. Table 5). On the other hand, pathways related to restoring apical junctions, regulation of actin cytoskeleton, and cell motility regulated by Rho GTPases, such as “regulation of actin-based motility by Rho” and “RHOA signaling” were predicted to be downregulated in NGL-1 KO mice (Sup. Fig. 11D, Sup. Table 5). These results support the notion that limiting NGL-1 in the TME can not only restrict tumor size but also modulate the functional traits of tumoral cells.

Collectively, the scRNAseq analysis comparing tumors growing in WT and NGL-1 KO hosts suggested that NGL-1 could be involved in the modulation of the phenotype and activity of different subsets of cells in the pancreatic TME, further supporting the hypothesis that stromal NGL-1 plays a pro-tumor role in pancreatic cancer tumorigenesis.

### Ablation of NGL-1 in CAFs results in downregulation of genes involved in myofibroblastic and immunomodulatory functions

The *in vivo* data, together with the scRNAseq analysis, suggested that NGL-1 controls important pro-tumor features in CAFs. To further evaluate the effects of NGL-1 in CAFs, we knocked down NGL-1 in immortalized CAFs previously isolated from PDAC patients by our group (29) using the CRISPRi system (Fig. 4A). Given the known importance of maintaining the CAF/ECM unit for *ex vivo* functional studies with fibroblasts (76), we cultured control (CON), NGL-1 KD1, and NGL-1 KD2 CAFs within their own ECM, as CAF/ECM units, using this system for all the functional studies conducted *in vitro* (29,77,78). To explore NGL-1 dependent global changes in gene expression, we ran bulk RNA sequencing (RNAseq) comparing CON and NGL-1 KD CAF units. The knockdown of NGL-1 led to unique differentially expressed genes (Fig. 4B-C, Sup. Table 7). Limiting NGL-1 expression in human CAFs phenocopied NGL-1 KO mice in that its loss diminished the expression of known pro-tumor CAF genes (e.g., *THBS1, CTHRC1, SPP1, LRRC15, IL11*) (Fig. 4B). Also, in accordance with the scRNAseq data, we identified the downregulation of *FOSL1* (gene that codes for FRA-1, a member of the AP-1 family) and *NFATC2* (gene that codes for NFAT1), further confirming a relationship between NGL-1 and pathways involved with the establishment of the immune response (Fig. 4B). As expected, the expression of genes involved in ECM deposition and remodeling, such as *COL1A2* and *ACTA2* (encoding α-SMA) were either upregulated or maintained in the KD CAFs, suggesting that targeting NGL-1 does not limit ECM production, but rather the function of the CAF/ECM units (Fig. 4B). Limiting NGL-1 expression was also accompanied by the downregulation of *FIBIN* (Fig. 4B), a growth factor expressed across different vertebrates’ embryonic tissues and highly expressed in adult cerebellum and skeletal muscle (79). This gene is involved in limb development and potentially interacts with several transcription factors of known importance in cancers, including pancreatic cancer, such as glucocorticoid receptors, cAMP responsive element protein (CREB), NF-κB, and the WNT-activated LEF/TCF transcriptional factors (79). Interestingly, the immune checkpoint *CD274* (PD-L1) was also downregulated in NGL-1 KD CAFs, implying a relationship between NGL-1 and immune inhibitory mechanisms in the TME (Fig. 4B). Altogether, these results indicate that NGL-1 is involved in different aspects of CAF functionality including both myofibroblastic and immunomodulatory features. In support of this, we noted that many of the pathways predicted to be upregulated and downregulated by NGL-1 KD CAFs were related to immunity, ECM modulation, and metabolism, all key hallmarks of human CAF biology (Fig. 4D, Sup. Table 8).

**Figure 4:**
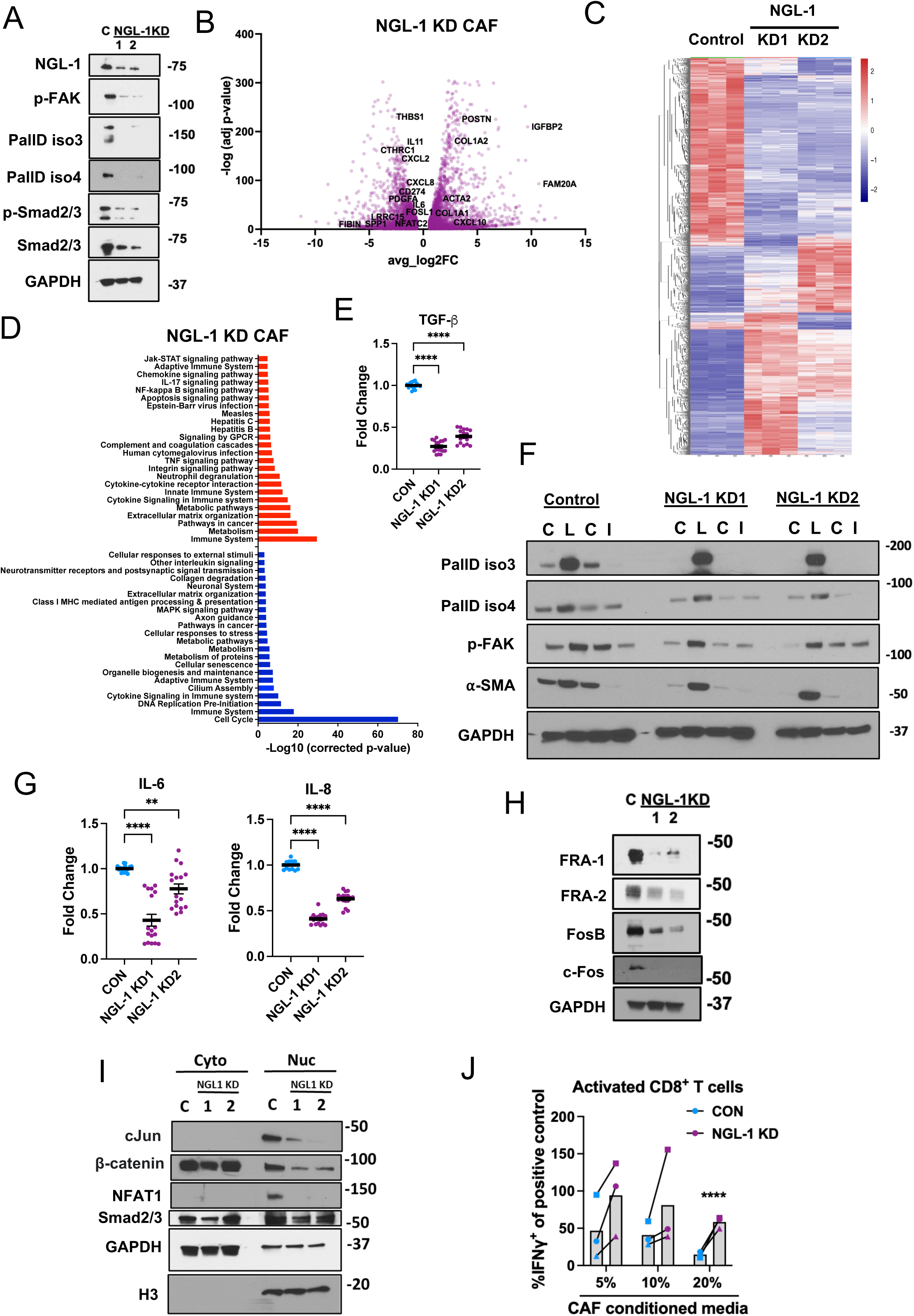
NGL-1 controls pro-tumor features of CAFs, including immunosuppression, in a TGF-β dependent manner. **A)** Representative western blotting for NGL-1 and the myofibroblastic markers p-FAK, PallD iso 3, PallD iso 4, p-Smad2/3 and Smad2/3 in Control (C), NGL-1 KD 1 and 2 CAFs, showing decreased expression of these markers in the NGL-1 KD CAFs compared to controls. GAPDH was used as loading control. N = 5. **B)** Control, NGL-1 KD1 and NGL-1 KD2 CAFs were subjected to RNAseq analysis, in triplicate. Volcano plot of the genes that are significantly upregulated or downregulated upon knockdown of NGL-1 in CAFs, identified by bulk RNAseq. Data are plotted as -log (adjusted p-value) against log2 (fold change). **C)** Heatmap of the differentially expressed genes (FDR5, fold change 1.5) between Control, NGL-1 KD1 and NGL-1 KD2 CAFs, in triplicate, with upregulated genes in red and downregulated genes in blue. Note the difference in the transcriptional profile of CAFs lacking NGL-1 compared to Control CAFs. **D)** Pathways predicted to be upregulated (red) and downregulated (blue) in the NGL-1 KD CAFs, according to the differentially expressed genes (-Log10, corrected p-value). Pathways were filtered with p < 0.01. **E)** Quantification of ELISA for TGF-β in the conditioned media of Control (CON), NGL-1 KD1 and NGL-1 KD2 CAFs growing in 3D. N = 3 independent experiments, with 5-6 replicates of each sample/experiment. The cytokine levels are presented as fold change compared to the Control group, which was set as 1. One-Way ANOVA, Dunnett’s multiple comparisons test, compared to Control. **F)** Representative western blotting for the myofibroblastic markers PallD iso 3, PallD iso 4, p-FAK and ⍺-SMA, in Control, NGL-1 KD1 and NGL-1 KD2 CAFs treated with 10 ng/mL recombinant TGF-β (ligand, L) and 10 µM TGF-β receptor inhibitor (inhibitor, I), and their respective vehicles (control, C, citric acid and DMSO, respectively). We observed that the NGL-1 KD CAFs retain the ability to respond to TGF-β treatment. GAPDH was used as loading control. N = 3. **G)** ELISA showing the decreased production of the immunosuppressive cytokines IL-6 and IL-8 by the NGL-1 KD CAFs compared to CON CAFs. The cytokines were detected in the conditioned media of CON, NGL-1 KD1 and NGL-1 KD2 CAFs growing in 3D. N = 3 independent experiments, with 5-6 replicates of each sample/experiment. The cytokine levels are presented as fold change compared to the Control group, which was set as 1. One-Way ANOVA, Dunnett’s multiple comparisons test, compared to Control. **H)** Representative western blotting for the AP-1 transcription factor members FRA-1, FRA-2, FosB and c-Fos in Control (C), NGL-1 KD 1 and 2 CAFs, displaying decreased expression of these markers in the NGL-1 KD CAFs compared to CON CAFs. GAPDH was used as loading control. N = 3. **I)** Representative western blotting of the cytoplasmic (Cyto) and nuclear (Nuc) fractions of Control (C), NGL-1 KD 1 and 2 CAFs, for c-Jun, β-catenin, NFAT1 and Smad2/3, showing downregulation of these transcription factors in the nuclear fractions of NGL-1 KD CAFs compared to control CAFs. The enrichment of the fractions was determined by the expression of GAPDH (for cytoplasmic fraction) and histone 3 (H3, for nuclear fraction) in each fraction. N = 3. **J)** CD8^+^ T cells were isolated from PBMCs from healthy donors and stimulated *in vitro* (2×10^5^ cells/well) with DynaBeads Human T-activator and IL-2 in the presence or absence of 5, 10 and 20% of conditioned media from CON (blue) or NGL-1 KD1 CAFs (purple). T cell activation was assessed by flow cytometry, showing the percentage of IFNγ^+^ T cells in relation to the positive control (stimulated CD8^+^ T cells, no conditioned media). The graphs show the effect of the different conditioned media, at different concentrations, in each donor. Note that the conditioned media of NGL-1 KD1 CAFs is less immunosuppressive, with CD8^+^ T cells becoming activated (increased percentage of cells expressing IFNγ), compared to Control CAFs at the same % CM. N = 3 independent experiments, each experiment conducted with cells from a different donor. Multiple unpaired t tests, compared to CON CAF CM. All graphs depict mean ± SEM. ** p < 0.01, **** p < 0.0001.

### NGL-1 modulates myofibroblastic features of CAFs, in a TGF-β-dependent manner

To evaluate the effects of NGL-1 in CAF functionality, we first characterized the phenotype of the ECM produced by the assorted CAF/ECM units. Limiting NGL-1 expression did not affect ECM deposition (measured by its thickness) or its topographical alignment (measured by fibronectin fiber orientation) (Sup. Fig. 12A). These results agreed with the scRNAseq data showing similar expressions of *Col1a2* and *Col3a1* between WT and NGL-1 KO fibroblasts, and the bulk RNAseq data where NGL-1 KD CAFs upregulated the expression of these collagens (Sup. Fig. 10F and Fig. 4B). Interestingly, NGL-1 KD CAFs displayed increased lipid droplet retention compared to CON CAFs, which constitutes a well-known feature of normal resident fibroblasts in the pancreas (80–84) (Sup. Fig. 12B). Results indicate that NGL-1 in CAFs shapes the pro-tumor ECM niche; while no major ECM fiber orientation changes were noted, the pro-tumoral ECM functionality was altered by NGL-1 loss. Notably, PDAC cells cultured under nutrient deprivation within NGL-1 KD CAF-derived ECM presented reduced survival compared to PDAC cells cultured within CON CAF ECM (Sup. Fig. 12C). This effect was maintained even when PDAC tumor cells were co-cultured with CON CAFs within NGL-1 KD CAF ECM, suggesting that limited NGL-1 in CAFs alone can compromise the pro-tumor function of the CAF-derived ECM. Consequently, these findings point to the ECM as a pro-survival cue for PDAC, and that NGL-1 in CAFs modulates the units’ tumor supportive role (Sup. Fig. 12C). The *in vivo* data supports this, as NGL-1 KO tumors lack mature collagen bundles (Fig. 2B).

To further understand these differences, we determined the expression of proteins known to be involved in CAF/ECM units’ function in support to PDAC survival, such as p-FAK and the actin-bundling protein palladin (PallD, isoforms 3 and 4), with the expression of these proteins being inversely correlated with PDAC patient overall survival (12,27,85–87). Interestingly, there was a downregulation of these and other myofibroblastic markers upon KD of NGL-1 in CAFs (Fig.4A). Based on the results observed in the chimeric mice, and because TGF-β is a key modulator of general (88) and PDAC-associated fibroblast functions (12,25,89), we hypothesized that NGL-1 controls TGF-β signaling. To this end, we determined the expression levels of p-Smad2/3 in CON vs. NGL-1 KD CAFs. As expected, there was downregulation of p-Smad2/3 by the NGL-1 KD CAFs, which together with the decreased levels of PallD iso 3 and 4, suggest that NGL-1 regulates pro-tumoral CAF activity via modulation of canonical TGF-β signaling (Fig. 4A). These results are in accordance with the overall decreased levels of p-Smad2 in total tissue and in the TME of the tumors from NGL-1 KO mice (Fig. 2D-E), and with the downregulation of members of the TGF-β signaling pathway across different cell types, including fibroblasts, observed in the scRNAseq analysis (Sup. Fig. 7D). We then asked whether NGL-1 deficiency also disrupts the TGF-β signaling by limiting its own secretion. As expected, NGL-1 KD CAFs produced substantially less TGF-β than CON CAFs (Fig. 4E), which might account for the differences in expression of other pro-tumor CAF markers (12,27), and for decreased tumor burden in the NGL-1 KO mice (Fig. 2A, Sup. Fig. 3C).

In the chimeric mice, restoring NGL-1 expression in the local stroma or in the bone marrow, recovered production of TGF-β, the pattern of mature collagen, and the expression of p-Smad2, resulting in no differences in tumor burden between groups. These results led us to the premise that not only is NGL-1 expression correlated with the ability of cells to secrete TGF-β, but that NGL-1 deficient cells retain the ability to respond to exogenous TGF-β. To test these two hypotheses, we treated CON and NGL-1 KD CAFs with recombinant TGF-β1 (here called Ligand, or Lig), TGFBR1 inhibitor (SB431542, Inh) and their respective vehicles (C1 for citric acid and C2 for DMSO) during ECM production and tested the expression of selected CAF/ECM unit markers, as in Fig. 4A. As expected, CON CAFs treated with TGF-β Lig upregulated the expression of p-FAK, PallD iso 3 and 4, and α-SMA, while these same markers were downregulated in response to TGFBR1 Inh (Fig. 4F), in a manner similar to NGL-1 KD CAFs. Moreover, as hypothesized, the expression of these myofibroblastic markers were restored in response to TGF-β Lig. These results suggested that NGL-1 deficient CAFs retain the ability to respond to TGF-β (Fig. 4F). Importantly, these observations were confirmed in a second immortalized CAF line (CAF line #2), whereby KD of NGL-1 also resulted in the normalization of CAFs, illustrated by increased lipid droplets (Sup. Fig. 12D), reduced TGF-β secretion (Sup. Fig. 12E), and decreased expression of PallD iso 3 and 4, α-SMA, p-Smad2/3, and p-FAK, with a similar response to TGF-β Lig (Sup. Fig. 12F). Collectively, these data suggest that availability of TGF-β in the TME could overcome the downregulation of NGL-1, but if NGL-1 is inhibited in different cell types (i.e., systemically), the production of TGF-β is globally affected, and its availability is drastically decreased, compensating for this non-cell autonomous mechanism. Supporting this premise, we noted that treatment of CON CAFs with TGF-β Lig led to the upregulation of NGL-1, demonstrating a co-regulatory signaling loop between NGL-1 and TGF-β (Sup. Fig. 12F).

### Loss of NGL-1 in CAFs reduces pro-tumor cytokine production

Another important feature of CAFs in PDAC is immunosuppression. It is known that CAFs produce a plethora of pro-tumor cytokines involved in the attraction and polarization of myeloid cells, Tregs and inhibition of cytotoxic cells such as T and NK cells (29,90–93). In fact, TGF-β is known for its effects on Treg polarization and conversion of macrophages into a tumor promoting phenotype (94–96). Therefore, we investigated if the knockdown of NGL-1 in CAFs affected their secretion of pro-tumor cytokines such as IL-6 and IL-8, which have important roles in the maintenance of immunosuppression in PDAC. As expected, NGL-1 KD CAFs produced significantly less IL-6 and IL-8 than CON CAFs, suggesting that NGL-1 also modulates pathways controlling immune response in CAFs (Fig. 4G, Sup. Fig. 13A).

In the scRNAseq analysis we noted that tumors in NGL-1 KO mice downregulated the expression of several members of the AP-1 transcription factor family, including *Fosl1* (encoding FRA-1), *Fosb* and *Fos*, as well as *FOSL1* (observed in the bulk RNAseq analysis). Accordingly, we confirmed that NGL-1 KD CAFs express decreased protein levels of FRA-1, FRA-2, FosB and c-Fos compared to CON CAFs (Fig. 4H, Sup. Fig. 12F). Furthermore, we found that the AP-1 transcription factor c-Jun is also downregulated in the nuclear fraction of NGL-1 KD CAF lysates compared to CON CAFs (Fig. 4I). Similarly, expression of p-Smad2/3 was decreased in the nuclear fraction of NGL-1 KD CAF lysates compared to CON CAFs (Fig. 4I). Based on this profile, we assessed the expression of β-catenin, a transcription factor known to bind to AP-1 transcription factors and co-regulate gene expression (97–99). We found that the nuclear expression of β-catenin (representing its active form) was indeed downregulated in the NGL-1 KD CAFs compared to CON CAFs, suggesting that the canonical WNT signaling is modulated downstream to NGL-1 as well (Fig 4I).

Since expression of NFAT genes were also downregulated in the tumors from NGL-1 KO mice, as well as in the human NGL-1 KD CAFs, we determined the protein expression of NFAT1 in CON and NGL-1 KD CAFs. Excitingly, NGL-1 KD CAFs expressed less NFAT1 compared to CON CAFs (Fig. 4I, Sup. Fig. 12F). Classically, NFAT is a family of transcription factors identified in activated T cells in response to phospholipase C signaling, Ca^2+^ influx to the cytoplasm and the activation of calcineurin by calmodulin (100). NFAT translocates to the nucleus and binds to assorted transcription factors, among these are members of the AP-1 family, which modulate cytokine secretion (101). These results explain the phenotype observed when targeting NGL-1, with decreased secretion of pro-tumor cytokines by CAFs.

We next questioned the effects of limiting NGL-1 expression in CAFs on the secretion of multiple immunomodulatory cytokines. The secretion of numerous pro-tumor cytokines and chemokines were decreased, such as GM-CSF, G-CSF, IL-1α, IL-1β, IL-15, CXCL10 and CCL20 in these CAF units compared to CON CAFs, as well as VEGF and FGF2, demonstrating a key modulatory role for NGL-1 (Sup. Fig. 13B). To evaluate the functional impact of this phenotype, we cultured naïve CD8^+^ T cells isolated from healthy human peripheral blood mononuclear cells (PBMCs) together with conditioned media (CM) collected from either CON or NGL-1 KD1 CAFs. Results showed that the CD8^+^ T cells cultured with increased percentages of the CM produced by NGL-1 KD CAFs (5, 10 or 20%) were more active, expressing higher IFNγ, compared to CM produced by CON CAFs (Fig. 4J). These results further support the notion that targeting NGL-1 in CAFs restricts the immunosuppressive TME, by boosting the capacity of cytotoxic T cell to secrete this inflammatory cytokine (Fig. 4J). Altogether, these results suggest that NGL-1 promotes the immunosuppressive phenotype in CAFs, through the modulation of TGF-β, and numerous other key immunomodulatory factors, such as members of the AP-1/NFAT transcription factor families and cytokines, which are known to co-regulate the WNT and TGF-β signaling pathways (102).

### NFAT1 partially mediates NGL-1 dependent signaling and function in CAFs

Because the expression of NFAT1 was significantly decreased in NGL-1 KD CAFs, we hypothesized that the pro-tumor activities driven by NGL-1 are dependent on NFAT1 expression. To test this premise, we queried whether knocking down NFAT1 (*NFATC2*) in CAFs phenocopies the loss of NGL-1 (Fig. 5). Strikingly, NFAT1 KD CAFs presented decreased expression of p-Smad2/3 compared to CON CAFs, suggesting reduced canonical TGF-β signaling, but these cells still effectively responded to modulation of TGF-β (TGFBR1 Inh and/or TGF-β Lig) (Fig. 5A). Further, TGF-β Lig led to the upregulation of NFAT1 in CON CAFs, suggesting that not only NFAT1 regulates canonical TGF-β signaling but that increased TGF-β signaling results in increased NFAT1 expression (Fig. 5A). Interestingly, while TGF-β signaling was modulated, the profile of myofibroblastic markers did not phenocopy that of NGL-1 KD CAFs; levels of α-SMA in NFAT1 KD CAFs were higher compared to CON CAFs, while levels of PallD iso 3 but not iso 4 were decreased (Fig. 5A). Similarly to NGL-1 KD CAFs, all these markers were significantly upregulated in response to TGF-β Lig, reinforcing that, similarly to NGL-1 loss, NFAT1 KD CAFs retain the ability to respond to this factor (Fig. 5A). Notably, levels of NGL-1 were decreased in the NFAT1 KD CAFs, yet treatment with TGF-β Lig was no longer sufficient to rescue the NGL-1 expression (Fig. 5A). These results suggest that upregulation of NGL-1 by TGF-β is dependent on NFAT1 expression. Despite the decreased levels of NGL-1 in NFAT1 KD CAFs, the AP-1 factor FosB was upregulated, while levels of FRA-1 remained low (Fig. 5A). Moreover, NFAT1 KD CAFs retain more lipid droplets than CON CAFs, which is suggestive of a functional normalization of CAFs (Fig. 5B). Also, NFAT1 KD CAFs produced decreased levels of both TGF-β and GM-CSF, yet the levels of IL-6 and IL-8 were higher compared to CON CAFs (Fig. 5C). These results suggest that while limiting NFAT1 expression shifts CAFs to a more normalized state, it does not completely phenocopy the consequences of losing NGL-1 in these cells. These discrepancies might be explained by the noted high levels of FosB in the NFAT1 KD CAFs.

**Figure 5:**
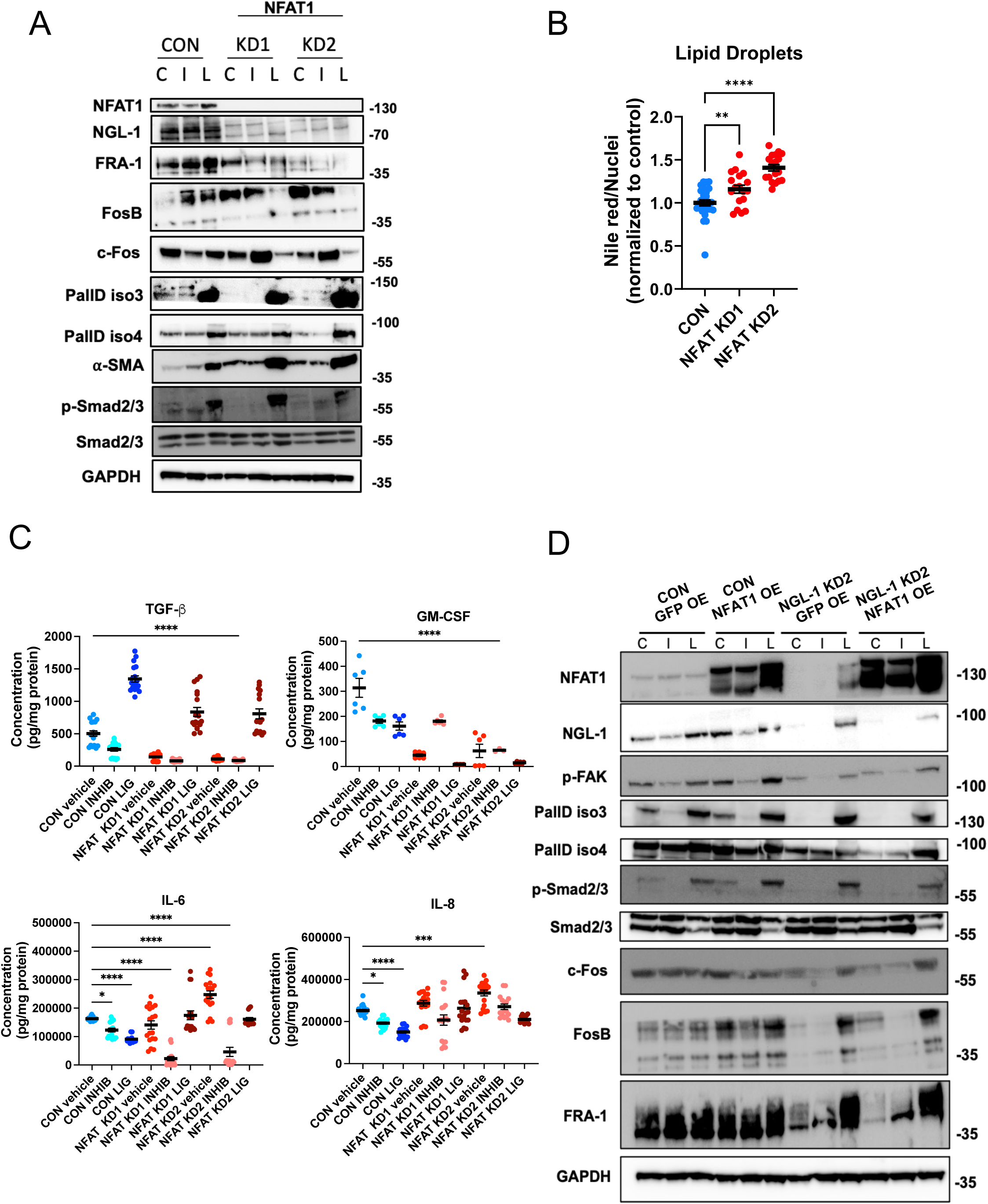
Loss of NFAT1 in CAFs partially phenocopies NGL-1 knockdown. **A)** Representative western blotting for myofibroblastic markers (PallD iso 3, PallD iso 4, p-Smad2/3, Smad2/3 and ⍺-SMA), NGL-1, NFAT1, AP-1 transcription factor members c-Fos, FosB and FRA-1, in Control, NFAT KD1 and NFAT KD2 CAFs treated with recombinant TGF-β (ligand, L) and TGF-β receptor inhibitor (inhibitor, I), and their respective vehicles (control, C, citric acid + DMSO). GAPDH was used as loading control. N = 5. **B)** Quantification of Nile Red intensity, normalized to DAPI^+^ nuclei, in the NFAT1 KD CAFs compared to Control (CON). All values were normalized to the levels of the CON CAFs, set as 1. Each experiment was repeated two times, at least 9 wells/cell line/experiment. One-Way ANOVA, Dunnett’s multiple comparisons test, ** p < 0.01, **** p < 0.0001. **C)** ELISAs for the detection of the different cytokines in the conditioned media of Control (CON), NFAT KD1 and NFAT KD2 CAFs growing in 3D, in the presence of vehicle, recombinant TGF-β or TGF-β receptor inhibitor. Note that NFAT KD CAFs present decreased TGF-β and GM-CSF production compared to Control CAFs, but increased IL-6 and IL-8 levels. N = 2 to 3 independent experiments, with 3 replicates of each sample/experiment. One-Way ANOVA, Dunnett’s multiple comparisons test. * p < 0.05, **** p < 0.0001, compared to Control treated with vehicle. **D)** Representative western blotting of Control or NGL-1 KD2 CAFs overexpressing (OE) either GFP as control, or NFAT1. Membranes were probed for myofibroblastic markers (PallD iso 3, PallD iso 4, p-Smad2/3, Smad2/3 and p-FAK), NGL-1, NFAT1, AP-1 transcription factor members c-Fos, FosB and FRA-1. All the cells were treated with recombinant TGF-β (ligand, L) and TGF-β receptor inhibitor (inhibitor, I), and their respective vehicles (control, C, citric acid + DMSO). GAPDH was used as loading control. N = 3.

To further explore the mechanistic interplay between NGL-1 and NFAT1, we overexpressed (OE) NFAT1 in CON or NGL-1 KD CAFs, while overexpression of green fluorescent protein (GFP) served as an experimental control (Fig. 5D). Results demonstrated that NFAT1 OE in CON CAFs enhanced NGL-1 expression, and this was decreased by TGFBR1 Inh with no further increases in response to TGF-β Lig. Also, while NFAT OE did not suffice to restore p-FAK, PallD iso 3 and 4, or p-Smad2/3 in NGL-1 KD CAFs, treatment with TGF-β Lig in the NFAT1 OE NGL-1 KD CAFs did (Fig. 5D). Interestingly, NFAT1 OE in CON CAFs was able to increase p-FAK but not PallDs or p-Smad2/3. Further, NFAT1 OE in CON CAFs sufficed to upregulate FosB but not c-Fos or FRA-1. TGFR1 Inh reduced FosB expression to CON CAF levels, while TGF-β Lig failed to increase these further. In accordance, NFAT1 OE in NGL-1 KD CAFs also only slightly restored FosB expression. Treating these CAFs with TGFBR1 Inh lowered c-Fos and FosB expression respectively, with TGF-β Lig resulting in increased c-Fos and FosB. Lastly, TGF-β Lig still induced a significant upregulation of FRA-1 in NFAT1 OE NGL-1 KD CAFs (Fig. 5D). In summary, we noted that the expression of PallD iso 3 and 4, p-Smad2/3, Smad 2/3, c-Fos and FRA-1 are dependent on NGL-1, yet independent of NFAT1 OE, since further increasing NFAT1 expression did not affect the levels of those proteins, but as expected, all responded to TGF-β (Fig. 5D).

Collectively, these results demonstrated that NFAT1 regulated certain, but not all, aspects of the NGL-1 driven phenotype in CAFs. Moreover, the loss of NFAT1 had a dramatic effect on the CAF phenotype, while overexpression of NFAT1 was insufficient to rescue NGL-1 loss. This signifies that a baseline expression level of NFAT1 in CON CAFs was necessary to mediate signaling, yet increased expression had little effect.

### Ablation of NGL-1 in immune cells shifts their polarization to an anti-tumor phenotype

To assess the intrinsic roles of NGL-1 in immune cells, we evaluated the proliferative profile of naïve CD8^+^ and CD4^+^ T cells from non-tumor bearing WT and NGL-KO mice after stimulation *in vitro* with anti-CD3 and anti-CD28 (Fig. 6A, Sup. Fig. 14A). Interestingly, stimulated NGL-1 KO CD8^+^ T cells proliferated 30% more than stimulated WT CD8^+^ T cells (Fig. 6B). Similarly, stimulated NGL-1 KO CD4^+^ T cells proliferated 110% more than stimulated WT CD4^+^ T cells (Fig. 6C), suggesting that NGL-1 expression can suppress T cell proliferation/activation. In accordance, single cell secretome analysis on the Isoplexis/Bruker platform (103) showed that stimulated NGL-1 KO T cells cluster distinctly from the WT T cells (Fig. 6D), indicating functional differences when NGL-1 is not expressed. Indeed, compared to stimulated WT T cells (CD3^+^ T cells), the NGL-1 KO T cells presented higher percentage of polyfunctionality (defined by the percentage of cells secreting two or more cytokines (103)) (Fig. 6E), and higher polyfunctionality strength index; a measure of the intensity of cytokine expression by polyfunctional cells (104) (Fig. 6F). These analyses showed that when stimulated, the NGL-1 KO T cells expressed cytokines related to effector, inflammatory and chemoattractant functions (Fig. 6F). Altogether, these results suggest that the absence of NGL-1 in T cells facilitates T cell activation, which may translate into restoring anti-tumor immune responses by blocking NGL-1 in these important TME cells.

**Figure 6:**
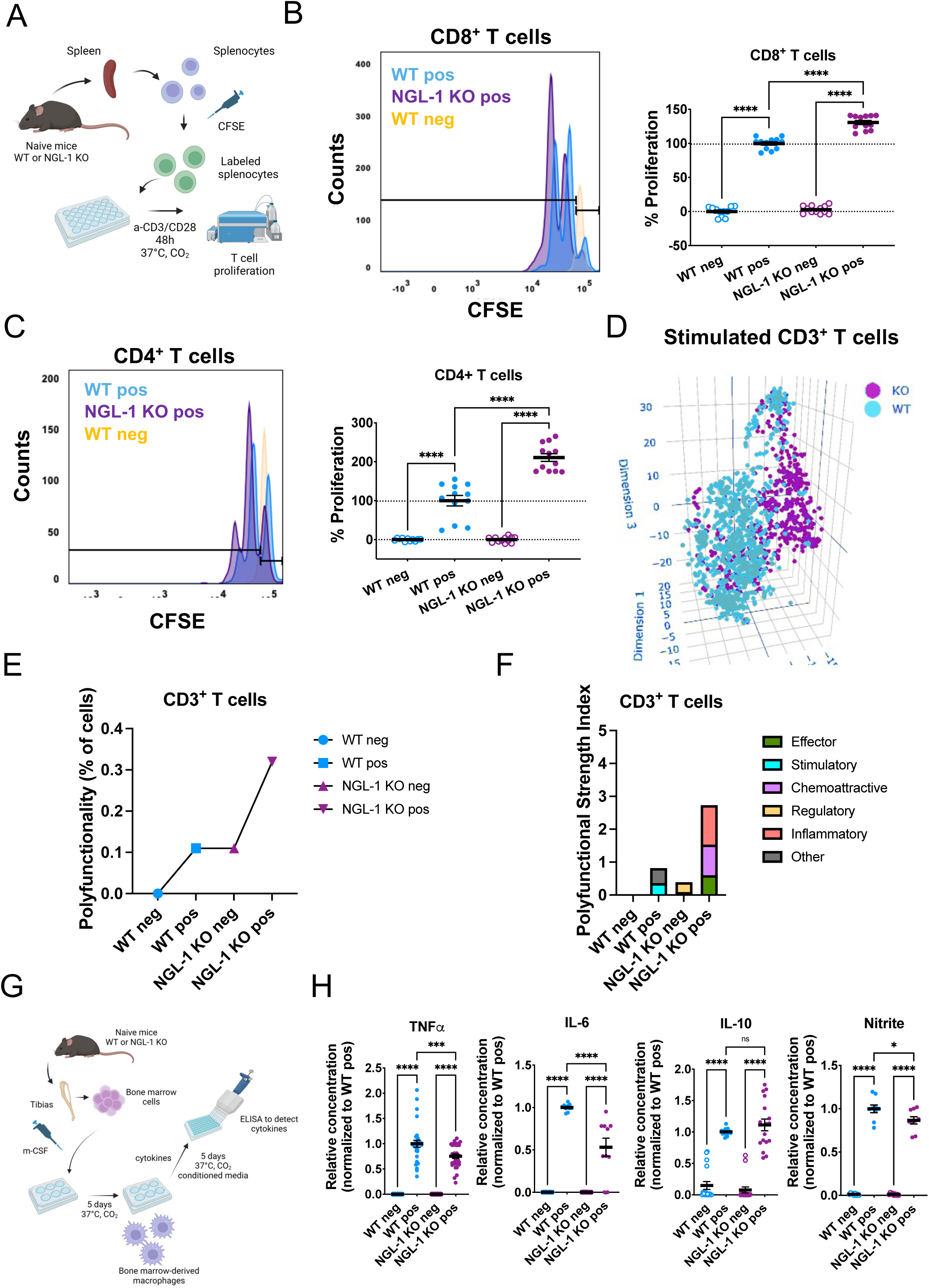
NGL-1 modulates the functionality of T cells and macrophages, serving as a break for T cell proliferation and functionality. **A)** Scheme depicting the *in vitro* stimulation of total splenocytes isolated from WT or NGL-1 KO spleens (naïve mice). CFSE-labeled cells were cultured in the presence (positive, pos) or absence (negative, neg) of anti-CD3 and anti-CD28 antibodies, for 48 hours, then CFSE dilution in CD8^+^ and CD4^+^ T cells was measured by flow cytometry as a readout of cell proliferation (more dilution means more proliferation). Figure made with BioRender.com. **B)** and **C)** Representative flow cytometry plots of the CFSE dilution in **B)** CD8^+^ T cells and **C)** CD4^+^ T cells of the different groups (WT neg, WT pos, NGL-1 KO pos). NGL-1 KO CD8^+^ and CD4^+^ T cells proliferate more than the respective WT cells, as quantified in the graph (mean ± SEM) showing the percentage of proliferation, where 0% represents the normalized proliferation of the WT unstimulated cells, and 100% represents the normalized proliferation of the stimulated WT cells. N = 3 different experiments conducted in triplicate. One-Way ANOVA, Tukey’s multiple comparisons test, **** p < 0.0001. **D-F)** CD3^+^ T cells were also isolated from the WT and NGL-1 KO spleens (naïve mice) by negative selection and stimulated *in vitro* with the same cocktail for the evaluation of polyfunctional index using IsoSpeak software (Bruker), measuring protein secretion at the single-cell level. **D)** t-SNE 3D mapping generated with IsoSpeak software (Bruker) of single-cell clustering based on functional cytokine data and polyfunctional subsets of WT and NGL-1 KO CD3^+^ T cells stimulated *in vitro*. Stimulated WT and NGL-1 KO T cells cluster separately, suggesting differences in phenotype. N = ∼900 cells were analyzed. **E)** Polyfunctional Strength Index (PSI) of CD3^+^ T cells from WT and NGL-1 KO mice, unstimulated or stimulated. This is a measurement of the percentage of polyfunctional single cells secreting multiple cytokines and at higher intensities. The NGL-1 KO T cells present higher PSI compared to the WT T cells in both conditions, unstimulated and stimulated, suggesting higher functionality when lacking NGL-1. **F)** Polyfunctional Strength Index (PSI) of CD3^+^ T cells from WT and NGL-1 KO mice, unstimulated or stimulated, and distribution of the functional categories: Effector, Stimulatory, Chemoattractive, Regulatory, Inflammatory and Other. **G)** Scheme showing the experimental procedures using WT and NGL-1 KO bone marrow cells to derive macrophages *in vitro* using m-CSF for 5 days. After that, the macrophages were stimulated (positive, pos) or not (negative, neg) with LPS + IFNγ overnight, and the conditioned media was recovered to evaluate the production of cytokines by ELISA. Figure made with BioRender.com **H)** ELISAS for TNF⍺, IL-6, IL-10 and Nitrites (as a measurement of activated macrophages) in the conditioned media from stimulated or unstimulated macrophages from WT and NGL-1 KO mice. Macrophages lacking NGL-1 produced decreased levels of the cytokines TNF⍺ and IL-6, as well as Nitrites. N = 3 – 6 independent experiments, with up to 2 mice/group/experiment. Graphs depicted mean ± SEM, One-Way ANOVA, Tukey’s multiple comparisons test. For all of the graphs, * p < 0.05, *** p < 0.001, **** p < 0.0001. ns: not significant.

Next, we inquired if NGL-1 also plays a role in modulating macrophage functions (Fig. 6G). BM derived macrophages from naïve NGL-1 KO mice produced less TNFα, IL-6 and Nitrite (indicative of iNOS activity and macrophage activation) compared to those from WT mice, when stimulated with LPS, and had no differences in the production of IL-10 (Fig. 6H, Sup. Fig. 14B). This phenotype is of a particular interest in PDAC, as TNFα and IL-6 are known TME factors that promote a pathogenic inflammatory response as well as pro-tumoral immunosuppression (90,105). Collectively, these results suggest that NGL-1 might have different roles in assorted immune cell subsets; while the presence of NGL-1 suppresses T cell proliferation and functionality, it promotes macrophage activation and production of cytokines. In PDAC, targeting NGL-1 could be an attractive approach since cytotoxic T cells are usually inhibited or exhausted, and because macrophages are mostly immunosuppressive. Broadly, these results suggest that targeting NGL-1 may restore anti-PDAC immune cell functions.

We explored this exciting premise by further characterizing differences between immune cells from WT and NGL-1 KO mice. For this, we sorted by flow cytometry CD45^+^CD90^+^CD8^+^ T cells and CD4^+^ T cells, CD45^+^CD11b^+^Gr1^+^ cells (granulocytes) and CD45^+^CD11b^+^Gr1^-^F4/80^+^ macrophages from the pancreatic tumors of WT and NGL-1 KO mice (Sup. Fig. 14C) and performed bulk RNAseq analyses. For rigor, we ran a correlation analysis between biological replicates, and confirmed that all were above 0.92, suggesting that the biological replicates were homogeneous and there were no outliers (Sup. Fig. 14D). All immune cell subsets isolated from the NGL-1 KO mice presented transcriptional changes compared to WT cells (Sup. Fig. 15A-B, 16A-B, Sup. Table 9). CD8^+^ T cells from tumors in NGL-1 KO mice upregulated antigen binding and presentation signatures, allograft rejection, and different pathways related to induction of immune response, indicative of activation (Sup. Fig. 15C, Sup. Table 10). Similarly, the tumors collected from NGL-1 KO mice included CD4^+^ T cells that encompassed constitutive upregulation of mTOR signaling, indicating active protein synthesis and thus suggestive of proliferative or secretory cells (Sup. Fig. 15D, Sup. Table 10). Associated with this phenotype, downregulation of HIF1α, fatty acid metabolism and arachidonic acid metabolism were also noted, resembling the metabolic state of memory T cells (106). Moreover, NGL-1 KO CD4^+^ T cells upregulated an IL-17 signature, while lowering activin signaling (i.e., TGF-β pathway), both known to be involved in CD8^+^ T cells suppression (107) (Sup. Fig. 15D, Sup. Table 10). The upregulation of the IL-17 related pathways by the NGL-1 KO cells might be a compensatory mechanism, since tumors in NGL-1 KO mice produced less IL-6 and TGF-β compared to tumors in WT mice, which are important for Th17 differentiation (108,109) (Sup. Table 10). Macrophages (F4/80^+^ cells) from tumors in NGL-1 KO mice upregulated pathways indicative of promoting an anti-tumor response, such as T cell receptor signaling and natural killer (NK) mediated cytotoxicity and signaling (Sup. Table 10). Accordingly, we detected upregulation of IL-12 and IL-23 signaling, which might also be driving the IL-17 profile in T cells (Sup. Fig. 16C, Sup. Table 10). Similarly, the NGL-1 KO granulocytes upregulated pathways related to antigen presentation and allograft rejection, T cell receptor signaling, and NK cell mediated cytotoxicity, while downregulating oxidative phosphorylation (Sup. Fig. 16D, Sup. Table 10). In sum, targeting NGL-1 has a broad effect in the immune TME, affecting the functions of these cells in favor of a more robust anti-tumor phenotype.

### Inhibition of NGL-1 in the TME is as effective as chemotherapy in pancreatic cancer models ***in vivo*.**

Knowing that NGL-1 loss in the TME resulted in decreased tumorigenesis, we questioned how this effect compares to the antitumor effect of chemotherapy. For this we treated tumor allografted WT and NGL-1 KO mice with the approved regimen FOLFIRINOX, a combination of chemotherapeutic drugs that is commonly used as standard of care for pancreatic cancer (110). Mice were treated twice a week for 3.5 weeks, starting 5 days after tumor cell injection into the pancreas (Fig. 7A). Due to the short term of this experiment, treated mice did not display any signs of toxicity, such as significant body weight loss (Sup. Fig. 17A) or changes in behavior (data not shown). Chemotherapy inhibited tumor growth in WT mice by 47% compared to PBS treated controls (Fig. 7B and C). As before, NGL-1 KO mice had 38% smaller tumors even without chemotherapy, mimicking its effect. Combining NGL-1 KO with chemotherapy did not significantly further reduce tumor size, suggesting that targeting NGL-1 alone is as effective as chemotherapy in inhibiting tumor growth (Fig. 7B and C), although the treatment of the NGL-1 KO group with chemotherapy was promising, with mice displaying smaller tumor burden (54% smaller than PBS treated mice). The tumor areas in all groups were mostly poorly differentiated, and there were no differences in percentage of proliferation between the different groups (Sup. Fig. 17B and C). Conversely, all groups receiving chemotherapy displayed more apoptotic cell death, measured by increased percentage of cleaved caspase 3 positive cells, with more cell death in the NGL-1 KO mice treated with chemotherapy (Sup. Fig. 17D). Further, there were no statistical differences noted between the experimental groups when assessing the percentages of different immune cells, as assessed by flow cytometry (Sup. Fig. 17E). There were tendencies for decreased overall percentages of total immune cells (CD45^+^ cells) in the chemotherapy treated groups, mostly represented by smaller percentages of T cells (CD3^+^ T cells) and dendritic cells (CD11c^+^ cells) (Sup. Fig. 17E). These results indicate that targeting stromal NGL-1 was effective at reducing tumor burden, and as effective as chemotherapy alone, with potentially less toxic side effects.

**Figure 7.**
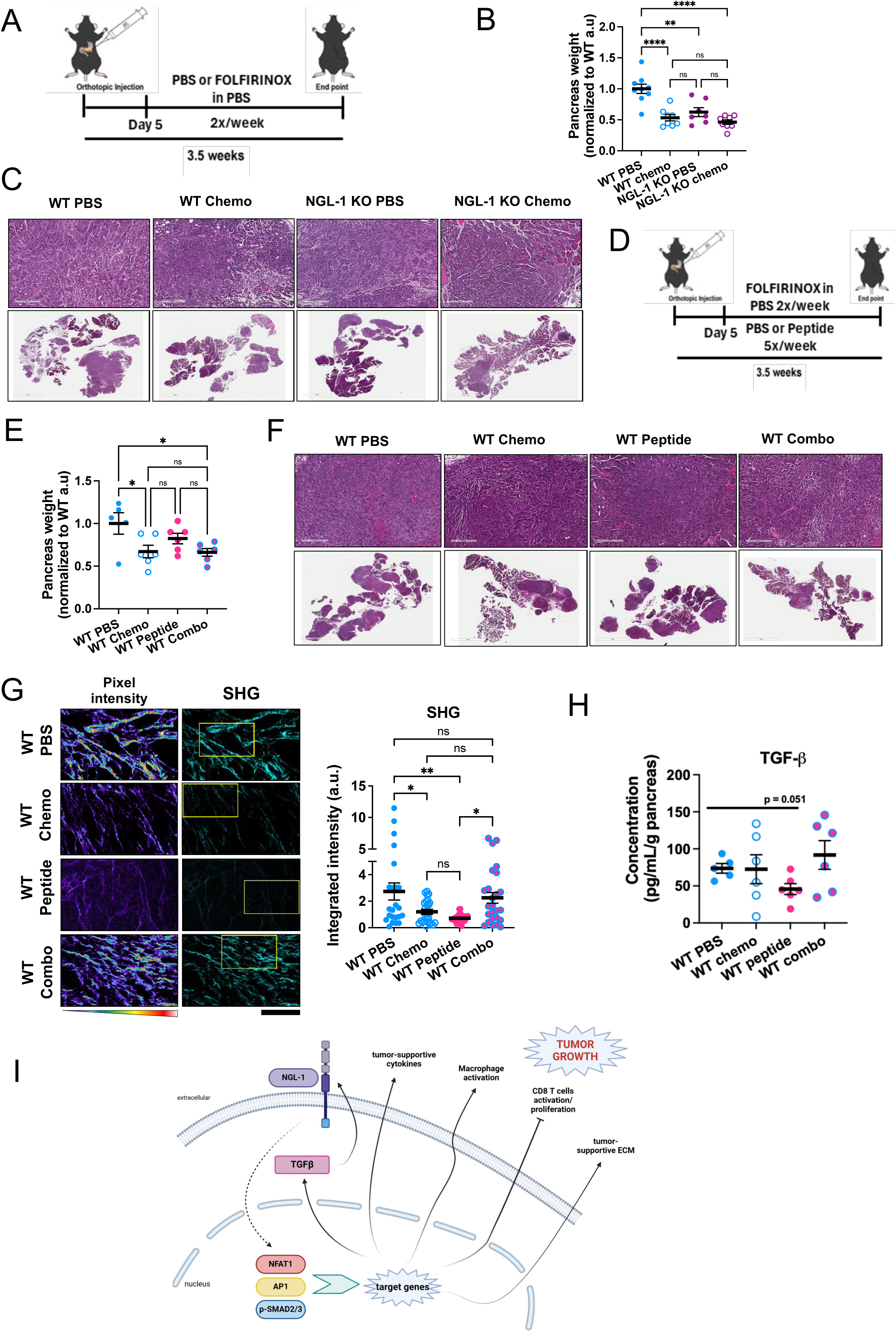
Loss or inhibition of NGL-1 is equally as effective as chemotherapy in orthotopic PDAC models. **A)** Schematics of the orthotopic allografts using WT and NGL-1 KO mice. 5 days post injection with KPC3 cells, mice were treated intraperitoneally with either PBS (10 mL/kg) or FOLFIRINOX (Leucovorin Calcium 50mg/Kg, 5-Fluorouracil 25 mg/Kg, Irinotecan hydrochloride 25 mg/Kg and Oxaliplatin 2.5 mg/Kg in PBS), 2 times per week until the end of the experiment (3.5 weeks post allografts). **B)** Pancreas weight normalized to the average of the WT PBS group, as a measurement of tumor burden. Pancreas weight was smaller in the WT mice treated with FOLFIRINOX (chemo) and in the NGL-1 KO groups (both treated with PBS and with chemo) compared to WT PBS group. N = 7 - 9 mice per group. One-Way ANOVA, Tukey’s multiple comparisons test. **C)** Representative images of the pancreas from WT and NGL-1 KO mice, treated with PBS or chemo, stained with H&E. Scale bars: 7mm and 300 µm. **D)** Scheme of the experimental design for the co-treatment with peptide and FOLFIRINOX. 5 days post injection with KPC3 cells, WT mice were treated intraperitoneally with either PBS (10mL/kg), FOLFIRINOX (Leucovorin Calcium 50mg/Kg, 5-Fluorouracil 25 mg/Kg, Irinotecan hydrochloride 25 mg/Kg and Oxaliplatin 2.5 mg/Kg in PBS) alone, peptide (250 µM in PBS) alone or the combination (combo, peptide + FOLFIRINOX), 2 times per week with chemo, 5 times per week with peptide, until the end of the experiment (3.5 weeks post allografts). **E)** Pancreas weight normalized to the average of the WT PBS group, as a measurement of tumor burden. Pancreas weight was smaller in the WT mice treated with chemo and combo, with a tendency for smaller pancreas weight in the group treated with peptide alone, compared to WT PBS group (p = 0.29). N = 5 - 6 mice per group. One-Way ANOVA, Tukey’s multiple comparisons test. **F)** Representative images of the pancreas from WT mice treated with chemo, peptide or combo, stained with H&E. Scale bars: 7mm and 300 µm. **G)** SHG analysis of the collagen patterns, pseudo-colored in cyan (right panel). The left column displays images color-coded based on pixel intensity heatmaps derived from the total SHG signal. The color scale transitions from cooler to warmer tones, indicating lower to higher pixel intensity signals. Scale bar: 50 μm. The graph shows the quantification of the integrated intensity of the pixels in arbitrary units. N = 3 mice per group, 25 images per group. One-Way ANOVA, Tukey’s multiple comparisons test. **H)** ELISA detecting the production of TGF-β by fragments of pancreatic tissue from mice of the different groups, showing the tendency for decreased levels of TGF-β in the group treated with peptide alone. The fragments of pancreas were weighed and cultured for 24 hours in DMEM media without serum. The cytokine concentration was calculated (pg/mL), normalized by the weight of each fragment. One-Way ANOVA, Tukey’s multiple comparisons test. **I)** Summary of the main findings. NGL-1 modulates the expression of transcription factors NFAT1, AP1 family and p-SMAD2/3, leading to changes in the expression of downstream targets. One of the major factors regulated by NGL-1 is TGF-β, which participates in a feed forward loop, further promoting NGL-1 expression and signaling. This circuit leads to the production of tumor-supportive cytokines and chemokines, production of a tumor-supportive ECM, inhibition of CD8 T cell functionality and intrinsic modulation of macrophage activation and T cell activation and proliferation. The combination of these functional outputs results in tumor growth and progression. Figure made with BioRender.com. All graphs depict mean ± SEM. * p < 0.05, ** p < 0.01, **** p < 0.0001. ns: not significant.

### A small peptide targeting NGL-1 phenocopies the genetic loss of NGL-1, validating NGL-1 as a potential therapeutic target in PDAC

Given the promising results with global loss of NGL-1 *in vivo*, we developed a therapeutic approach to target NGL-1 in mice. Using a publicly available data on the crystal structure of NGL-1/NetG1 in complex (PDB: 3ZYJ) (111), we designed a small peptide (here called anti-NGL-1 peptide, or peptide), composed of 8 amino acids (NPYMSNNE), simulating a fragment of the NGL-1 binding domain of NetG1 (the NGL-1 discriminant loop I of residues 84-91 of NetG1 with C88S mutation), which was hypothesized to function as an inhibitor (or antagonist) of NGL-1 signaling (image and predicted binding site of the crystal structure on Sup Fig 18A and B). The ability of the peptide to bind NGL-1 was confirmed *in vitro* using a “sandwich ELISA” approach, in which the plate was coated with either recombinant NetG1 (positive control, 50nM), BSA (negative control, 50nM), or anti-NGL-1 peptide (1mM), followed by recombinant NGL-1 incubation and probed with anti-NGL-1 antibody (Sup. Fig. 18C). Recombinant NGL-1 bound equally well to both recombinant NetG1 and the peptide, confirmed by detection of the anti-NGL-1 antibody-HRP conjugate (Sup. Fig. 18C). Next, we tested if the treatment of CAFs with this peptide would phenocopy the knockdown of NGL-1. Strikingly, CAFs treated with the peptide at nanomolar concentrations, during *in vitro* CAF/ECM unit culture, produced less IL-6 and IL-8 compared to CAFs treated with vehicle control, thereby mimicking the genetic loss of NGL-1 (Sup. Fig. 19A). Therefore, we demonstrated functionally that this small peptide can bind to and inhibit the function of NGL-1.

These *in vitro* results suggested that the anti-NGL-1 peptide could reproduce the effects of the loss of NGL-1 in mice. Prior to defining the final working dose for treatments *in vivo*, we treated WT mice (n = 2) intraperitoneally with 2.5 mM peptide to evaluate possible acute toxicity. Mice were observed continuously for 4 hours after treatment and then daily for 15 days after treatment. No signs of toxicity were observed, as well as no loss of body weight (data not shown). We also confirmed absence of toxicity in animals treated with peptide (250 µM in PBS) for the length of the orthotopic allografts experiment (3 weeks). We orthotopically injected PDAC cells into WT mice and treated them with 250 µM of peptide, intraperitoneally, five days/week, starting five days after tumor cell allografting (Sup. Fig. 19B). As controls, we evaluated tumor burden in NGL-1 Het and KO mice treated with PBS. There were no differences between groups in body weight gain, suggesting absence of toxicity for 3 weeks (Sup. Fig. 19C). Similarly to tumor bearing Het and KO hosts, we noted a tendency for decreased pancreata weight (24% het and 38% NGL-1 KO, compared to WT group treated with PBS) in the peptide-treated group (37% decrease) (Sup. Fig. 19D). There were no differences between groups in tumor differentiation status (most were poorly differentiated) nor in the percentage of cells positive for Ki67 (Sup. Fig. 19E and F). Interestingly, all the experimental groups presented with less cell death compared to the WT control group, as measured by cleaved caspase 3 (Sup. Fig. 19G). These results suggest that the peptide might phenocopy NGL-1 loss *in vivo*.

Because the genetic loss of NGL-1 in mice had a comparable efficacy to chemotherapy alone (Fig. 7A), we decided to treat a group of mice with FOLFIRINOX and another with both FOLFIRINOX and peptide (combo), to inquire if the peptide was as effective as chemotherapy (Fig. 7D). The peptide treatment resulted in tumors 20% smaller than the control treated with PBS (Fig. 7E). The groups treated with chemotherapy alone or combo presented tumors 33% and 34% smaller than the control group respectively, suggesting a similar phenotype observed for the NGL-1 KO mice, although less prominent (Fig. 7E). Further, there were no differences observed between groups in tumor differentiation status, with most of PDAC area being poorly differentiated (Fig. 7F, Sup. Fig. 20A). The mice treated with chemotherapy presented increased proliferation (p < 0.05) and cell death (p < 0.0005) compared to the PBS treated group, with no statistical differences between the PBS and the peptide treated groups for any of these parameters (Sup. Fig. 20B and C). We also assessed immune cell composition in the tumors of all the groups. There were decreased percentages of total immune cells (CD45^+^ cells) in the chemotherapy treated groups, mostly due to less CD3^+^ T cells, dendritic cells (CD11c^+^) and monocytes (Ly6C high^+^ cells), while some of the mice treated with peptide alone tended to present increased percentages of CD4^+^ T cells and NK cells (Sup. Fig. 20D). Despite the discrete effects on tumor burden, the treatment with peptide phenocopied the tumor bearing NGL-1 KO hosts with decreased collagen bundling and maturation compared to PBS and combo treatments, suggesting modulation of the ECM compartment towards a less tumor supportive phenotype, as before (Fig. 7G). This observation was accompanied by decreased production of TGF-β, showing again the involvement of NGL-1 in the modulation of this key pro-tumor/immunosuppressive cytokine (Fig. 7H). Taken together, our results provide initial evidence that short-term targeting of NGL-1 with a small peptide in mice mimics genetic loss of NGL-1, tending to normalize the TME in this experimental model of pancreatic cancer through potential downregulation of TGF-β signaling.

## Discussion

PDAC is an extremely aggressive cancer, with only 13% of the patients surviving more than 5 years after diagnosis (1). It is predicted that by 2026, pancreatic cancer will be the second leading cancer-related cause of death in the United States (2). Part of the explanation for such unfavorable outcomes is the fact that diagnosis occurs at late stages, with most of the cases being unresectable (112). Contributing to these abysmal statistics, the unique biology behind PDAC is to be blamed; PDAC features a very distinct TME, characterized by the expansion of CAFs, deposition of a fibrous-like and dense interstitial ECM that triggers the collapse of local vessels, and the establishment of a highly immunosuppressive TME (3,15,17,113,114). CAFs are among the major stromal cell types found in PDAC and are known to sustain the pathological features of this disease (13,16,115–117). This is achieved by their ability to provide metabolic support to surrounding cells (29,31,118–123) and secretion of chemokines and cytokines to modulate immune cell recruitment and function (29,75,124–126). CAFs and their ECM act as a functional unit (CAF/ECM unit) that define and sustain the complex relationship between the different tumor components, especially in PDAC (25–27,127–129), which is critical in the study of CAF biology.

Despite the importance of the stroma in PDAC, there are numerous lessons remaining to be learned. Several studies investigated CAF lineages and subtypes (74,75,124,130–137), from which clearly demonstrated that CAFs are a heterogeneous population, in both their origins and functional phenotypes. These, in part, explain the controversial results obtained when attempts were made to target or eliminate key components of the CAF/ECM units in murine models and in PDAC patients. Therefore, understanding the stromal heterogeneity in PDAC will be imperative to overcome drug resistance and resilience towards immunotherapy. Uncovering new stromal targets and modulating them towards stromal normalization rather than elimination appears to be the best approach for PDAC therapy (24).

The present study uncovered the expression of a post-synaptic protein/receptor, Netrin G1 Ligand (NGL-1), by stromal cells in PDAC. Recently our group reported the ectopic expression of NetG1, the binding partner and ligand for NGL-1, in fibroblasts from PDAC patients. We also reported the expression of NGL-1 in cancer cells, with NetG1 and NGL-1 having both common and independent pro-tumor effects. Specifically, while NetG1 was found to promote pro-tumor functions of CAFs, cancer cells lacking NGL-1 presented impaired ability to scavenge nutrients through macropinocytosis and to develop tumors *in vivo* (29). NGL-1, or LRRC4C (leucine-rich repeat-containing 4C), is a type I transmembrane postsynaptic adhesion molecule that interacts intracellularly with postsynaptic density protein-95 (PSD-95) and binds to the presynaptic membrane protein NetG1 (40,138). NGL-1 is vertebrate-specific and found in excitatory glutamatergic synapses (139). Structurally, NGL-1 contains an extracellular region with ten leucin-rich repeats (LRRs), a cysteine-rich capping domain in each LRR end and a C2-type immunoglobulin (Ig) domain. These are followed by a single transmembrane domain and a cytoplasmic domain that interacts with a PDZ binding motif (139,140). It has been shown that NGL-1 mutant mice present hyperactive, anxiolytic-like behavior and impaired learning and memory (40). Otherwise, in our experiments, we observed that these mice are healthy and fertile. Functionally, members of the LRR family of proteins are involved in the development of the nervous system and in the regulation of innate immune response (141). Accordingly in this study, we show that NGL-1 controls several functional aspects in CAFs, including those involved with ECM remodeling and immunosuppression. Moreover, we also demonstrated that NGL-1 influences the pro-tumoral functionality of immune cells such as macrophages and T cells, reporting for the first time the functional roles of NGL-1 in stromal cells.

Cell adhesion molecules at synaptic sites, such as NGL-1, are known to be multifunctional proteins involved in the coordination of assorted aspects of development, as well as cellular function and plasticity (142). Despite that, only a few studies propose a role for NGL-1 in cancer and in other diseases (29,31,140,143). Analyzing RNAseq data from patients with colon cancer and stomach cancer (available at TCGA) using the algorithm “Estimation of STromal and Immune cells in MAlignant Tumor tissues using Expression” (ESTIMATE), Yang et al found that *LRRC4C* was indicative of an altered TME. They observed that NGL-1 expression correlated with disease progression and inversely correlated with overall survival (143).

In the present study, we too observed that NGL-1 expression correlated with disease progression, being overexpressed in tumor-adjacent and in the tumor-associated stroma compared to the stroma in normal pancreas in tissue from PDAC patients. In accordance, we reported previously that high expression of tumoral NGL-1 correlated with the basal classification of pancreatic cancer, which presents worse prognosis (29). Here, the expression of NGL-1 in fibroblasts inversely correlated with overall survival in two different cohorts of PDAC patients, indicating a functionality for NGL-1 in stromal cells. It is important to note that there was a trend for the expression of NGL-1 in immune cells to also inversely correlate with overall survival. We believe that this is because immune cells overexpress NGL-1 in early stages of the disease, which we observed when analyzing murine neutrophils (Ly6G^+^) and CD8^+^ T cells for NGL-1 expression that were collected at different time points of disease progression. Moreover, tissue from genetic murine models for pancreatic cancer bearing precursor pancreatic lesions, such as ADM and PanINs, have a higher percentage of TME cells expressing NGL-1 compared to normal tissue, with this percentage being highest in PDAC tissues. Despite being a receptor-ligand pair, NetG1 and NGL-1 expression only weakly correlated when queried in human PDAC tissues, suggesting that these proteins have both dependent and independent roles from each other, as previously shown by our group (29,31).

Our study suggests that NGL-1 is a pro-tumoral TME protein and a viable stromal target in PDAC. We found that mice germline knockout and heterozygous for NGL-1, orthotopically allografted with PDAC cells, developed smaller tumors compared to control mice, with more consistent results in older mice. We were surprised to see such an effect in this experimental model, since orthotopic allografts tend to be very aggressive and to not display such prominent stromal expansion as the genetic models (144). But despite that, modulating host expression of NGL-1 was sufficient to limit the growth of the pancreatic cancer allografts, suggesting that stromal NGL-1 has a strong effect on tumor cells. Confirming this premise, scRNAseq analysis comparing tumors formed in control vs. NGL-1 KO mice, showed that these were molecularly distinct, despite orthotopic introduction of the same PDAC cancer cells in the distinct hosts.

As pointed out earlier, one of the features of PDAC stroma is the deposition of a dense ECM, that can be tumor restrictive or tumor supportive. Functionally, the ECM produced by NGL-1 KD CAFs was less supportive of PDAC cell survival under nutrient deprivation, a phenotype that was also observed with CAFs downregulating NetG1 (29). Here, we also show that modulating stromal NGL-1 changed patterns of collagen bundling and maturation in the tumors, while maintaining the expression of Col1A1, Col1A2 and Col3A1, known to be abundant in PDAC (128). Higher levels of collagen bundling, maturation and alignment, together with higher ECM stiffness correlates with a CAF signature and worse prognosis in PDAC (145–148). Moreover, a recent study showed that ablation of Col1A1 in α-SMA^+^ fibroblasts led to progression of pancreatic cancer (20). Therefore, loss of NGL-1 in CAFs did not disrupt the formation of a CAF/ECM unit, but instead limited its pro-tumoral function, which is especially important because the ablation of CAF units is known to be detrimental to survival (19–23). One of the major factors modulating the different CAF-ECM phenotypes, as well as immunosuppression and resistance to therapies in PDAC is TGF-β (149–154). Here we show that tumors growing in NGL-1 KO hosts downregulated many factors involved in this pathway across different cell types, including p-Smad2, an indication that TGF-β signaling is downregulated upon loss of NGL-1. Moreover, the secretion of TGF-β was also decreased in the tumors from NGL-1 KO mice, illustrating their influence to generate a less tumor supportive TME. In agreement, the bone marrow chimera experiment confirmed that NGL-1 pro-tumor effects are dependent on TGF-β.

Immunosuppression is an important feature of PDAC, characterized by increased secretion of cytokines, chemokines, abundance of immunosuppressive cells and absence or dysfunction of antitumor immune cells, such as CD8^+^ T cells and NK cells (155). This, together with having one of the lowest mutational burdens, explains why PDAC is resistant to immunotherapy (156–158). Here we found that NGL-1 KO tumors produced more CXCL9, CXCL10 and IFNγ, factors known to promote immune response (44,159–163). This was associated with increased percentages of CD8^+^ T cells in the NGL-1 KO tumors compared to control tumors, which presented a naïve/stem cell-like phenotype and downregulated immune checkpoint molecules. Several studies report that naïve-like and stem cell-like memory T cells tend to respond better to immune checkpoint inhibition (164–166). Furthermore, naïve T cells from NGL-1 KO mice proliferated more than the control cells when stimulated *in vitro*, while also upregulating pathways related to innate immune response and antigen presentation response (according to our scRNAseq and bulk RNAseq data). These results suggest that the NGL-1 KO T cells are more capable of engaging in an immune response in the TME. It is also important to emphasize that T cells from NGL-1 KO tumors downregulated inhibitory markers such as *Tox, Ctla4, Pdcd1* (PD-1), suggesting that NGL-1 KO T cells were less dysfunctional than control cells. In fact, the NGL-1 KO T cells were more polyfunctional than WT T cells, producing two or more cytokines upon stimulation, and expressed less TGF-β (scRNAseq data), which also plays a role in T cell suppression, especially through production of this immunosuppressive cytokine by Tregs (167–169). These results suggest that modulation of NGL-1 in the T cell compartment leads to reprogramming of these cells to be less tumor supportive.

It is known that CAFs, myeloid cells and Tregs are important components of the immune cell landscape in PDAC (90,126,170–174). Tumors from NGL-1 KO mice produced less GM-CSF, G-CSF and IL-6, cytokines known to attract and promote differentiation of granulocytes and other cells in the TME (51,175). Moreover, NGL-1 KD CAFs produced less cytokines and chemokines such as GM-CSF, IL-6, IL-8 and TGF-β than control CAFs, which together with the higher expression of antigen-presenting factors and low PDL-1 expression, further confirms the immunomodulatory nature of NGL-1 in different cell compartments in the TME. CAFs are recognized for secreting factors that enhance pro-tumor immune cell function (16). Additionally, recent research indicates that CAFs can establish immunological synapses with regulatory T cells, thus fostering tumor development (176). This suggests that CAFs, and possibly immune cells expressing NGL-1, may exert direct cell-cell interactions, beyond their secretory effects, on various cells within the TME through heterotypic synapses, such as immunologic ones. Myeloid cells derived from NGL-1 KO tumors downregulated expression of immunosuppressive genes, such as *Arg1*, *Apoe*, *Trem2*, *C1qa*, *C1qb* and *C1qc*. *Arg1* is known to be upregulated in immunosuppressive myeloid cells in PDAC and other types of cancer (174,177,178). In the same way, *Trem2*, *Apoe* and genes related to the complement pathway are also involved in the immunosuppressive phenotype in myeloid cells, highlighting the role of NGL-1 in myeloid cell function in the TME (57,58,179,180). Functionally, bone marrow derived macrophages from NGL-1 KO mice stimulated *in vitro* present decreased production of IL-6 and TNFα, suggesting an overall modulation of their functionality.

A common theme among the pathways downregulated in assorted cells from NGL-1 KO tumors was immunoregulation. There was overall downregulation of genes in the “coronavirus pathogenesis pathway,” “IL-6 pathway” and others related to immune cell function. In fact, proteins displaying the LRR domain are largely involved in innate immunity (181), and our study places NGL-1 for the first time in this scenario. A plausible explanation is the fact that genes known to be related with the AP-1 transcription factor family were downregulated in the NGL-1 KO tumors. It is known that AP-1 members (Jun and Fos proteins), in conjunction with NFAT (also downregulated in NGL-1 KO tumors), act to modulate the expression of genes important for immune cell activation, function and cytokine secretion (182–185). Moreover, a recent study linked the AP-1 factors *Junb* and *Fosl1* (FRA-1) with a proto-oncogenic transcriptional program driven by *Kras*G12D mutations, the most common *Kras* mutation in pancreatic cancer, to unleash tumorigenesis in the context of chronic inflammation (186). Accordingly, here we uncovered a new connection between NGL-1, NFAT and the AP-1 family proteins, whose modulation leads to a tumor restricting TME in PDAC.

One of the most important players in PDAC is the CAF/ECM unit (26). Several phenotypic states of CAFs can be found in the TME, exerting major functions such as modulation and deposition of ECM and immunomodulation (75,124,133,135,136,187). While these states might suggest functional polarizations, CAFs are highly plastic, and they can exert multiple phenotypes (188). For instance, we previously demonstrated that NetrinG1^+^ CAFs, while being immunosuppressive, maintain their myofibroblastic phenotype (29). Here, we showed that NGL-1 is important for both myofibroblastic and immunomodulatory features of CAFs. NGL-1 KO CAFs downregulated immunosuppressive factors, such as TGF-β and PDL-1, and upregulated markers related to antigen presentation, while maintaining the expression of collagens important for the structure of the PDAC TME, suggesting the establishment of a more immune reactive stroma, without compromising its structure. In fact, bulk RNAseq comparing control and NGL-1 KD CAFs showed upregulation of *Col1a1*, *Col1A2* and *Cxcl10*, with downregulation of *Cxcl8*, *Fosl1* (encoding FRA-1), *Nfatc2* (encoding NFAT1), and *Lrrc15*. LRRC15^+^ CAFs display both myofibroblastic and immunosuppressive features, in a TGF-β-dependent manner (124,189). Interestingly, NGL-1^+^ CAFs share many similarities with LRRC15^+^ CAFs, including the connection with TGF-β and the fact that NGL-1 KD CAFs upregulate a more universal fibroblastic signature marked by upregulation of JAK-STAT, NF-κB and TNF signaling, similarly to fibroblasts depleted of LRRC15 (189).

In this study, we uncovered that NGL-1^+^ CAF functions are also dependent on TGF-β, since loss of NGL-1 led CAFs to express decreased levels of myofibroblastic markers known to be involved in TGF-β signaling, such as PallD iso 3 and 4 and p-Smad2/3, while also producing less TGF-β. These factors have also been linked to the immunosuppressive features of CAFs (27,87). Interestingly, while loss of NGL-1 in CAFs ablated their ability to produce TGF-β, they still respond to it, showing that a systemic inhibition of NGL-1 could lead to inhibition of TGF-β-associated signaling and tumorigenesis. Surprisingly, modulation of TGF-β controls NGL-1 expression, suggesting the existence of a positive feedback loop. The LRRC transmembrane proteins LRRC32 and LRRC33 can induce retention of TGF-β at cell surface, providing a localized source for TGF-β activation (190–192). This might be the case of NGL-1, but further studies are needed to confirm this hypothesis.

Loss of NGL-1 in CAFs also led to decreased expression of AP-1 transcription factors (FRA-1, FRA-2, FosB, c-Fos and c-Jun), NFAT1 and β-catenin, all factors deeply involved in tumor development and immunity (193,194). Interestingly, while overexpression of NFAT1 and induction of TGF-β signaling independently induced the expression of NGL-1, TGF-β signaling did not rescue the expression of NFAT in the absence of NGL-1, suggesting that NFAT1 expression is dependent on NGL-1 in CAFs. Conversely, loss of NFAT1 only partially phenocopies the loss of NGL-1, showing that other factors may be involved in mediating the NGL-1 phenotype. The relationship between NFAT1 and TGF-β was described before (195–197), and here, for the first time, we show NGL-1 as a key regulator of these pro-tumor factors.

Finally, it is known that chemo and radiotherapy can promote resistance through deposition of collagen and induction of pro-tumoral cytokines, among other mechanisms (87,198,199). Different from the established therapies, targeting NGL-1 (either genetically or using a small prototype peptide from NetG1) was as effective as chemotherapy in decreasing tumor burden, while reprogramming the stroma to be less tumor supportive, such as decreased collagen bundling and maturation in the tissue treated with peptide targeting NGL-1. In addition, targeting NGL-1 inhibits TGF-β signaling, but potentially without the side effects of complete ablation of TGF-β itself. It is important to point out that despite not leading to a decrease in tumor burden that was evident in NGL-1 KO mice, the effects of the anti-NGL-1 peptide in the TME are encouraging.

Further dosing strategies, length of treatment, and optimization of peptide structure will be explored in the future with the goal of improving its therapeutic benefits. Overall, in this study, we introduce NGL-1, a synaptic protein, as a potential stromal target in PDAC, whose inhibition modulates, rather than eliminates, the physical stroma and reduces immunosuppression. We further propose NGL-1 as a new immunomodulatory molecule, and its inhibition may improve anti-tumor T-cell functionality.

### Material and Methods

*(Information about key reagents and resources can be found in the Key Resources Table)*

#### Ethical Collection of Human Samples

Written informed consent was obtained for every patient who donated samples for research purposes. The collection of human samples was approved by the Institutional Review Boards from Fox Chase Cancer Center and MD Anderson. Fox Chase Cancer Center adheres to the International Society for Biological and Environmental Repositories and National Cancer Institute Best Practices for Biospecimen Resources and participates in the NCI Office of Biorepositories and Biospecimen Resources and Biospecimen Research Network.

#### Primary Cancer Associated Fibroblasts (CAFs) from human PDAC tissue

Primary CAFs used in this study were previously isolated and characterized as described (78). Briefly, pancreatic tissue from PDAC patients was collected in phosphate buffered saline (PBS) containing 2% Penicillin-Streptomycin (P/S) (10,000 U/mL) and 1% Amphotericin B. The tissue was minced with sterile scalpel in a petri dish containing Dulbecco Modified Eagle Medium (DMEM) supplemented with 3% Bovine Serum Albumin (BSA) and P/S, and digested overnight with collagenase III (2mg/mL). The homogenate was subsequently filtered in 100 and 40 μm filters and the suspensions were spun down at 1500 rpm for 10 minutes. The cell pellet was cultured in flasks containing DMEM + 10% Fetal Bovine Serum (FBS) + 1% P/S for 6 hours, then the medium was refreshed, and only the confluent cells were kept. Fibroblasts were defined as cells lacking cytokeratins and expressing vimentin (by western blotting), plus their ability to secrete substantial extracellular matrix (ECM) (thickness >5 µm and 3 cell layers) and to acquire lipid droplets after TGF-β signaling inhibition (11,12). For immortalization, cells were transduced with pBABE-neo-hTERT vector (gift from Dr. Robert Weinberg, Addgene plasmid # 1774)(12,29).

#### Cell culture

##### Cancer cells

Cancer cells (PANC-1, labeled with mCherry/red fluorescent protein, RFP were maintained by sub-culturing the cells twice a week in DMEM supplemented with 5% FBS and 1% P/S (10,000 U/mL). Murine KPC3 cells, derived from the KPC tumors (citation) were maintained by sub-culturing the cells three times per week, in DMEM supplemented with 10% FBS and 1% P/S. Cells were maintained in incubator at 37°C, 5% CO_2_, and vetted free of pathogens and Mycoplasma (IDEXX Bioresearch and Invivogen) prior to injections.

##### Cancer associated fibroblasts

CAFs were maintained by sub-culturing the cells twice a week in DMEM supplemented with 15% FBS, 1% P/S and 4 mmol/L L-glutamine. In certain experiments, CAFs were cultured in DMEM without FBS and L-glutamine (to be specified in the corresponding methods sections and figure legends).

##### Immune Cells

Functional assays with immune cells were performed in Roswell Park Memorial Institute medium 1640 (RPMI) supplemented with 10% heat inactivated FBS, 1% P/S and 50 mM 2-Mercaptoethanol.

#### Cell line authentication and *Mycoplasma* detection

All cell lines were authenticated by IDEXX BioResearch and genetic analysis was performed to determine the species of origin and the genetic profile based in 16 different allelic markers (AMEL, CSF1PO, D13S317, D16S539, D18S51, D21S11, D3S1358, D5S818, D7S820, D8S1179, FGA, Penta_D, Penta_E, TH01, and vWA).

All lines were regularly tested for *Mycoplasma*, using PCR or PCR plus immunochromatographic strip detection methods, according to the manufacturer protocols. Every cell line used in this study tested negative for *Mycoplasma*.

#### CRISPR interference (CRISPRi) mediated knockdown of target genes

Using the CRISPRi-Puro plasmid generated previously (29), gRNAs targeting the promoter regions of target genes were obtained from the study “Compact and highly active next-generation libraries for CRISPR-mediated gene repression and activation.” The top two oligos for each target gene generated from this study were cloned into the CRISPRi-Puro vector, as described below in *gRNA cloning and vector generation.* Functional lentivirus production, transduction into target cells, and selection of transduced cells was performed as described below in *Lentivirus Production and Transduction of Target Cells*. Cells transduced with empty vector alone were used as controls.

gRNA sequences are as follows (overhangs added, ***bold and italics***):

NGL-1 CRISPRi 1.1 **<u>CACC</u>**GAAATGGTAAGAGGAATGGG NGL-1 CRISPRi 1.2 **<u>AAAC</u>**CCCATTCCTCTTACCATTTC NGL-1 CRISPRi 2.1 **<u>CACC</u>**GTAAGCCAAAAACTCATCAA NGL-1 CRISPRi 2.2 **<u>AAAC</u>**TTGATGAGTTTTTGGCTTAC NFAT1 P1 CRISPRi 1.1 ***CACC***GCGCGCCCGGGGAAGCTGAG NFAT1 P1 CRISPRi 1.2 ***AAAC***CTCAGCTTCCCCGGGCGCGC NFAT1 P2 CRISPRi 1.1 ***CACC***GGCGATCCGGCTTACTCCAG NFAT1 P2 CRISPRi 1.2 ***AAAC***CTGGAGTAAGCCGGATCGCC

##### gRNA cloning and vector generation

gRNAs (1.1 with 1.2, 2.1 with 2.2 were annealed in a thermal cycler and phosphorylated using T4 PNK (New England Biolabs). The annealed and phosphorylated gRNAs were diluted 1:200 in molecular biology grade water (Ambion) and ligated (Quick Ligase Kit, New England Biolabs) into the Esp3I (New England Biolabs) digested, dephosphorylated (Fast Alkaline Phosphatase, Thermofisher) CRISPRi-Puro vector. Ligated vectors were transformed into competent Stbl3 bacteria and plated onto LB agar plates containing 100 µg/mL ampicillin to select for bacteria containing ligated vectors. Colonies were selected and screened by colony PCR (using forward primer for U6 promoter [GAGGGCCTATTTCCCATGATT] and 0.2 gRNA oligo as the reverse primer) to confirm successful cloning of gRNAs. Positive colonies were expanded, and the plasmid DNA was purified with the ZymoPURE II Plasmid Midiprep Kit (Zymo Research, #D4201). Purified samples were sequenced to ensure proper cloning of gRNA sequences and subsequently used to generate functional lentivirus for transduction.

##### Lentivirus Production and Transduction of Target Cells

Lentiviruses were produced in 293T cells by transfecting 10 μg LentiCRISPRv2, 5 μg psPAX2 (a gift from Didier Trono; Addgene plasmid # 12260), 2 μg VSVg, and 30 μL X-tremeGene9 transfection reagent (Sigma-Aldrich) that was pre-mixed for 30 min in 1 mL of serum/antibiotic free DMEM. The resultant transfection mix was then added dropwise to 293T cells containing 5 mL of serum/antibiotic free DMEM and cells were incubated overnight at 37°C. The following day, the media was aspirated and 10 mL of complete DMEM was added to the 293T cells. Next, on days 2 and 4 post-transfection, media was collected and passed through a 0.45 μm filter. Lentiviral media was then used immediately to transduce target cells in the presence of 10 μg/mL polybrene (Santa Cruz Biotechnology) or stored at -80°C for future use. After 72 hours, target cells were selected with puromycin (2 μg/mL) and cells that survived selection for 2 weeks were used in experiments. Cells transduced with empty vector were used as a control.

#### Overexpression of NFAT1 in Target Cells

##### Cloning of NFAT1 (NFATC2 gene) into lentiviral overexpression plasmid

*NFATC2* (gene encoding NFAT1 protein) was PCR amplified from a plasmid containing NFATC2 gene (Addgene 74050, a gift from Ria Baumgrass)(200), using primers (see below for sequences) that added the Xba/Xho restriction enzyme cut sites flanking the gene for insertion into the lentiviral overexpression plasmid pLV-CMV-H4-puro vector (kindly provided by Dr. Alexey Ivanov, West Virginia University School of Medicine, Morgantown, WV)(29). The resultant PCR product was column purified using the GeneJET PCR Purification Kit (ThermoFisher). The overexpression plasmid was simultaneously cut with Xba/Xho restriction enzymes (New England Biolabs) and dephosphorylated with FastAP Thermosensitive Alkaline Phosphatase (ThermoFisher) and was gel purified using the Zymoclean Large Fragment DNA Recovery Kit (Zymo Research). Finally, the purified PCR product and plasmid were ligated together using the Quick Ligation Kit (New England Biolabs). Ligation products were transformed into STBL3 bacteria (as above) and colonies were screened using the CMV FW primer and reverse primer for NFAT1. Positive colonies were amplified, and plasmid DNA was purified and sequenced (as above). Plasmids containing successfully NFAT1 were used to generate lentiviruses and transduce target cell lines (as outlined above).

The primer sequences are listed below (***bold italics indicates overhangs added***): NFAT1 FW Xba ***TAGGTATCTAGA***atgaacgcccccgagcg

NFAT1 RV Xho **<u>TAGGTACTCGAG</u>**ttacgtctgatttctggcaggagg

CMV FW CGCAAATGGGCGGTAGGCGTG

3D microenvironment: preparation of cell-derived extracellular matrices (CDM) for survival assays

The preparation of CDM was performed as described (12,29,78). Briefly, wells from a 24 wells plate were coated with 0.2% cold water fish skin gelatin (Sigma) for 30 minutes, then washed three times with PBS. Then, wells were crosslinked with 1% (v/v) glutaraldehyde (Sigma) for 30 minutes, washed three times with PBS and incubated with 1M ethanolamine (Sigma) for another 30 minutes, followed by three washes with PBS. CAFs (CON or NGL-1 KD) were then seeded into each well in complete DMEM media (125,000 cells/well in 0.5 mL). The next day, the wells were aspirated and complete DMEM media containing 50 µg/mL L-ascorbic acid was added to the culture. This procedure was repeated for 5 days. After that, cells were lysed and extracted from the CDM using 0.5% Triton X-100 supplemented with 20 mM Ammonia Hydroxide, followed by five washes with PBS. Prior to survival assay experiments, the CDM were treated with DNAse I (2 units/mL) for 4 hours at 37°C and washed 5 times with PBS. The resulting CDM was used as 3D substrate for the survival assays.

#### Preparation of conditioned media (CM) from CAFs

CAF CM was prepared by culturing CAFs (CON or NGL-1 KD) in 6 well plates (500,000 cells/well in 3 mL of complete media) coated with 0.2% gelatin and inducing them to produce CDM (5 days treatment with L-ascorbic acid in complete media as described above). After that, the complete media was replaced with 3 mL DMEM without FBS, and L-glutamine and cells were allowed to condition the media for two days. The CM was collected, aliquoted and frozen at -80°C until use (29). The remaining cells were lysed in RIPA buffer and lysates were collected for western blotting.

#### *In vitro* CAF treatments during CDM and CM production

In some experiments, cells received different treatments during CDM production. In these cases, cells were treated daily from the second day after plating the cells in gelatin, until the last day of CDM production unless stated differently. Treatments were: 10 µM TGF beta receptor I inhibitor (SB-431542, Sigma) and DMSO (Sigma) as a vehicle control, 10 ng/mL recombinant TGF beta (Sigma) and 10 mM citric acid (Sigma) as a vehicle control, and anti-NGL-1 peptide (1.79, 17.9 and 179 nM) and PBS as vehicle control.

#### Immunofluorescence for CDM characterization

CAFs were plated in 24 wells plates (125,000 cells/well, 0.5mL) with coverslips coated with 0.2% gelatin, as described above, and allowed to produce CDM. After 5 days of CDM production, CAFs and their CDM were fixed and permeabilized for 5 minutes at RT in 4% paraformaldehyde containing 0.5% TritonX-100 and 150 mmol/L sucrose and allowed to continue fixation, using the same preparation but lacking Triton, for an additional 20 minutes at RT. Fixed-permeabilized CAFs + CDM were then blocked for 1 hour at RT with Odyssey Blocking Buffer (PBS base; Li-Cor). CAFs + CDM were stained for 1 hour at RT with 1:200 fibronectin antibody diluted in blocking buffer. Then, coverslips were washed 3 times with 0.05% Tween in phosphate-buffered saline (PBST), and the secondary antibody diluted 1:100 in blocking buffer was added for 1 hour at RT (Alexa Fluor 647). Next, coverslips were washed 3 times with 0.05% PBST, and nuclei were counterstained with DAPI (1:20,000; Sigma-Aldrich). Stained coverslips were imaged using the Nikon Confocal A1S system. 5-10 images per coverslip were captured. Fibronectin alignment was quantified as described below.

#### CDM fiber orientation analysis

The analysis of fiber orientation (i.e., the alignment of the fibers) was performed as previously described (78). Briefly, fiber orientation analysis was carried out on the confocal images of fibronectin in CAFs + CDM, using OrientationJ plugin for ImageJ (201). The obtained fiber angles were normalized to fit within the angle range of −90° to +90°, and these were plotted in Excel to generate orientation curves (alignment output at every angle). The percentage of fibers between −15° and +15° was plotted as a measurement indicative of CDM fiber alignment.

#### Lipid Droplets Assay

Lipids Droplets Fluorescence Assay Kit (Cayman Chemicals) was used according to manufacturer’s instructions, with slight modifications. Briefly, 5,000 CAFs/well were plated on glass bottom, black 96 well plates, in complete DMEM media. The following day, CAFs were treated with Oleic Acid (100 µL, 1:1000, provided with kit) to induce lipid droplet formation, as a positive control, or 100 µL extra media, for 24 hours. Next, media was aspirated from wells, and cells were washed (100 µL diluted wash solution, provided with kit) and fixed (50 µL diluted fixative, provided with kit) for 10 min. CAFs were then washed in 100 µL diluted wash solution, followed by 15 min staining of lipids droplets with a 1:1000 diluted Nile Red solution (provided with kit). CAFs were then washed again with 100 µL diluted wash solution, and stained with 100 µL of a 1:30,000 DAPI solution for 10 min, to stain nuclei for normalization of Nile Red signal. Finally, cells were washed with 200 µL diluted wash solution, aspirated, and 100 µL diluted wash solution was added before reading the plate. Nile Red and DAPI fluorescence was read on a plate reader, at excitation/emission of 485nm/535nm for Nile Red, and excitation/emission of 340nm/488nm for DAPI. Nile Red signal was normalized to DAPI signal for each well, to correct for cell number. Finally, the Nile Red/DAPI ratio was normalized to the mean of the CON CAF condition for each experiment.

#### Acquisition and Use of Human Tissue

The human tissues used in this study are the same used in the study by Francescone et al. 2021 (29), and were collected using HIPAA-approved protocols and exemption approval of the FCCC’s Institutional Review Board. Patients signed a written informed consent agreeing to donate surgical specimens to research. All samples were classified, coded and de-identified prior to distribution by the Institutional Biosample Repository Facility. Normal pancreatic tissue was obtained through US Biomax Inc. following ethical standards for collecting tissue from healthy expired donors.

#### Tumor microarray (TMA)

##### Acquisition of the TMAs

The FCCC TMAs were generated by the Biosample Repository at FCCC, totaling 80 patients (29). The MDA TMA was generated at The University of Texas MD Anderson Cancer Center, totaling 153 patient samples. The unstained slides from the TMA blocks were obtained. Tumor grading and staging were performed following the guidelines of the American Joint Committee on Cancer (AJCC), 8^th^ edition. The clinical parameters for each TMA can be found in Francescone et al. 2021 (29). The use of human tissue was approved by FCCC and University of Texas MD Anderson Cancer Center.

##### Immunohistochemistry (IHC)

The staining for NGL-1 was performed as described by Francescone et al. 2021 (29). Briefly, the sections were deparaffinized in xylene, hydrated in decreasing grades of alcohol and quenched in 3% H_2_O_2._ After, the slides were subjected to heat-induced epitope retrieval in citrate buffer (pH 6.0) for 1 hour, then permeabilized in Triton X-100 for 20 minutes, and blocked for 1 hour (1% BSA, 2% normal goat sera, or 0.2% cold water fish skin gelatin in PBS). The sections were incubated with anti-NGL-1 antibody (GTX) at 1:100 dilution, overnight at 4°C. After, the sections were washed and incubated with anti-rabbit HRP (Vector Labs), stained with DAB (Vector Labs) for 2.5 minutes and counterstained in Mayer’s Hematoxylin. The slides were then dehydrated, clarified in toluene and mounted in Cytoseal-60.

##### Scoring of TMA

The expression of NGL-1 was blindly scored by two pathologists (Dr. Cai and Dr. Klein-Szanto) for the expression of NGL-1 in fibroblasts and in immune cells on a scale of 0 to 4, based on staining intensity and percentage of positive cells (29). Two representative tumor cores were stained and scored per marker per patient, and these scores were average to generate one score per patient. All the scores and overall survival data are available in Supplementary Table 1.

##### Kaplan-Meier (KM) Plots

The KM plots were generated in the software GraphPad Prism 7, correlating overall survival (OS) with the scores from the IHC staining for NGL-1 in fibroblasts and immune cells. Long-rank test was used to examine the statistical difference in survival among different groups.

#### Simultaneous Multichannel Immunofluorescence (SMI) and image analysis

##### FFPE tissue processing

We followed reported SMI protocols (12,29) to process human and murine pancreas tissue specimens for immunofluorescence (IF). Briefly, tissue sections were deparaffinized and rehydrated before subjecting them to heat-induced antigen retrieval using a universal epitope recovery buffer (Electron Microscopy Sciences, Cat. No. 62719-20), followed by a permeabilization step with PBS-TritonX-100 (0.5%). To prevent nonspecific antibody binding, tissues were treated with an ultra-pure casein-based solution (Electron Microscopy Sciences, Cat. No. 62711) supplemented with 1% donkey serum prior to primary antibody incubation.

We conducted a two-step immunofluorescence protocol that began with an indirect IF, followed by Qdot (Q)-labeled primary antibody direct IF, and concluded with nuclear counterstaining. We previously described the methodology for Q-labeling of antibodies (12,29), and key reagents used in this procedure are detailed in the Key Resources Table.

##### Imaging of fluorescently labeled specimens

Human and murine pancreata labeled with fluorescent probes were imaged with a Caliper multispectral imaging system (LifeSciences, PerkinElmer), Nuance-FX, equipped with a Tunable Liquid Crystal imaging module and a 40X Pan Fluor 40x/0.75 Ph2 DLL lens. Spectral libraries were built using control pancreatic tissues labeled with individual fluorophores and unstained samples to account for autofluorescence. Excitation was achieved using a high-intensity mercury lamp, with different emission-excitation filter combinations (DAPI, FITC, TRITC, CY5), and scanning was conducted from infrared to ultraviolet to avoid energy transfer and excitation of nearby fluorophores. Emission for Qdot-labeled markers was collected using the DAPI filter, combined with conventional FITC, TRITC, and CY5 filters. Alternatively, a Leica SP8 DIVE confocal/multiphoton microscope system (Leica Microsystems, Inc., Mannheim, Germany) with a 63X HC PL APO 63x/1.4 oil CS2 - oil immersion lens was also used. OPSL 638, 552, and 488 lasers were used to excite DRAQ5, Cy3, and AF488 molecules, respectively. The corresponding emitted fluorescence was detected by calibrated detectors within the ranges of 700-760, 565-625, and 449-540 nm. Additionally, Q-labeled tissues were excited with an 850 nm multi-photon wavelength using an IR laser Chameleon Vision II (Coherent Inc., Santa Clara, CA). Non-descanned detectors were set up to collect signals from Q655, Q625 and Q565 within the ranges of 655-670, 625-640 and 565-580 nm respectively. To capture the images, the Leica Application Suite X 3.5.5 SW (SCR_013673) was used to scan 4 regions of interest, each consisting of 4 imaged fields, for sequential fluorescent signal collection. Identical settings were used for each ROI, and monochromatic, 16-bit image stacks corresponding to a 5 μm scan depth (Z axis displacement) were obtained.

##### SMI images analysis

Using the batch processing function of Photoshop 24.2.0 software (Adobe Inc., San Jose, CA, SCR_014199), we first standardized levels from positive signal pixels identically for each marker detected in control tissues and applied them to all respective 16-bit files, prior to transforming them into 8-bit images. We then segmented the images into “tumoral cells” or “stromal cells” compartments using MetaMorph 7.8.0.0 software (Molecular Devices Downingtown, PA, SCR_002368). We incorporated the software’s Cell Scoring function into an automated pipeline for batch processing of epithelial markers (e.g., pan-cytokeratins and amylase cocktail, positive) in conjunction with corresponding total nuclei images. The pipeline outputs correspond to unique digitally segmented masks corresponding to the tumor compartment, containing cytoplasms and nuclei sub-masks (dark and light magenta tones, respectively shown in Sup. Fig. 3I). Additionally, we obtained nuclei masks from the non-tumoral (e.g., stromal) compartment (shown as cyan masks in the figures).

Later, we incorporated these masks into our previously reported analysis pipeline using the state-of-the-art algorithm, SMIA-CUKIE (SCR_014795) (12). We scrutinized the occurrence of distinct markers in selected stromal sub-compartments (e.g., p-Smad2 levels restricted to the non-tumoral nuclei compartment, etc.), and plotted the integrated intensity values (fluorescent signal divided by compartment mask area) in corresponding graphs, expressed in arbitrary units.

#### SDS-PAGE Electrophoresis and Western Blot

##### Lysate Preparation

CAFs were plated at a cell density of 500,000 cells in 6 well plates and cultured in the presence of complete DMEM supplemented with 50 µg/mL L-ascorbic acid for 5 days. Then media was replaced with serum/glutamine free DMEM for 2 days, after which conditioned media was obtained and stored for future use and cells + CDM were washed three times with PBS. Next, 150 µL of standard RIPA buffer was added to each well and cells + CDM were removed from the wells with a cell scraper. Cells + CDM were homogenized by sonication, and lysates were spun down at 15,000 rpm for 15 minutes to pellet cellular debris. The supernatant was collected and stored at -80°C until use. For immune cells, at least 100,000 cells were isolated from mice (spleen), treated with red blood lysis buffer, washed in 10 mL PBS and then resuspended in 100 µL of RIPA buffer. Lysate was prepared as described above.

##### SDS-PAGE Electrophoresis

Prior to electrophoresis, protein concentration in the lysates was determined by the Bradford (Bio-Rad) or BCA assay (Invitrogen). Then, lysates containing 30 μg of protein were diluted in 2x loading buffer (Bio-Rad) and were loaded onto 4% to 20% polyacrylamide gels (Bio-Rad) or diluted in 4x LDS Sample Buffer (Invitrogen) and loaded onto 4 to 12%, Bis-Tris gels (Invitrogen). The gels were run and transferred to PVDF membranes using either the Bio-Rad or Invitrogen reagents and systems (Key Resources Table). Membranes were blocked in 5% nonfat milk in 0.1% Tween in tris-buffered saline (TBST) for 1 hour at RT, and primary antibodies were incubated with the membranes overnight at 4°C. Next, membranes were washed 5 times, 10 min per wash, with 0.1% TBST, and then secondary antibodies linked to horseradish peroxidase (HRP) were diluted in 5% nonfat milk in 0.1% TBST and incubated for 2 hours at RT. Membranes were washed 5x in 0.1% TBST, 10 min per wash, and incubated for 5 minutes in Immobilon Western Chemiluminescent HRP substrate (Millipore) or in SuperSignal West Pico Plus Chemiluminescent substrate (ThermoFisher). Membranes were developed using film and scanned to create digital images or using the ChemiDoc Imaging System (Bio-Rad). All the antibodies are listed in the Key Resources Table.

#### PDAC Survival assays

PDAC cells alone (RFP^+^, 20,000 cells/well) or together with CON or NGL-1 KD CAFs (GFP^+^, 20,000 cells/well) were seeded in the decellularized CDM (CON or NGL-1 KD CDM, as indicated) produced in the 24 well plates. Cells were cultured for 96 hours under nutrient depleted conditions (DMEM glutamine/FBS free media). Then, 10-15 images were captured of RFP^+^ cells and/or GFP^+^ CAFs using a Nikon A1 confocal system equipped with a 10x (plan Apo) non-oil objective. Dead PDAC cells were excluded by cell size (29). Images were batch processed as previously described by our group (29).

#### Enzyme-linked immunosorbent assay (ELISAS)

ELISAS for human TGF-β, IL-6, IL-8, GM-CSF, and mouse TGF-β, TNFα, GM-CSF, IL-6, and IL-10 were performed following the manufacturers’ instructions. Buffers were prepared using the DuoSet ELISA Ancillary Reagent kit 1 and 2. Standards, CM from CAFs, BM derived macrophages or from tumor explants were incubated for 2 hours in plates pre-coated with capture antibody. Samples were either diluted or probed without dilution, and in all cases, media alone was probed as controls (Blank). For the TGF-β ELISAS, the samples were pre-activated using the Sample Activation Kit 1 before incubation. The plates were read using Spark Multimode (Tecan) or SpectraMax ID3 Microplate Reader (Molecular Devices). The concentration of proteins in each sample was analyzed by interpolating the values using a standard curve generated by dilutions of the standard (provided with each kit) and correcting with the values of the blanks for each sample. The corrected concentrations were normalized to total protein content (for CAFs and immune cells) or to the weight of each pancreas fragment (tumor explants).

#### Multiplex Assay (multiplex ELISA)

The multiplex assays for the detection of multiple cytokines were carried out as recommended by the manufacturers (Millipore and Meso Scale Discoveries) and performed in the FCCC High Throughput Screening Facility. Briefly, standards, calibrator and experimental samples (CM from CON and NGL-1 KD CAFs, CM from murine tumor explants) were added (10 μL/well) to pre-coated plates containing beads bounded to analyte-specific capture antibodies and incubated overnight. Detection of cytokines in the CM was measured according to the manufacturer’s instructions, using the cytokine detection magnetic bead assay kit (MILLIPLEX MAP Mouse 32-plex cytokine/chemokine panel, MilliporeSigma and U-Plex, Meso Scale Discoveries). Standard curves were generated utilizing the kit’s standards and analyzed in the BioPlex 200 biomarker analyzer (Bio-Rad, Hercules, CA) or MSD SECTOR Imager 2400 (Meso Scale Discoveries), and the data were processed using the BioPlex manager software or the MSD Workbench 4.0 software (Meso Scale Discoveries), applying 5 parameter logistic regression to provide the concentration of cytokines in pg/mL.

#### Reverse Transcription and Quantitative Polymerase Chain Reaction (q-PCR)

Total RNA was extracted using the PureLink^TM^ RNA mini kit according to the manufacturer’s instructions (ThermoFisher). The RNA quality was evaluated using a Bioanalyzer and contaminating genomic DNA was removed using the Turbo DNA-free kit (ThermoFisher). RNA concentration was determined by spectrophotometry (NanoDrop 2000, ThermoFisher). Reverse transcription reactions were performed using the High-Capacity cDNA RT Kit (ThermoFisher), as directed by the manufacturer’s instructions. qRT-PCR reactions were done in barcoded 96-well plates optimized for use in the StepOnePlus Real Time-PCR system (Applied Biosystems). qRT-PCR reactions contained the following components: Power SYBR Green PCR master mix (Applied Biosystems), gene-specific primers, and template DNA. The reactions were performed under the following conditions: initial denaturation at 95°C for 30 seconds and then 40 cycles of a 5 second denaturation at 95°C, 15 second annealing at 60°C, and a 10 second extension step at 72°C. Standard melt curves were also collected to confirm primer efficiency. All primer sequences were obtained from PrimerBank (202) and are available in the Key Resources Table. Relative mRNA expression for each gene was calculated by the ΔΔCt method, using *Rn18s* as a house keeping control. These data were then normalized to immune cells isolated from the spleens of naïve (non-tumor bearing) mice to determine fold-change differences in gene expression.

#### Human T cell isolation and activation assay

##### Peripheral blood sampling and CD8+ T cell isolation

Four healthy donors participated in the study. Written consent was obtained prior to obtaining blood samples. The study was conducted according to the ethical standards of the Helsinki Declaration and approved by Fox Chase Cancer Center institutional review board. Peripheral blood samples (15-20 mL) were drawn into Vacutainer Sodium Heparin plastic tubes. Peripheral blood mononuclear cells (PBMC) were isolated by a standard method of centrifugation using Lymphoprep medium density gradient. The interface was collected and adjusted to 50,000,00 cells/mL in Ca^2+^/Mg^2+-^free Dulbecco’s phosphate-buffered saline (DPBS). Further steps were performed according to the recommendations of the EasySep Human CD8^+^ T Cell Isolation Kit manufacturer. Briefly, the isolation cocktail (50 µL/mL) was added to PBMC suspension, then incubated for 5 minutes at RT. After the incubation, the RapidSpheres reagent (50 µL/mL) was added, cells were diluted up to 2.5 mL with Ca^2+^/Mg^2+^ free DPBS and put on the EasySep Magnet. After 3 minutes of incubation, the cell suspension was transferred to a new tube. The purity of the CD8^+^ T cell suspension reached 93%. All reagents were from STEMCELL Technologies.

##### IFN-γ Secretion Assay

To stimulate IFN-γ response, CD8^+^ T cells (200,00 cells/well) were incubated for 20 hours in 5% CO_2_ at 37°С in various stimulatory conditions: cells were resuspended in 200 µL DMEM media containing 25% donor’s serum, with or without CAF media (CON and NGL-1 KD, 5, 10, or 20%) in the presence of 5 µL DynaBeads Human T-activator (Gibco) and 1000 U/mL recombinant human IL-2. Cells with no DynaBeads stimulation were used as a negative control, and cells with stimulation but without CAF CM were used as positive control. After the activation, cells were transferred to 5 mL round bottom polystyrene FACS tubes, washed, and placed on the EasySep Magnet for 1 minute to remove leftover DynaBeads. Catching anti-CD45/anti-IFN-γ antibodies (10 μL, IFN-γ Catch Reagent, Miltenyi) were added to the tubes with activated CD8^+^ T cells and cultured 10 minutes on ice. To initiate IFN-γ secretion, 3 mL of warm RPMI-1640 medium was added to each tube, then tubes were placed on an automatic shaker for 45 minutes in 5% CO_2_ at 37°C. Cells were washed and stained with CD3-FITC, CD8-Pacific Blue, and IFN-γ-PE (Detection antibody, Miltenyi). Finally, cells were washed again with a buffer containing propidium iodide (100 ng/mL). Analysis was performed on BD Aria II flow cytometer using BD FACS DiVa Software 8.0.1. The .fcs files were then analyzed with FlowJo v10 Software.

#### Peptide design

The crystal structure of a complex of NetG1 and NGL-1 (PDB: 3ZYJ (111)) was investigated for the identification of peptides from NetG1 that might independently bind to NGL-1. Four surface loops of NetG1 are in contact with the surface of NGL-1. Three of these have previously been labeled the NGL discriminant loops I-III of netrin G family proteins (111) (residues 78-91, 209-219, and 270-285 of human NetG1, respectively; numbering according to UniProt entry Q9Y2I2; sequence numbering in PDB: 3ZYJ differs). In addition, the loop 179-190 also contacts NGL-1 in the crystal structure. AlphaFold-Multimer was utilized to investigate the complex NetG1 and NGL-1, since the crystal structure had none of the expected disulfide bonds predicted to form in netrin G and NGL family proteins. AlphaFold-Multimer formed 25 disulfides in the complex (20 in NetG1 and 5 in NGL-1). The top-ranked model had an RMSD of 1.2 Å over 596 amino acids to the complex in PDB: 3ZYJ. We focused on NetG1 discriminant loop I, residues 78-91 with sequence TFCAMGNPYMCNNE for peptide design. We chose a short fragment of this peptide with sequence NPYMCNNE, which forms direct contacts (within 5 Å) with 8 of the 10 leucine-rich repeats of NGL-1, comprising 19 residues (K59, L80, N82, H84, T104, Q106, T128, E130, F132, E152, W154, R156, R176, D178, Y201, N203, E223, K247, W248). Since the two cysteines in the longer fragment form a disulfide bond, we mutated the Cys residue in the shorter fragment to serine and synthesized the peptide NPYMSNNE.

#### Binding assay (Sandwich ELISA)

For this binding assay by sandwich ELISA, wells in a 96 well plate were coated overnight with either anti-NGL-1 Peptide (1 mM), recombinant NetG1 (50 nM), PBS or BSA (50 nM) as control. The next day, wells were washed with ELISA wash buffer (R&D Biotechne), and then blocked with 1% BSA for 1h at room temperature. Afterwards, the wells were treated with recombinant NGL-1 (50 nM), and binding was allowed to occur for 2h at room temperature. Next, after washing, wells were treated with an antibody against NGL-1 (5 µg/mL, Genetex) for 1h, and then an antibody against rabbit antibodies, linked to HRP, was added. Finally, TMB substrate was added, and the absorbance (450 nm) was a direct read-out of the amount of NGL-1 bound to the well.

#### Mice and animal care

C57BL/6J (stock number 000664, CD45.2 mice) and the B6.SJL-*Ptprc^a^Pepc^b^*/BoyJ (stock number 002014; CD45.1 mice) mice were purchased from Jackson Laboratory and maintained at the FCCC Animal Facility. Other strains are described below. All of the mice used in this study were maintained in the AAALAC-approved Laboratory Animal Facility at Fox Chase Cancer Center (FCCC, Philadelphia, PA) under controlled conditions (22 ± 3°C for 12 h light/dark cycle, free access to food and water). Animal care, research and animal euthanasia protocols were in accordance with the principles and guidelines adopted by Report of the American Veterinary Medical Association Panel on Euthanasia and approved by the FCCC Institutional animals care and use committee (IACUC). For every experiment, we used equal numbers of male and female mice.

#### NGL-1 Knockout (KO) mice

Embryos of NGL-1 KO mice were kindly donated by Dr. Shigeyoshi Itohara from the RIKEN Brain Science Institute in Japan and rederived in Laboratory Animal Facility at Fox Chase Cancer Center. The NGL-1 knockout mice were generated by homologous recombination using C57BL/6-derived embryonic stem cells; to disrupt the translation of viable protein, the translation initiation codon in the third exon was replaced with a noncoding set of nucleotides, including a *loxP* site (39). The mice were maintained in the C57BL/6 background and crossed as heterozygotes, to generate wild-type, heterozygous and knockout mice. Mice genotype was obtained by PCR (Taq polymerase, Thermo Fisher), using the following primers:

NGL-1 WTf GATGACCTTACATCCACAGC NGL-1WTr GACGATTGTCAAAGAGTTCC NGL-1KOf CAGTGGCAGGATTATACACC NGL-1KOr GCCGCCCTTTAGTGAGGGTT

PCR reaction conditions were as follows:

5 μL of isolated tail or ear DNA (proteinase K, ethanol extraction method) added to a reaction mixture containing 0.5 µL TAQ polymerase, 5 μL of a mixture of all 4 primers (0.5 µM final concentration), 4 μL MgCl_2_ (2 mM final concentration), 5 μL dNTPs (0.2 mM final concentration, and 30 μL nuclease free water. Cycling conditions: 95°C for 2 min, then 40 cycles of 95°C for 30 seconds, 50°C for 30 seconds, 72°C for 1 minute, and a final extension after 40 cycles of 5 minutes at 72°C. WT band = 395 bp, KO band = 805 bp.

#### Spontaneous pancreatic cancer models

We analyzed bone marrow, spleen and pancreas tissue from the KC (Pdx1-Cre; Kras^LSL-G12D/+^) and KPC (Kras^LSL-G12D/+^; p53^flox/WT^; Pdx-Cre/^+^) mice at different ages: 2-12 months old for KC animals and 15-28 weeks for KPC animals (37,38). Both mouse models were provided by Dr. Kerry Campbell (FCCC, Philadelphia, PA).

#### Orthotopic allografts

Orthotopic injections were performed as in Kim et al. (203), with modifications. We used C57BL/6 mice, WT, Het or KO for NGL-1 as hosts, 8-16 weeks old (younger cohorts) or 12-20 months old (older cohorts). Mice of both sexes were equally distributed. During the surgery, all animals were kept under anesthesia with isoflurane (1 – 3%) and 1 L/min oxygen and received analgesic (Buprenorphine, 0.1 mg/mL, subcutaneously) and local anesthetics (Bupivacaine, 2 mg/kg, subcutaneously). Prior to the incision, the animals were shaved in the site of the surgery and disinfected. A small incision was made, and the spleen was gently grasped to expose the pancreas. Then, 200,000 KPC3 cells were injected (in 30 μL of PBS) orthotopically in the tail of the pancreas. The incision was stitched, and the animals were observed until recovered from anesthesia. All the animals received analgesics for 2 additional days and were monitored for signs of disease weekly. After 2.5-3.5 week, the animals were euthanized and the spleen, bone marrow and pancreas were removed for further analysis. Each pancreas was weighed, a small fragment was kept for conditioned media generation and the rest was kept for histological analysis. In some cases, the pancreas was digested to obtain a single cell suspension (29,204).

#### *In vivo* treatments

In some experiments, mice were treated with FOLFIRINOX (from the FCCC pharmacy) twice a week, intraperitoneally, after 5 days of the tumor cells injection. Doses were: Leucovorin Calcium (50 mg/kg), 5-Fluorouracil (25 mg/Kg), Irinotecan hydrochloride (25 mg/kg) and Oxaliplatin (2.5 mg/kg), in PBS (205). In other experiments, mice were treated with the anti-NGL-1 peptide (250 μM/mouse, in PBS), intraperitoneally, every day after 3 days of the tumor cells injection. In all cases, control groups received PBS in the same doses and regimens.

#### Murine histopathological analysis

Murine tissue (pancreas, liver, lung, kidney, spleen) were fixed in 10% phosphate-buffered formaldehyde (formalin) for 24 to 48 hours, dehydrated (70% ethanol 3 hours, 95% ethanol 2 hours, 100% ethanol 2 hours, 100% ethanol xylene mixed 1 hour, xylene 3 hours), then immersed in paraffin. Sections of 5 μm were mounted on microscope slides and stained with Hematoxylin and Eosin (H&E) for general characterization or proceeded to immunohistochemistry for specific markers. All slides were visualized in a Nikon Eclipse 50i microscope and photomicrographs were taken with an attached Nikon DS-Fi1 camera.

##### Tumor characterization

H&E slides from pancreatic tumors were scanned using the Aperio ScanScope CS2 Scanner (Leica) and the images were visualized with ImageScope (Aperio). The areas of interest were outlined and characterized blindly by a pathologist (Dr. Cai). Analysis included tumor differentiation (well, moderately and poorly differentiated) and lesion stage (ADM, PanINs and PDAC).

##### TUNEL staining

Dead cells were quantified using the DeadEnd Fluorometric TUNEL System (Promega). Briefly, the tissue was deparaffinized and fixed in 4% formaldehyde for 15 minutes at room temperature. The slides were then washed twice in PBS and treated with 20 μg/mL proteinase K solution for 10 minutes at room temperature. Following two washes with PBS, the slides were fixed again with 4% formaldehyde for 5 minutes at room temperature and then submerged in equilibration buffer for 5 minutes. Slides were then labeled with TdT reaction mixture for 1 hour at 37°C and the reaction was stopped by adding SSC buffer for 15 minutes at room temperature. The slides were washed three times with PBS and then stained with DAPI (1:10,000) for 15 minutes at room temperature. After one wash with PBS, the slides were mounted and imaged using the Eclipse Ti2-E Inverted Microscope. Metamorph software was used to quantify dead cells (green) in total nuclei count (blue) in each field of view. Ten images per slide were acquired at 20X.

##### Immunohistochemistry (IHC)

For all IHC stainings, formalin-fixed and paraffin embedded (FFPE) sections were deparaffinized using EZ Prep solution for 16 min at 72 °C. Epitope retrieval was accomplished with CC1 solution at high temperature (95 – 100 °C) for 32 or 64 minutes. Primary antibodies were tittered with a TBS antibody diluent into dispensers on an automated stainer (Ventana, Roche). As negative control, the primary antibody was replaced with rabbit, mouse or rat IgG to confirm specificity. Immune complexes were detected using the Ventana OmniMap anti-rabbit, anti-mouse or anti-rat detection kits and developed using the Ventana ChromMap DAB detection kit, according to the manufacturer’s instructions. Slides were then counterstained with hematoxylin II (Roche) for 8 minutes followed by Bluing reagent for 4 minutes. The slides were then dehydrated in ethanol series, cleared in xylene and mounted. All slides were viewed with Nikon Eclipse 50i microscope and photomicrographs were taken with an attached Nikon DS-Fi1 camera. Slides were scanned using the Aperio ScanScope CS 5 Scanner (Leica Aperio) and the images were visualized with ImageScope (Aperio). The areas of interest were outlined by a pathologist (Dr. K.Q. Cai). Quantification of positive areas was performed using Aperio (Leica).

The following primary antibodies and respective conditions were used: anti-NGL-1 (Rabbit, 1:150, Genetex, antigen retrieval EDTA 32 minutes), anti-CD4 (Rabbit, 1:100, Cell Signaling, antigen retrieval EDTA 64 minutes), anti-CD8 (Rabbit, 1:50, Cell Signaling, antigen retrieval EDTA 64 minutes), anti-F4/80 (Rat, 1:100, Bio-Rad, antigen retrieval citrate 32 minutes), anti-FoxP3 (Rabbit, 1:20, Cell Signaling, antigen retrieval EDTA 64 minutes), anti-NK1.1 (Mouse, 1:50, Invitrogen, antigen retrieval EDTA 32 minutes), anti-pSmad2 (Rabbit, 1:300, Cell Signaling, antigen retrieval EDTA 64 minutes), anti-Ki67 (Rabbit, 1:500, Cell Signaling, antigen retrieval EDTA 32 minutes), anti-cleaved caspase 3 (Rabbit, 1:500, Cell Signaling, antigen retrieval EDTA 32 minutes).

#### Second Harmonic Generation (SHG) of Polarized Light Microscopy

Slides stained with H&E and containing murine pancreata from the different groups were evaluated by a pathologist (Dr. Cai) to identify areas containing tumoral and normal tissue. Random regions of interest were defined for downstream analysis (12-18 images per mouse), from which the relative levels of SHG emitted by fibrillary collagen were determined. Polymerized and fibrillar collagen fibers were analyzed in images collected by SHG of polarized light from FFPE mounted glass slides. Briefly, a water-immersion 25X HC FLUOTAR L 25x/0.95NA W VISIR objective mounted in a Leica SP8 DIVE confocal/multiphoton microscope system was used to image the tissue, which was excited with an 850 nm wavelength generated by multiphoton IR laser Chameleon Vision II. Stromal areas were examined and a non-descanned detector was used to capture the backward SHG emission (signal between 410-440 nm wavelength) from three ROI, using the Leica Application Suite X 3.5.5. software, each containing 4-6 imaging fields, for automatic SHG signal collection. The settings were identical for each ROI, and monochromatic, 16-bit image stacks with a depth of 5 μm (Z total distance) were recorded as results. Additionally, H&E monochromatic images corresponding to each ROI were mapped by illuminating the stained tissue with a 488nm laser and collecting the signal with a transmitted light detector (27,206).

The stacked images were reconstituted as maximum projections using the software FIJI (ImageJ 1.52p). Initially, we reconstructed raw tri-dimensional (x, y, z) image stack files into maximal projected two-dimensional (x, y) 16-bit images. Identical signal-to-noise thresholds were applied to all images to select SHG positive-signal pixels. We generated digital masks using corresponding H&E images, which were then used to calculate the SHG integrated intensity (i.e., SHG signal divided by tissue area). Mean integrated intensity values were normalized with the control group (either WT or WT/WT depending of the set of experiments). Results represent arbitrary units compared to control groups.

#### BaseScope

Target probe oligonucleotides (Lrrc4c) were designed (double Z design strategy) by Advanced Cell Diagnostics (Hayward, CA) and manufactured by Eurofins MWG Operon (Huntsville, AL). A single color BaseScope assay (Advanced Cell Diagnostics, Newark, CA) for the detection of NGL-1 was performed using the BaseScope protocol on a Roche Ventana Discovery Ultra (Roche Ventana Medical Systems, Tucson, AZ). Briefly, sections were baked (32 min at 60°C) and deparaffinized on the instrument, followed by target retrieval (24 min at 97°C for tissues) and protease treatment (16 min at 37°C). Probes were then hybridized for 2 h at 42°C followed by BaseScope amplification and chromogenic detection using BaseScope 2.5 VS Red detection reagents (Fast Red with alkaline phosphatase). The following RNAscope probes were used in this study: dapB (bacterial gene; negative control), Ubc (housekeeping gene; positive control), Lrrc4c-1zz-st and Lrrc4c-1zz-st1.

#### Blood collection and processing

For all experiments that required analysis of the blood, 200 μL of blood from the submandibular vein was collected from each animal using a 4 mm lancet (Goldenrod), in 1.5 mL of RPMI medium plus heparin (Sigma). The mixture was carefully layered on top of 1 mL of Lympholyte M (Cedarlane) at room temperature in 5 mL polystyrene tubes. The samples were centrifuged at 2600 rpm for 20 minutes at room temperature. Lymphocytes (visible ring of cells) were then harvested from the interface between Lympholyte M and medium, placed in another 5 mL polystyrene tube and washed twice with 3 mL staining buffer (PBS Ca2+ and Mg2+ free, 5mM EDTA, 5% heat inactivated FBS) and centrifuged for 7 min at 1800 rpm. After washes, the red blood cells were lysed with ACK Lysing buffer (KD Medical) for 5 min, washed again in staining buffer and cells proceeded to staining for flow cytometry analysis.

#### Bone marrow chimera

For the chimerization, we used the following male and female mice, 8-16 weeks old: WT C57BL/6J (B6 CD45.2), B6.SJL-*Ptprc^a^Pepc^b^*/BoyJ (B6 CD45.1) from The Jackson laboratory (stock number 002014) and NGL-1 KO mice (CD45.2).

##### Donors

Donor animals (WT C57BL/6J, B6 CD45.1 or NGL-1 KO mice, were euthanized by cervical dislocation. The bone marrows were removed in a sterile environment, and the bone marrow cells were flushed into single cell suspensions with sterile PBS. The final concentration of cells was 5 million bone marrow cells in 100μL of PBS.

##### Recipient mice, irradiation, and transplants

Recipient animals were WT C57BL/6J (CD45.2), B6 CD45.1 or NGL-1 KO mice and received antibiotics in drinking water (6.4mg/mL Polymyxin B Sulfate and 250mg/mL Neomycin) one day before the irradiation. Animals were lethally irradiated (total of 1100 rad) through 2 rounds of irradiations: 600 rad in the first round and 500 rad after 3-5 hours from the first round of irradiation. After 24 hours of irradiation, the recipient animals received the donor cells intravenously via tail vein injection, under laminar flow hood. Following the bone marrow transfer, these animals were kept on antibiotics in drinking water for another 4 weeks. For the tail vein injections, each mouse was placed under a heat lamp for 2-5 minutes, then restrained, the tail vein skin disinfected with ethanol 70%, and injected with 100 µl of PBS containing 5 million cells, using sterile 27 G needle attached to a 1 mL syringe. Gauze was applied on the bleeding site for approximately 30 seconds to achieve hemostasis.

##### Assessment of bone marrow reconstitution

10 weeks after bone marrow cell transplantation, blood from all animals was collected from the submandibular vein as described above and stained for CD45.1 (APC), CD45.2 (FITC), CD3 (PE) and CD11b (APC-Cy7).

##### Orthotopic injection of murine PDAC cells

12 weeks after bone marrow cell transplantation, PDAC cells were injected in the pancreas as described above. Final groups were: host WT/immune cells WT (1 male and 4 females), host WT/immune cells NGL-1 KO (2 males and 3 females) and host NGL-1 KO/immune cells WT (4 males and 1 female).

#### Generation of conditioned media from tumor explants

Conditioned media from tumor explants was generated by culturing a fragment of the pancreatic tumor from the murine models (the fragment is weighed prior culture) in 2 mL of RPMI medium + 1% P/S (10,000 U/mL) for 24 hours at 37°C. Afterwards, the conditioned medium is centrifuged at 2,000 rpm for 5 min to eliminate cells, transferred to a clean 2 mL tube and frozen at 80°C until analyses (29).

#### Single cell suspensions from pancreatic tumors

Approximately half of the pancreas (longitudinal cut) was digested to obtain a single cell suspension. Briefly, the tissue was cut into small fine pieces (2-3 mm) using sterile blades. The pieces were transferred to gentleMACS C tubes (Miltenyi Biotec) containing 2.35 mL of DMEM with 1% P/S and tumor dissociation enzymatic cocktail (100 μL of enzyme D, 50 µL of enzyme R and 12.5 of enzyme A, Miltenyi Biotec). The suspension was dissociated in a gentleMACS Dissociator (Miltenyi Biotec) (program m_impTumor_02) and incubated at 37°C under continuous rotation for 40 minutes. Samples proceeded to a second round of dissociation (twice in program m_impTumor_03), then strained through a 70 μm strainer. The suspension was washed with 10 mL of RPMI + 10% heat inactivated FBS and centrifuged at 1,500 rpm for 7 minutes. Afterwards, red blood cells were lysed with ACK Lysing buffer for 5 min, washed again in staining buffer and cells proceeded to staining for flow cytometry analysis. For single cell RNA sequencing analysis, the suspensions were subjected to an extra step for dead cell removal using (Dead Cell Removal Kit, Miltenyi Biotec) (207).

#### Single cell suspensions from spleens and bone marrows

To obtain single cell suspensions from the spleen, spleens were removed and placed in 70 μm strainers. The bottom of a 1 mL syringe (sterile) was used to smash the spleen towards the strainer, and the suspension was washed in 10 mL of staining buffer. Similarly, bone marrow cells from the tibias were flushed with staining buffer using a 1 mL syringe coupled to a 25G needle. The cells were strained through a 70 µm strainer and washed in 10 mL of staining buffer. Cells suspensions (from spleen or bone marrow) were centrifuged at 1500 rpm for 5 minutes and proceeded to red blood cell lysis with ACK Lysing buffer for 5 min, washed again in staining buffer and cells were stained for the subsequent flow cytometry analysis or used for downstream assays.

#### Immunophenotyping and cell sorting by Flow cytometry

Single cell suspensions were incubated with anti-mouse CD16/CD32 (Fc Block, BD Biosciences) for 12 minutes and surface staining was performed at 4°C for 20 minutes. Cells were run on LSRII flow cytometer (BD Biosciences) for immunophenotyping or in a FACSAria II flow cytometer (BD Biosciences) for cell sorting. All the data was analyzed in FlowJo (FlowJo, LLC).

*Antibodies*: Every staining was performed with 1:100 or 1:200 monoclonal anti-mouse antibodies in staining buffer. Antibodies used can be found in the Key Resources Table. Dead cells were excluded with Sytox Blue, 1:500.

*Sorting strategy*: To obtain CD4 T cells, CD8 T cells and NK cells, isolated cells were gated in the following manner: singlets, live cells, CD3^+^ T cells or CD90.2^+^ (Thy1) cells, then CD4^+^ T cells and CD8^+^ T cells, or CD3 negative T cells, then NK1.1^+^ cells for NK cells). To obtain neutrophils and monocytes, isolated cells were gated as follows: singlets, live cells, CD11b^+^ cells, then Ly6G^+^ for neutrophils and Ly6C^high^ for monocytes. To obtain macrophages, the following gating strategy was used: singlets, live cells, CD11b^+^ cells, then Ly6C ^low/negative^ and Ly6G ^negative^, then F4/80^+^ cells. Cells were sorted in complete medium (RPMI medium + 10% heat inactivated FBS+ 1% P/S (10,000 U/mL) 50 mM 2-Mercaptoethanol or directly in Trizol (Thermo Fisher) or RNA lysis buffer (Thermo Fisher) for subsequent RNA extraction.

#### *In vitro* murine T cell stimulation and proliferation assay

Sorted CD4^+^ or CD8^+^ T cells or total splenocytes from WT and/or NGL-1 KO mice were counted and 10 million cells were stained with 1 μM of Cell Trace – CFSE (Invitrogen) in 10 mL of Ca^2+^ Mg^2+^ Free PBS for 10 minutes at room temperature. The tube was carefully homogenized two times during these 10 minutes. Then, the CFSE was carefully (drop by drop) quenched with heat inactivated FBS (1.5%) and the suspension was centrifuged at 1,500 rpm for 5 minutes. The pellet was washed again in complete medium, centrifuged and the cells were counted. Subsequently, 100,000 cells were plated in triplicate in U-bottom 96 wells plates (Corning) in 200 μL of complete medium alone (negative control) or in the presence of 0.5 μg/mL of anti-CD3 (BD Biosciences) and 0.25 μg/mL of anti-CD28 (BD Biosciences) (positive control) for 48 hours. After that, cells were subjected to staining for flow cytometry. Importantly, a freshly CFSE-stained and a non-stained sample were prepared on the same day as the experimental samples were prepared for flow cytometry analysis as positive and negative controls of CFSE staining, respectively. For analysis of T cell proliferation, cells were gated on singlets, then live (using the Sytox Blue dead cell exclusion), followed by CD90.2^+^ cells, then CD4^+^ and CD8^+^ cells, and finally CFSE^+^ cells, where the negative and positive gates were set using unstimulated control cells. Results were expressed as percentage of proliferation.

#### Enrichment for Pan T cells

Spleens from WT and NGL-1 KO mice were processed into single cell suspensions and red cells were lysed as described above. Using the Pan-T cell isolation Kit II (Miltenyi), the resultant cells were resuspended in staining buffer (complete PBS, pH 7.2, 0.5% BSA and 2 mM EDTA) at a concentration of 10 million cells for every 40 µL of buffer and stained at 4°C for 5 minutes with Biotin-Antibody cocktail (10 µL for every 10 million cells). Then, cells were stained with anti-biotin microbeads (20 µL of microbeads per 10 million cells, in 30 µL of buffer) for 10 minutes, at 4°C. Cells were then magnetically isolated, using the LS Columns (Miltenyi) in the magnetic field of a MACS^TM^ separator (Miltenyi) for enrichment of T cells by negative selection. Purity was assessed by flow cytometry, staining for live CD3+ T cells and confirmed to be over 90% in all cases.

#### T cell polyfunctionality analysis

The enriched CD3^+^ T cells from WT and NGL-1 KO mice were plated in triplicate in U-bottom 96 wells plates (Corning) in 200 μL of complete medium alone (negative control) or in the presence of 0.5 μg/mL of anti-CD3 (BD Biosciences) and 0.25 μg/mL of anti-CD28 (BD Biosciences) (positive control) for 48 hours. Next, cells were collected and processed following the protocol of the manufacturer (IsoPlexis/Bruker Single-Cell Adaptive Immune Chip - (M), ISOCODE-1004-4). Briefly, cells were washed with PBS and 30,000 viable cells in complete media were loaded in the IsoCode chip (Isoplexis/Bruker) and incubated for 16 hours in the IsoSpark machine at 37°C, with 5% CO_2_. Subsequently, secreted proteins from ∼1000 single T cells were captured by the 32-plex antibody barcoded chip and analyzed by backend fluorescence, at single cell resolution. The chips were from the mouse adaptive immune panel (S-PANEL-1004-4), consisting of the following cytokines: Granzyme B, IFN-γ, MIP-1α, TNF-α, GM-CSF, IL-2, IL-5, IL-7, IL-12p70, IL-15, IL-21, sCD137, CCL11, CXCL1, CXCL13, IP-10, RANTES, Fas, IL-4, IL-10, IL-13, IL-27, TGFβ1, IL-6, IL-17A, MCP-1, IL-1β. The fluorescent signals were analyzed in the IsoSpeak software, providing the following parameters: T cell polyfunctionality, as frequency of T cells that secreted two or more factors and Polyfunctional Strength Index, as percentage of polyfunctional single cells in a sample that secretes two or more proteins, multiplied by the average signal intensity of the secreted proteins from individual functional groups from each cell. The functional groups were defined by the manufacturer, and are as follows: Effector (Granzyme B, IFN-γ, MIP-1α, TNF-α), Stimulatory (GM-CSF, IL-2, IL-5, IL-7, IL-12p70, IL-15, IL-21), Chemoattractive (CCL11, CXCL13, IP-10, RANTES), Regulatory (Fas, IL-4, IL-10, IL-13, IL-27, sCD137), Inflammatory (IL-6, IL-17A, MCP-1, IL-1β) and others (CXCL-1 and TGFβ1).

#### Generation of bone marrow-derived macrophages and macrophage stimulation

Total bone marrow cells from WT and/or NGL-1 KO mice were counted and plated in 6 well plates in the presence of complete RPMI medium + 100 ng/mL of recombinant mouse m-CSF (R&D Systems). Cells were cultured for 5 days, then the medium was refreshed with extra m-CSF and cells were cultured for an extra 2 days. On day 7, macrophages were stimulated with 100 ng/mL of LPS (Sigma) in RPMI free of serum for 4 hours or with 100 ng/mL of LPS (Sigma) + 50 ng/mL of IFNγ (Peprotech) in complete medium for 24 hours. After that, the medium was changed to RPMI free of serum overnight and then the supernatants were collected, centrifuged at 1,500 rpm for detection of cytokine production by ELISA (208).

#### Single-cell RNA Sequencing

Single cell suspensions were obtained from the pancreatic tumors of WT and NGL-1 KO mice, 3 weeks after the orthotopic allografts. The tumors were minced and digested as described above, using the Mouse tumor dissociation Kit (Miltenyi Biotec) and dissociated using the GentleMacs dissociator (Miltenyi Biotec). Dead cells were removed using the Dead cell removal microbeads (Miltenyi Biotech) as described above. The single cell droplets were generated with Chromium single-cell controller, then single cells suspensions were converted to barcoded libraries using Chromium Single Cell 30 Library, Gel Bead & Multiplex Kit, and Chip Kit V3 (10X Genomics, #PN-1000092). Estimated 7,000 cells per sample were recovered to make cDNA at single-cell level. The Illumina adapters were ligated to the fragmented cDNA. The libraries quality and concentration were analyzed on the bioanalyzer with the High sensitivity DNA kit (Agilent, # 5067-4626). The scRNA libraries were sequenced on IlluminaNextseq 2000. Fastq files were generated by using the bcl2fastq wrapper of CellRanger algorithm (10X Genomics) and a minimum of 20,000 read pairs/cell were obtained. The single-cell gene expression data were processed by using CellRanger software and visualized using Loupe cell browser from 10X Genomics.

#### scRNAseq Bioinformatics

Single cell RNA sequencing data was processed as previously described (43). Briefly, FASTQ files were aligned to the mm10 reference genome and processed in Seurat (v4) (209). Each sample was normalized and scaled, and standard principal component analysis and downstream clustering was performed to define cell clusters. Lineage markers were used to define cell types. Differential gene expression comparisons were performed using FindMarkers (identifies differentially expressed genes between two groups of cells using a Wilcoxon Rank Sum test) function in Seurat. P-value adjustment was performed using Bonferroni correction based on the total number of genes in the dataset. Feature plots or Dot plots were generated to visualize specific gene expression profiles. The data generated is available at Gene Expression Omnibus (GEO) under the accession number GSEXXXXX (to be deposited).

#### Bulk RNA sequencing

##### Human fibroblasts

CON, NGL-1 KD1 and NGL-1 KD2 CAFs were seeded in 6 wells plates, (500,000 cells/4mL of complete medium), in triplicate, and allowed to produce their own ECM for 5 days as described above. Then, the complete media was replaced for glutamine/serum free media for another 2 days, and after that the cells were lysed in TRI reagent LS (Sigma) for subsequent RNA extraction.

##### Immune cells from murine pancreatic tumors

6 WT and 6 NGL-1 KO mice were orthotopically allografted with KPC3 cells as described above. After ∼3 weeks, the tumors were minced and digested as described above, using the Mouse tumor dissociation Kit (Miltenyi Biotec) and dissociated using the GentleMacs dissociator (Miltenyi Biotec). Two animals/genotype were combined to provide enough input cells for sorting, totaling 3 samples/genotype (each sample composed by 2 animals). The cells suspensions were cleared of red blood cells (using ACK buffer) and processed for staining with flow cytometry antibodies. The sorting strategy was as follows: isolated cells were gated in singlets, live cells (Sytox blue^negative^), CD45^+^, CD11b^+^, then Gr1^+^ for granulocytes and GR1^negative^ and F4/80^+^ for macrophages. Also, CD11b^negative^ and CD90.2^+^ cells labeled T cells, which were further gated as CD4^+^ or CD8^+^ T cells. Cells were sorted directly in TRI Reagent LS (Sigma) for subsequent RNA extraction.

RNA was extracted using Direct-zol RNA mini prep plus kit. Total RNA (20 – 50 ng) from each sample was used to make the library according to the NEBNext® Ultra™ Directional RNA Library Prep Kit for Illumina. Briefly, mRNAs were enriched twice via poly-T based RNA purification beads and subjected to fragmentation at 94°C for 15 min via divalent cation method. The 1^st^ strand cDNA was synthesized by reverse transcriptase and random primers, followed by 2^nd^ strand synthesis. A single ‘A’ nucleotide was added to the 3’ ends of the blunt fragments. Adapters with Illumina P5, P7 sequences as well as indices were ligated to the cDNA fragment. After SPRIselect bead (Beckman Coulter) purification, PCR reaction was used to enrich the fragments. Libraries were again purified using SPRIselect beads, submitted to a quality check on Agilent 2100 bioanalyzer (Agilent Technologies) using Agilent high sensitive DNA kit (Agilent Technologies), and quantified with Qubit 3.0 fluorometer (Invitrogen) using Qubit 1x dsDNA HS assay kit (Invitrogen). Paired end reads at 65bp were generated by using Nextseq 2000 high output reagent kit v2.5 (Illumina). Faseq files were obtained at Illumina base space (https://basespace.illumina.com). For each sample, approximately 60 million PE reads were obtained.

#### Bulk RNA sequencing Bioinformatics

Sequence reads from RNA-Seq experiments using human CAFs were aligned to indexed human HG38 genome with STAR. Sequence reads from RNA-Seq experiments using murine immune cells were aligned to indexed murine mm10 genome (210). The number of raw counts in each known gene from the RefSeq database was enumerated using htseq-count from the HTSeq package (211). Differential expression between samples and across different conditions was assessed for statistical significance using the R/Bioconductor package DESeq2 (212). Genes with a false discovery rate (FDR) ≥ 0.05 and a fold-change ≥ 1.5.

Gene ontology (GO) and pathway analysis (Human CAFs): The enriched canonical pathways and gene interaction networks of significant genes were generated using Ingenuity Pathway Analysis (QIAGEN Inc.,). Conditional hyper-geometric method implemented in GO-stats package (213) were used to identify enriched GO functional categories among genes of interest with a p-value cut off of 0.001.

Murine samples: Using the DE gene lists (FDR ≥ 0.05 and fold-change ≥ 1.5), heatmaps were built showing the differentially expressed genes between the different cell populations, where red bars represent upregulated genes and blue bars represent downregulated genes, using WT or CON cells as references. The heatmaps were created using the web tool Morpheus (https://software.broadinstitute.org/morpheus).

The gene enrichment analyses were performed using KOBAS (http://kobas.cbi.pku.edu.cn), which determines the enriched pathways by the DE genes using KEGG pathway (https://www.genome.jp/kegg/pathway.html) and REACTOME (https://reactome.org) databases as references. The cut-off criterion used was a Corrected p-Value < 0.05. The statistical test method used as hypergeometric test/Fisher’s exact test. The method used for FDR correction was Benjamini and Hochberg. The red bars represent the terms related to upregulates genes, the blue bar is related to the downregulates genes.

#### Statistical Analysis

All the statistical analyses were performed using Prism Graph Pad Software (versions 7-10). Comparison between two groups was performed by Unpaired t test. When comparing more than two groups, One-way ANOVA followed by Tukey’s multiple comparison test or Dunnett’s T3 multiple comparisons test was applied, depending on the experiment, as specified in the legend of the corresponding figure. Groups were considered statistically different from each other if the P value was less or equal to 0.05. Visually, significance was defined as: * p < 0.05, ** p < 0.01, *** p < 0.001, **** p < 0.0001. For the correlation of the SMI analysis, linear regression was applied to the SMI-generated values of two markers at the designated components, and the R^2^ value was calculated. All the *in vitro* experiments were performed at least 2 times in triplicate, unless specified in the figure legend. Mice of each background that were orthotopically injected with cancer cells were distributed into experimental groups assuring that all the groups had equal numbers of males and females. All the *in vivo* experiments had a N of at least 6 mice per group, unless specified differently in the figure legend.

## Supporting information

Key resource table

## Acknowledgments

This study is dedicated to the memories of Cyndy Francescone (our biggest fan, our eternal mom and Nana), Maria José Antero Barbosa (grandma, our “vó Maria”), Neelima Shah (a great biologist and an amazing soul), and Dr. Patricia Keely (an influential TME researcher that enormously influenced our studies), who will forever inspire our team-based work. We thank the patients that donated samples for research, as well as Jim Oesterling for his feedback and technical assistance with flow cytometry. We are grateful for the kind gifts provided by Robert Weinberg (pBABE-neo-hTERT vector; Addgene plasmid # 1774), Dr. Didier Trono (psPAX2; Addgene plasmid # 12260), Charles Gersbach (pLV hU6-sgRNA hUbC-dCas9-KRAB-T2a-Puro; Addgene plasmid #71236), Ria Baumgrass (Addgene plasmid # 74050) and Dr. Alexey Ivanov (pLV-CMV-H4-puro vector). We thank Dr. Marina Pasca di Magliano for her support with scRNAseq analysis and her advice, Dr. Stephen Sykes for his advice on the bone marrow chimera experiments, and Dr. Igor Astsaturov for his comments and suggestions.

This work was supported by funds from T32CA009035 (R. Francescone and D. Barbosa Vendramini-Costa, independently), DOD PA220131P1 (D. Barbosa Vendramini-Costa and E. Cukierman), Pancreatic Cancer Action Network (PanCAN) Career Development Award in memory of Skip Viragh 21-20-FRAN (R. Francescone), the 5th AHEPA Cancer Research Foundation, Inc. (E. Cukierman), the Pancreatic Cancer Cure Foundation (E. Cukierman), as well as NIH/NCI grants R01CA113451 (E. Cukierman and J. Franco-Barraza), R01 CA232256-S03 (D. Barbosa Vendramini-Costa and E. Cukierman), R01CA269660 (E. Cukierman), U54CA272686 (E. Cukierman), R35 GM122517 (R. Dunbrack), R00CA263154 (N. Steele), Pennsylvania DOH Health Research Formula Funds (K.S. Campbell), the equipment grant S10ODO23666, and the Core Comprehensive Cancer Center Grant CA06927 in support to FCCC’s facilities including: Bio Sample Repository, Light Microscopy, Small Animal Imaging, Biostatistics and Bioinformatics, Flow Cytometry, Laboratory Animal, Cell Culture, DNA sequencing, Genotyping and Real-time PCR, Histopathology, and Immune Monitoring Facility.

## Conflict of interest disclosure statement

The following authors disclose these IPs that have some relevance to this study: D.B.V.C, R.F., J.F.B., and E.C., with US-20220289823-A1; R.F., J.F.B., and E.C., with US-20210309729-A1; and J.F.B and E.C with US-20230152325-A1.

## Author contributions

D.B.V.C., R.F. and E.C.: conceptualization, data curation, visualization, review and editing, wrote final version of the manuscript. D.B.V.C and R.F.: *in vivo* studies, *in vitro* studies with immune cells, study design, analysis, visualization, wrote original draft of the manuscript, funding acquisition. D.B.V.C., R.F., T.L., J.C.G., S.A.A.S., C.O., A.M.A and M.G.: *in vitro* studies with fibroblasts, analysis, visualization. J.F.B., and E.C.: SHG acquisition, analysis, visualization; multiplex IF acquisition, analysis, visualization. E.M. and L.B.: SHG acquisition, analysis. Y.T.: performed scRNAseq libraries, ran scRNAseq, ran bulk RNAseq. N.S.: analysis and visualization of scRNAseq. Y.Z.: analysis of bulk RNAseq and pathway analysis of scRNAseq. D.B.V.C, H.L. and S.A.A.S.: analysis and visualization of bulk RNAseq. D.B.V.C. and E.M.: irradiation of mice, bone marrow transplants. D.I.Z.: experiments with human T cells, data analysis. K.Q.C. and A.J.K.S.: IHC staining and analysis of human TMA. K.Q.C.: IHC staining in murine tissue, analysis and quantifications, H&E scoring. M.A. and R.L.D.: peptide conceptualization and design. H.W.: patient annotation, analysis, resources. K.C.: provided KC and KPC mice, resources, advice and feedback on the project. E.C.: supervision, resources, funding acquisition.

All the authors edited the manuscript and approved the final version before submission.

---

**Sup. Figure 1:**
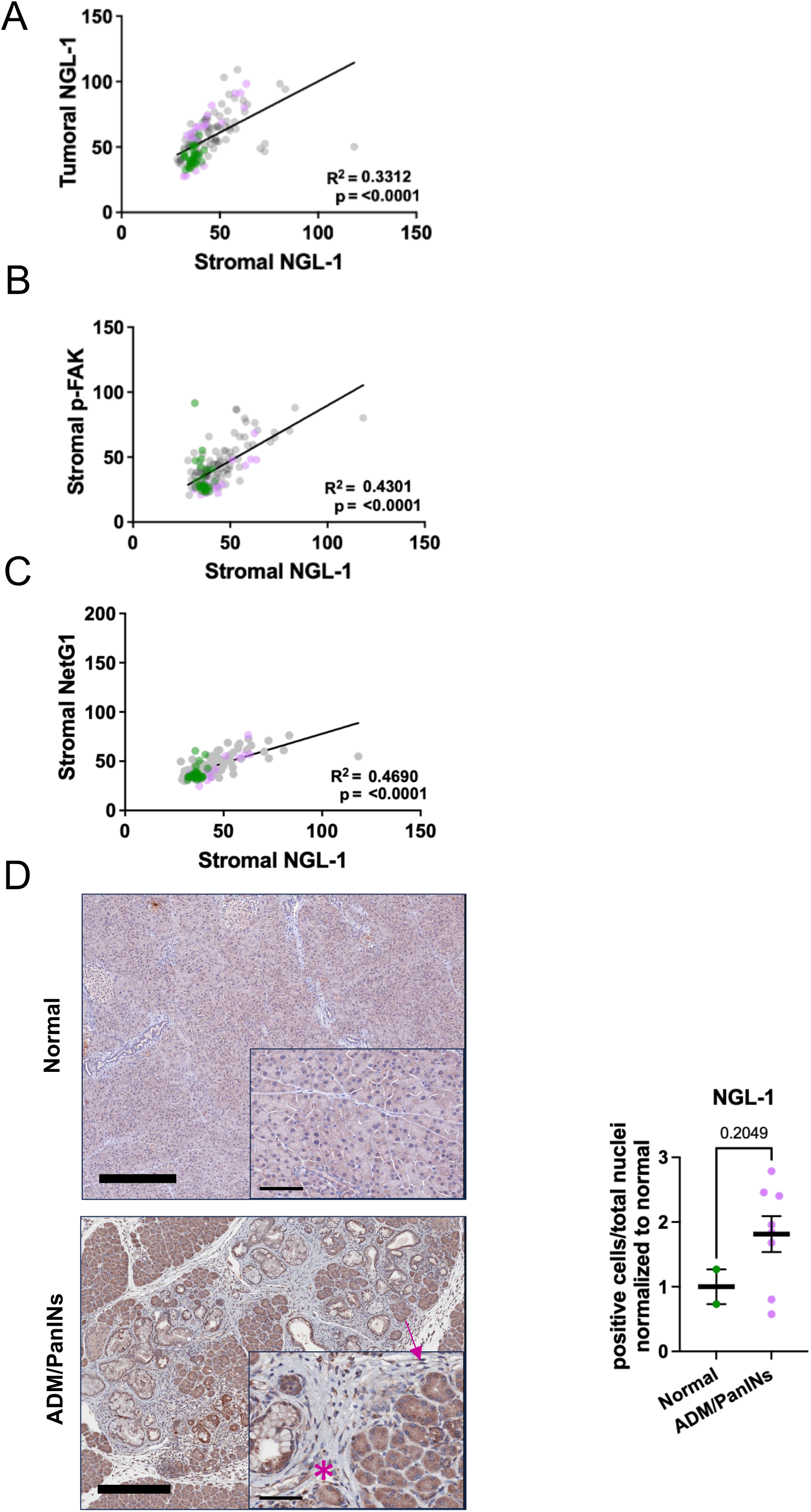
Stromal expression of NGL-1 weakly correlates with stromal expression of the pro-tumor markers NetG1 and p-FAK. **A)** Correlation between the expression of NGL-1 in tumor tissue (pan-cytokeratin positive PDAC areas) and in the stroma (pan-cytokeratin negative areas), by linear regression. Each dot represents the mean intensity of each marker in a single image (Immunofluorescence, representative images in Figure 1A. Green dots: normal pancreas. Violet dots: tumor adjacent tissue. Gray dots: PDAC tissue. **B)** Correlation between the stromal (pan-cytokeratin negative areas) expression of NGL-1 and p-FAK. **C)** Correlation between the stromal (pan-cytokeratin negative areas) expression of NGL-1 and Netrin G1 (NetG1). **D)** Representative images and quantification (mean ± SEM) of the expression of NGL-1 in normal pancreatic tissue (n = 2, ages 8-12 weeks old) and in early pancreatic lesions (n = 8, ages 6-10 months old) of KC mice. Unpaired t test, p = 0.2049. ADM: acinar to ductal metaplasia. PanINs: pancreatic intraepithelial neoplasia. Scale bar: 300 µm. Insert: 60 µm. Arrow: fibroblast. Asterisk: immune cell.

**Sup. Figure 2:**
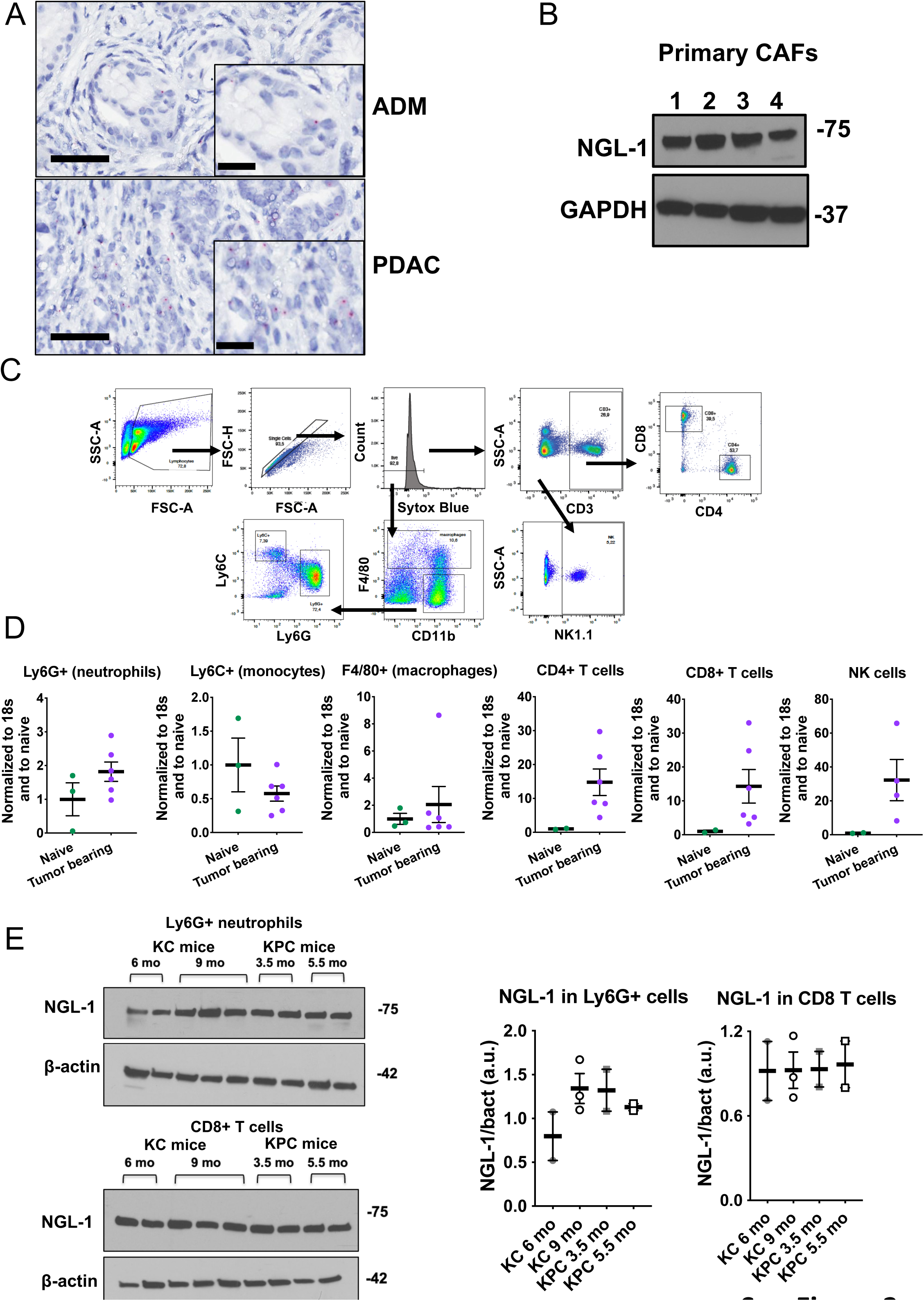
NGL-1 is expressed by many stromal cell types, correlating with disease progression. **A)** BaseScope detecting *Lrrc4c* (NGL-1, magenta puncta) in a representative ADM and PDAC lesion of KPC mice. The tissue was counterstained with hematoxylin. Scale bar: 50 µm. Inserts: 31 µm. **B)** Western blotting for NGL-1 in four primary human CAF lines. GAPDH was used as loading control. **C)** Gating strategy for sorting of different immune cell populations from the spleens of KPC tumor bearing mice, by flow cytometry. **D)** Expression of *Lrrc4c* (NGL-1) by real time PCR in different immune cells sorted from the spleens of naïve mice (green, n = 2 – 3 mice) and KPC tumor bearing mice (violet, n = 4 - 6 mice). The mRNA expression of NGL-1 was normalized with the expression of the housekeeping gene *Rn18s*, and the normalized expression of naïve was set to 1. **E)** Western blotting for NGL-1 in primary neutrophils (CD11b^+^/Ly6G^+^) and CD8^+^ T cells (CD3^+^/CD8^+^) isolated from mice representing different disease stages (KC 6 and 9 months old, n = 2-3 mice/age; KPC 3.5 and 5.5 months old, n = 2 mice/age). β-actin was used as loading control. All graphs depict mean ± SEM. ns: not significant.

**Sup. Figure 3:**
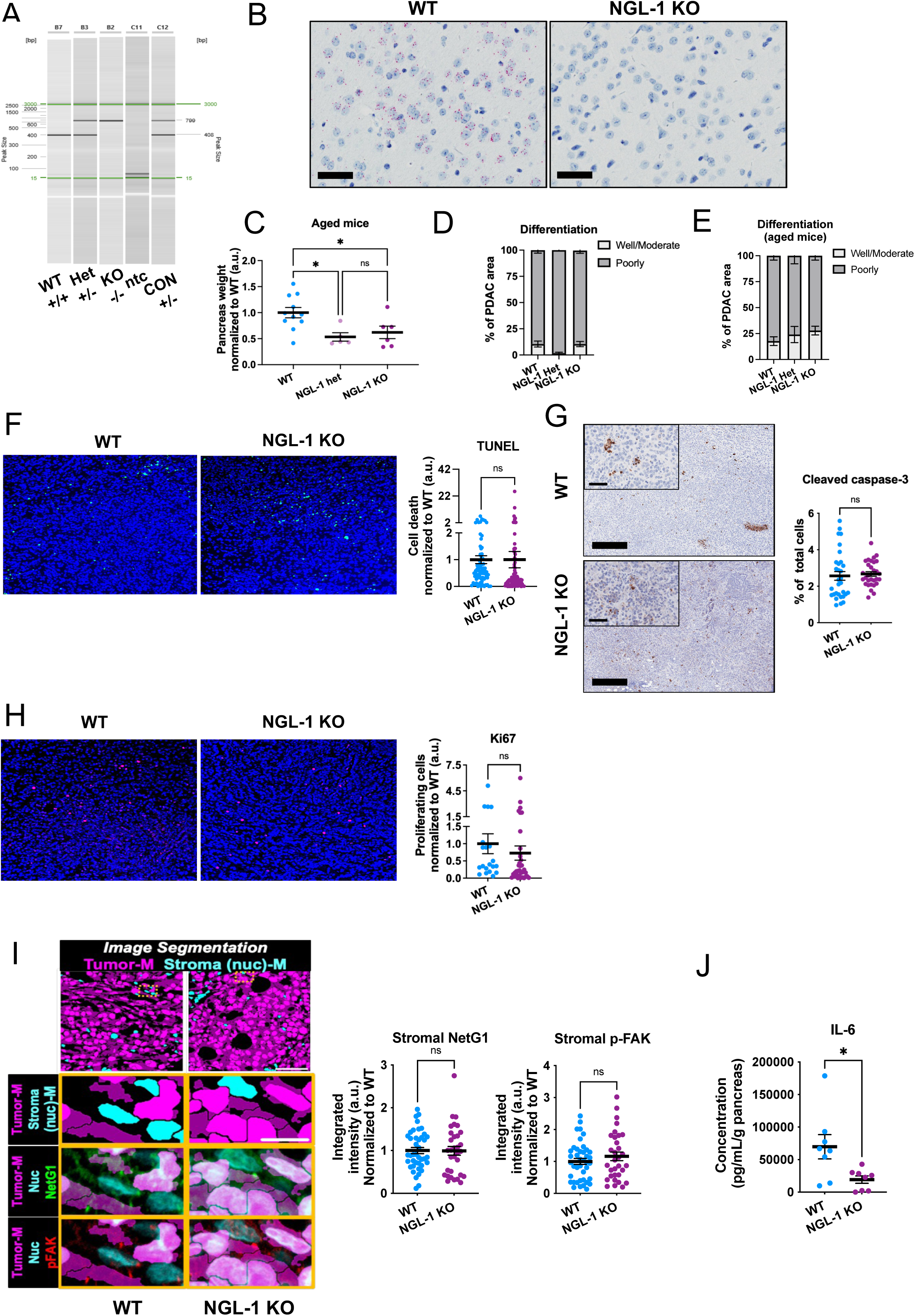
Orthotopic tumors isolated from NGL-1 deficient mice do not display differences in cell death, proliferation or stromal expression of pro-tumor markers compared to WT tumors. **A)** Representative image of the DNA genotyping targeting WT and KO NGL-1 in total DNA. Ntc: no template control. CON represents previously validated mouse heterozygous for NGL-1 (+/-) and used as a positive control (WT band ∼400 bp, KO band ∼700 bp). **B)** Representative images of the mRNA expression of *lrrc4c* in brain tissue of WT and NGL-1 KO mice, tissue expressing the highest levels of NGL-1. Note the absence of *lrrc4c* expression in the NGL-1 KO mice. mRNA is represented by the magenta puncta. Hematoxylin was used for counterstaining. Scale bar: 50 µm. **C)** Pancreas weight as a measurement of tumor burden in the different groups of aged mice (12-18 months old). WT (n = 11) in blue, NGL-1 Het (n = 5) in violet and NGL-1 KO (n = 6) in purple. The pancreas weight was normalized to the average weight of the WT group. One-Way ANOVA, Tukey’s multiple comparisons test. **D)** and **E)** Graphs depicting differentiation status (poorly differentiated, dark gray; well/moderately differentiated; light gray) of PDAC as a % of tumor area quantified from H&E images of pancreas tissue derived from **D)** younger and **E)** aged WT, NGL-1 Het or NGL-1 KO mice. **F)** Representative immunofluorescence images and quantification of the cells positive for TUNEL (green) in relation to total number of cells (nuclei stained with DAPI), as an index of cell death. The percentage of dead cells was normalized to the average percentage of death of the WT group, with no statistical difference between groups. N = 4 - 5 mice/group, each dot represents the quantification of a single image. Scale bar: 50 µm. Unpaired t test. **G)** Representative immunohistochemistry images and quantification of the cells positive for cleaved caspase 3 (brown) in relation to total number of cells, as an index of apoptosis. The percentage of apoptotic cells was normalized to the average percentage of apoptosis on the WT group, with no statistical differences between groups. WT (n = 3), NGL-1 KO (n = 3), each dot represents the quantification of a different field. Scale bar: 300 µm. Inserts: 50 µm. Unpaired t test. **H)** Representative immunofluorescence images and quantification of the cells positive for Ki67 (red) in relation to total number of cells (nuclei stained with DAPI), as an index of cell proliferation. The percentage of proliferating cells was normalized to WT group. N = 4 - 7 mice per group, each dot represents the quantification of a single image. Scale bar: 50 µm Unpaired t test. **I)** Representative digital segmentation masks (M) of SMI images of PDAC tissue from WT and NGL-1 KO mice. DRAQ5^+^ tumor nuclei in cytokeratin^+^ cells are shown in light magenta (cytoplasm in dark magenta), while DRAQ5^+^ stroma nuclei in cytokeratin^-^ cells appears in cyan (Scale bar: 50 µm). Inserts represent magnified areas marked per yellow boxes in main images, showing NetG1 (middle panel, green) or p-FAK (bottom panel, red), in cytokeratin^-^ areas (Scale bar: 10 µm). Masks are provided for reference. The quantifications of the integrated intensity of NetG1 and p-FAK normalized to the average of the WT group are presented in the graphs. N = 3 mice/group, at least 6 images per mouse. Unpaired t test. ns = no statistical significance. **J)** ELISA detecting the production of the cytokine IL-6 by fragments of pancreatic tissue from WT (n = 8) and NGL-1 KO mice (n = 8). The fragments of pancreas were weighed and cultured for 24 hours in DMEM media without serum. IL-6 concentration was calculated by pg/mL, normalized to the weight of each fragment. Unpaired t test. All graphs depict mean ± SEM. * p < 0.05. ns: not significant.

**Sup. Figure 4:**
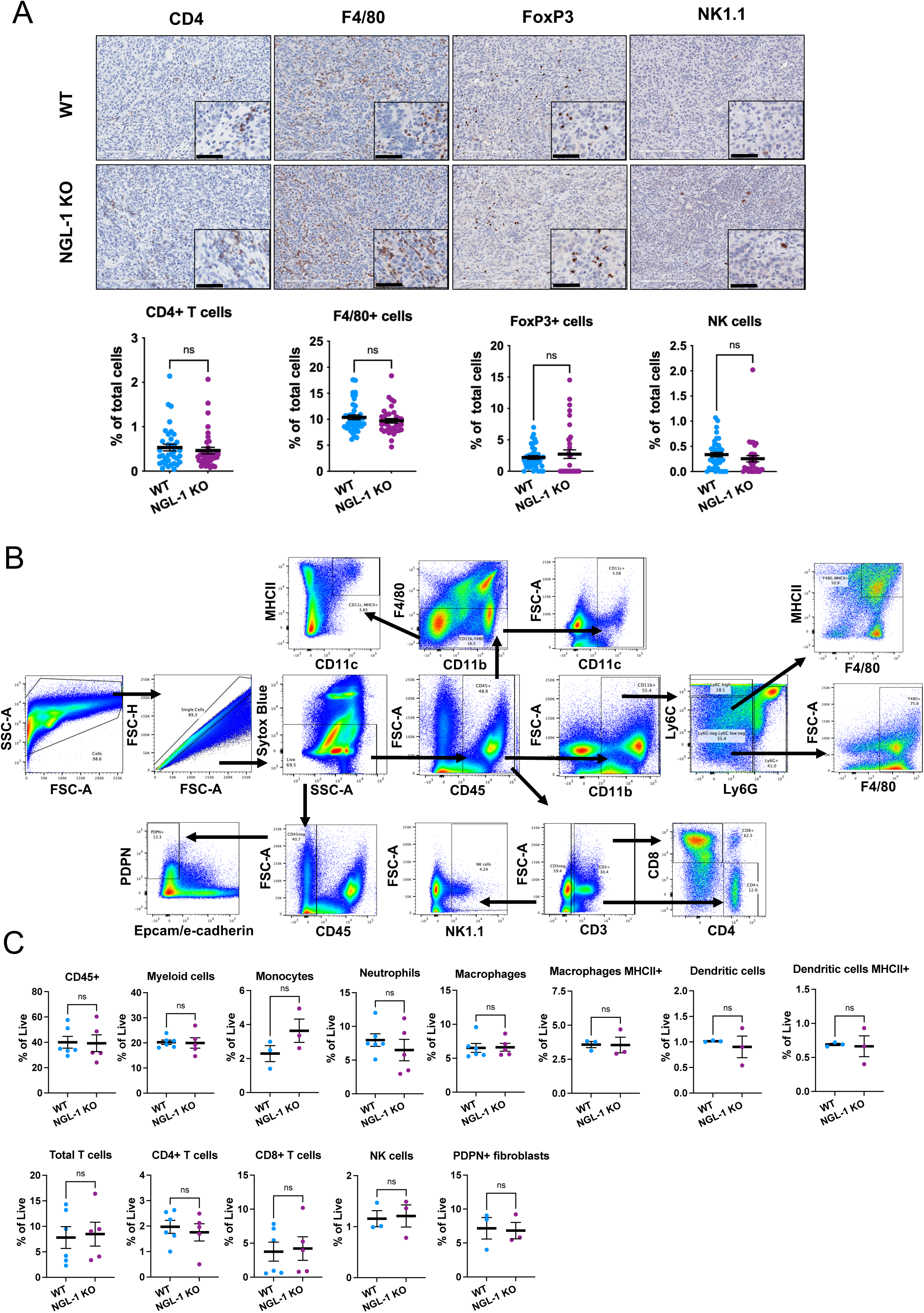
WT and NGL-1 KO tumors display similar percentages of different subsets of immune cells. **A)** Representative immunohistochemistry images of WT and NGL-1 KO tumors stained for CD4, F4/80, FoxP3 and NK1.1, to detect CD4+ T cells, macrophages, Tregs and NK cells, respectively. Markers of interest can be visualized in brown, while the tissue was counterstained with hematoxylin. Scale bars: 200 µm. Inserts: 50 µm. The graphs show the percentage of positive cells in the WT (n = 8) and NGL-1 KO (n = 7) tumors; each dot represents the quantification of a different field. Unpaired t test. **B)** Gating strategy for the evaluation of the different intratumoral immune cell subsets by flow cytometry. **C)** Percentage of the different immune cells out of total live cells in the tumors of WT and NGL-1 KO mice (n = 3 to 6 mice/group). Unpaired t test. All graphs depict mean ± SEM. ns: not significant.

**Sup. Figure 5:**
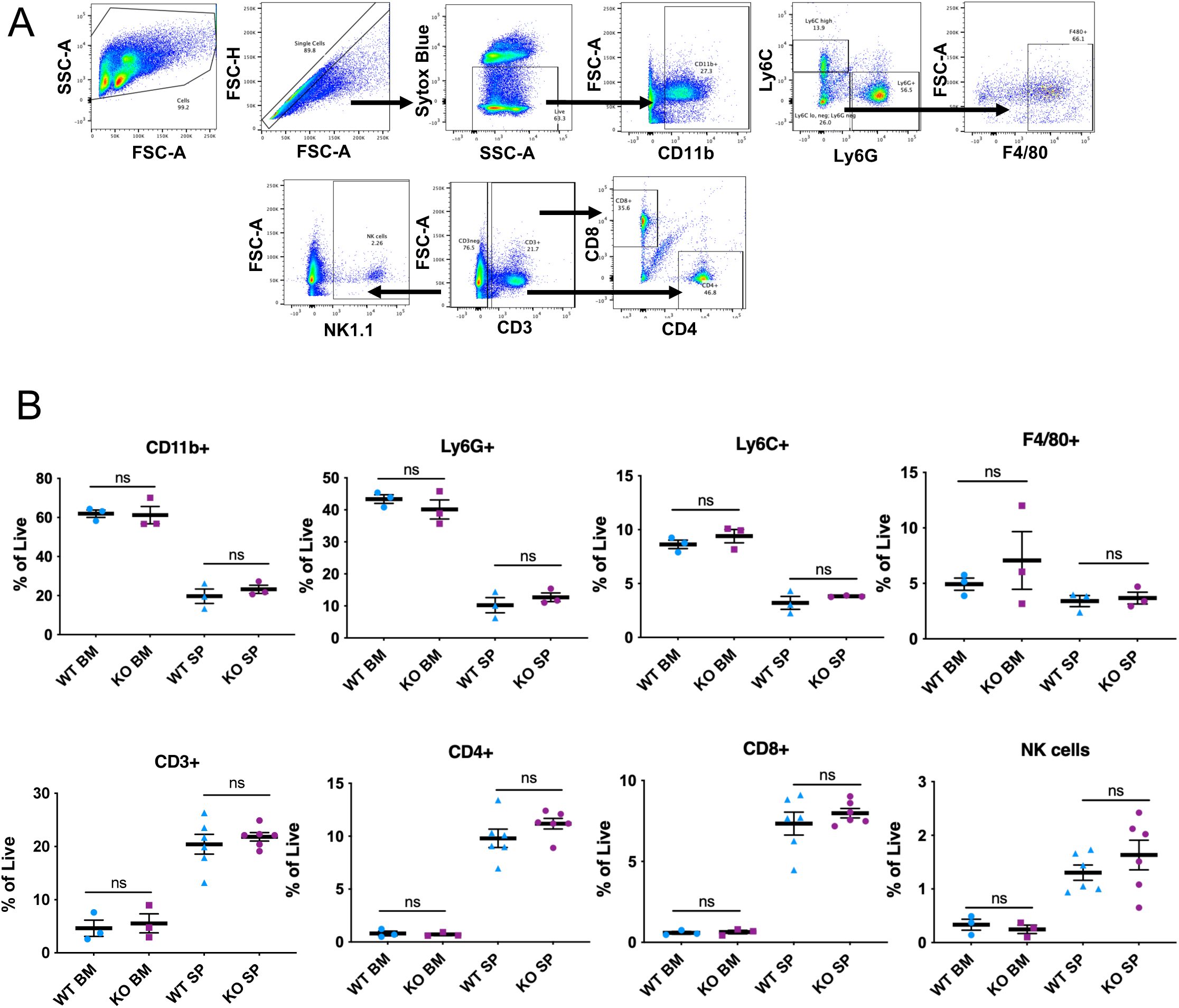
Tumor bearing NGL-1 KO mice do not present differences in peripheral immune cell composition. **A)** Gating strategy for the detection of different subsets of immune cells in the bone marrow and spleens of WT and NGL-1 KO mice. **B)** Percentages of the different immune cells out of total live cells in the tumors of WT and NGL-1 KO mice (n = 3 to 6 mice/group). All graphs depict mean ± SEM. ns: not significant.

**Sup. Figure 6:**
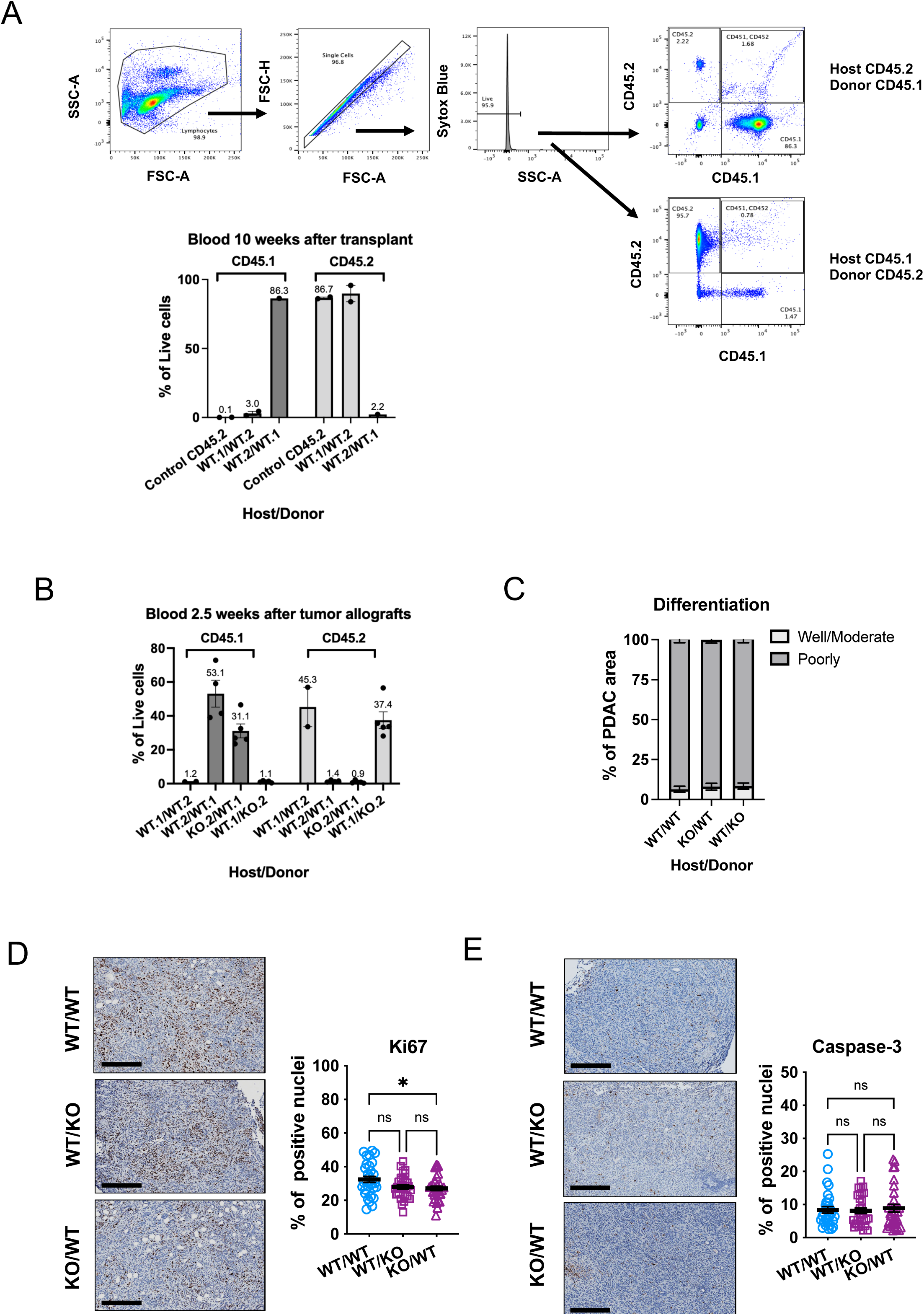
Successful Bone marrow reconstitution and chimerization for bone marrow chimera model of tumorigenesis. **A)** Gating strategy for the detection of total immune cells (CD45^+^ cells) carrying the allele *Ptprc^a^* (CD45.1) or *Ptprc^b^* (CD45.2) in the blood of chimerized mice, in order to differentiate between donor or host in the bone marrow chimera model. After 10 weeks of bone marrow transplantation, over 97% of all CD45^+^ live cells were of the donor’s background. N = 1 - 2 mice per group. **B)** Percentages of CD45.1^+^ and CD45.2^+^ T cells out of total live cells in the blood of all transplanted groups 2.5 weeks after tumor allografts (n = 2 to 5 mice/group). **C)** Graph showing that there were no differences between the different groups in terms of % of PDAC areas with well/ moderate (light gray) and poorly (dark gray) differentiated tumors, scored by a blinded pathologist. **D)** Representative immunohistochemistry images and quantification of the cells positive for Ki67 (brown) in relation to total number of cells (stained with hematoxylin), as an index of cell proliferation. The tumors of the NGL-1 KO mice transplanted with WT immune cells demonstrated a decrease in proliferation. N = 3 mice/group, each dot represents the quantification of a single field. Scale bar: 300 µm. One-Way ANOVA, Tukey’s multiple comparisons test. **E)** Representative immunohistochemistry images and quantification of cells positive for cleaved caspase 3 (brown) in relation to total number of cells (stained with hematoxylin), as an index of apoptosis, with no statistical differences between groups. N = 3 mice/group, each dot represents the quantification of a different field. Scale bar: 300 µm. One-Way ANOVA, Tukey’s multiple comparisons test. All graphs depict mean ± SEM. * p < 0.05. ns: not significant.

**Sup. Figure 7:**
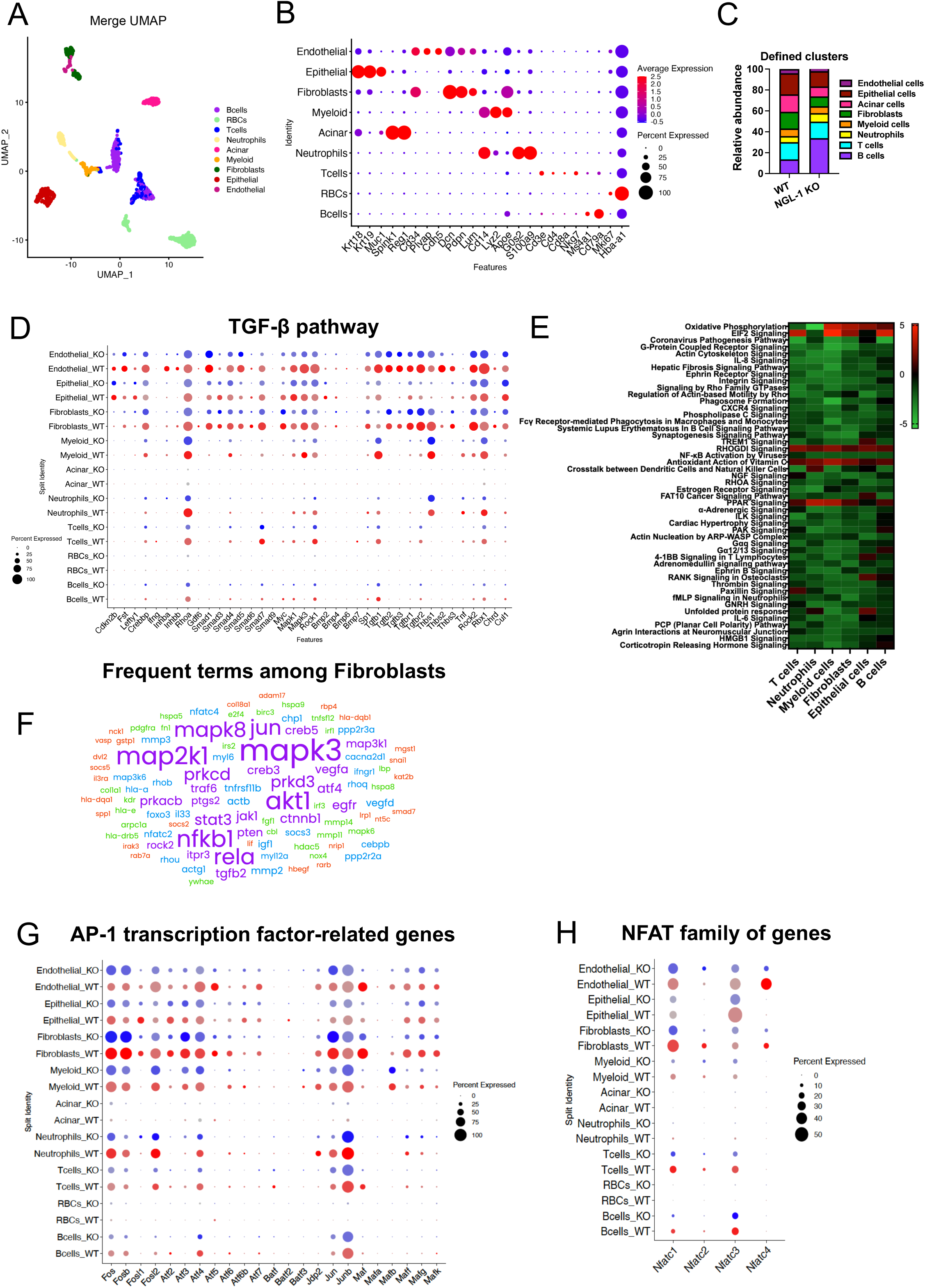
Single cell RNA sequencing analysis reveals widespread downregulation of genes in the TGF-β pathway, AP-1 transcription factors and NFAT family in NGL-1 KO tumors compared to WT tumors. **A)** UMAP dimensionality reduction plot showing the different cell clusters identified in the tumor cell suspension. Annotations show the identified cell clusters, represented by different colors. **B)** Dot plot of the key markers used to identify the different cell population. The redder the dot is, the higher is the expression of such marker, while the size of the dot represents the percentage of cells expressing that marker. **C)** Graph showing the relative abundance of the different cell populations in the tumor of WT and NGL-1 KO mice. Red Blood cells were excluded. **D)** Dot plot of the expression of genes pertaining to the TGF-β pathway expressed by the different cell populations in WT (red) and NGL-1 KO (blue) mice. The darker the dot is, the higher is the expression of such marker, while the size of the dot corresponds to the percentage of cells expressing that marker. **E)** Heatmap of the top 50 canonical pathways that were globally affected (in T cells, neutrophils, myeloid cells, fibroblasts, epithelial cells and B cells) in the tumors by the loss of NGL-1. Positive values (tones of red) indicate predicted activation, while negative values (tones of green) indicate predicted inhibition in the cells of the NGL-1 KO tumor, according to activation z-score. **F)** Representative Word cloud graph (FreeWordCloud Generator.com) showing the most common terms (genes) among all the canonical pathways enriched in the NGL-1 KO mice, here showing the analysis of fibroblasts as an example. The bigger the size of the term, the more frequent it was among all the pathways. Note the high frequency of terms related to the AP-1 transcription factors and the presence of those from NFAT family. **G) and H)** Dot plots of the expression of genes related to the **G)** AP-1 transcription factor genes and **H)** NFAT family of genes by the different cell populations in WT (red) and NGL-1 KO (blue) mice. The darker the dot is, the higher is the expression of the marker, while the size of the dot corresponds to the percentage of cells expressing that marker.

**Sup. Figure 8:**
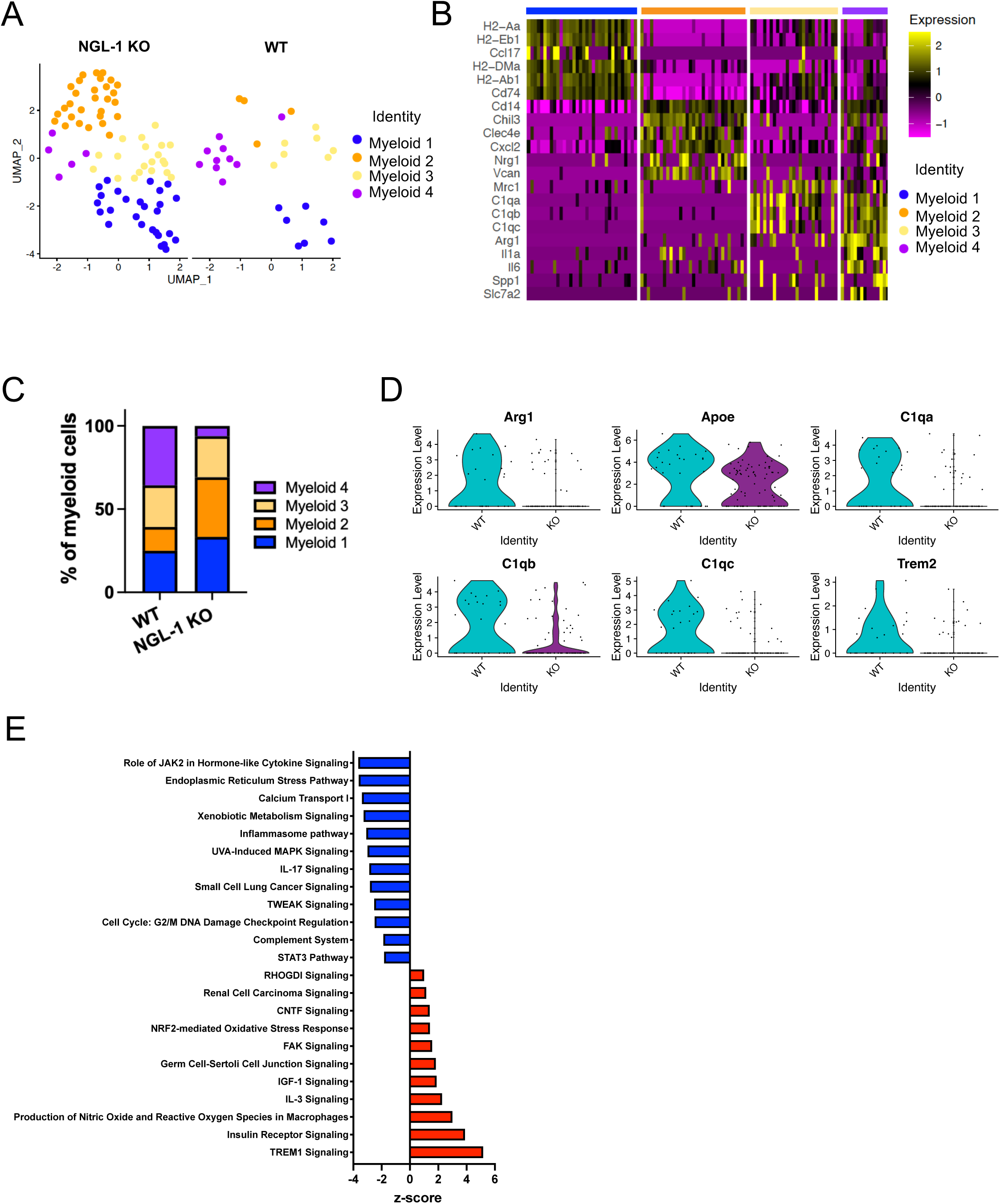
scRNAseq analysis shows that macrophages are polarized towards a less immunosuppressive phenotype in the NGL-1 KO tumor. **A)** UMAP analysis of the myeloid populations from WT and NGL-1 KO tumors. There were four different subsets of myeloid cells identified (represented by different colors). **B)** Heatmap analysis of the top genes in each identified subset. Upregulated genes are in yellow tones while downregulated genes are in magenta tones. **C)** Graph showing the relative abundance of cells in the different myeloid clusters in the tumor of WT and NGL-1 KO mice. **D)** Violin plots depicting the expression of genes related to pro-tumor phenotype in myeloid cells, at the single cell level, in WT and NGL-1 tumors. **E)** Selected canonical pathways predicted to be downregulated (in blue) or upregulated (in red), as determined by z-score, in myeloid cells from the NGL-1 KO tumor compared to WT tumor. Pathways were filtered with p < 0.01.

**Sup. Figure 9:**
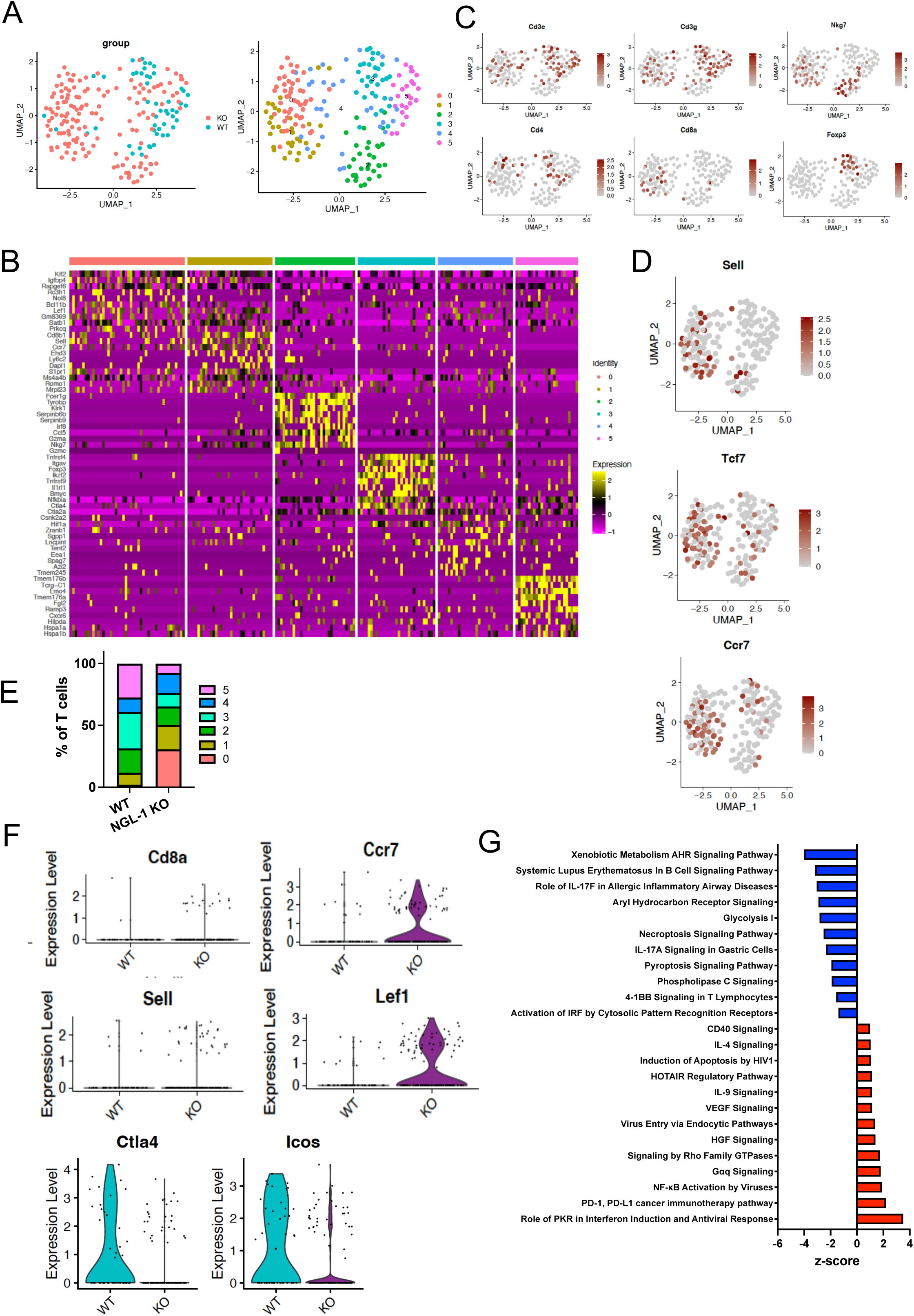
T/NK cells are polarized towards a naïve/stem like state and less exhausted phenotype in the NGL-1 KO tumor. **A)** UMAP analysis of the T/NK cell populations from WT and NGL-1 KO tumors (Left Panel). There were six different subsets of T/NK cells identified (represented by different colors; Right Panel). **B)** Heatmap analysis of the top genes in each identified subset. Upregulated genes are in yellow tones, while downregulated genes are in magenta tones. **C)** Feature plots of *Cd3e*, *Cd3g*, *Nkg7*, *Cd4*, *Cd8a*, *Foxp3* in T/NK cell population, from WT and NGL-1 KO tumors. **D)** Feature plots of *Sell, Tcf7 and Ccr7,* markers for naïve/stem like cells in T/NK cell populations, from WT and NGL-1 KO tumors. **E)** Graph showing the relative abundance of cells in the different T/NK cell clusters in the tumors of WT and NGL-1 KO mice. **F)** Violin plots depicting the expression of genes defining CD8^+^ T cells (*Cd8a*), naïve/stem like cells (*Sell and Lef1*) and exhausted cells (*Ctla4 and Icos*), at the single cell level, in WT and NGL-1 tumors. **E)** Selected canonical pathways predicted to be downregulated (in blue) or upregulated (in red), as determined by z-score, in T/NK cells from the NGL-1 KO tumor compared to the WT tumor. Pathways were filtered with p < 0.01.

**Sup. Figure 10:**
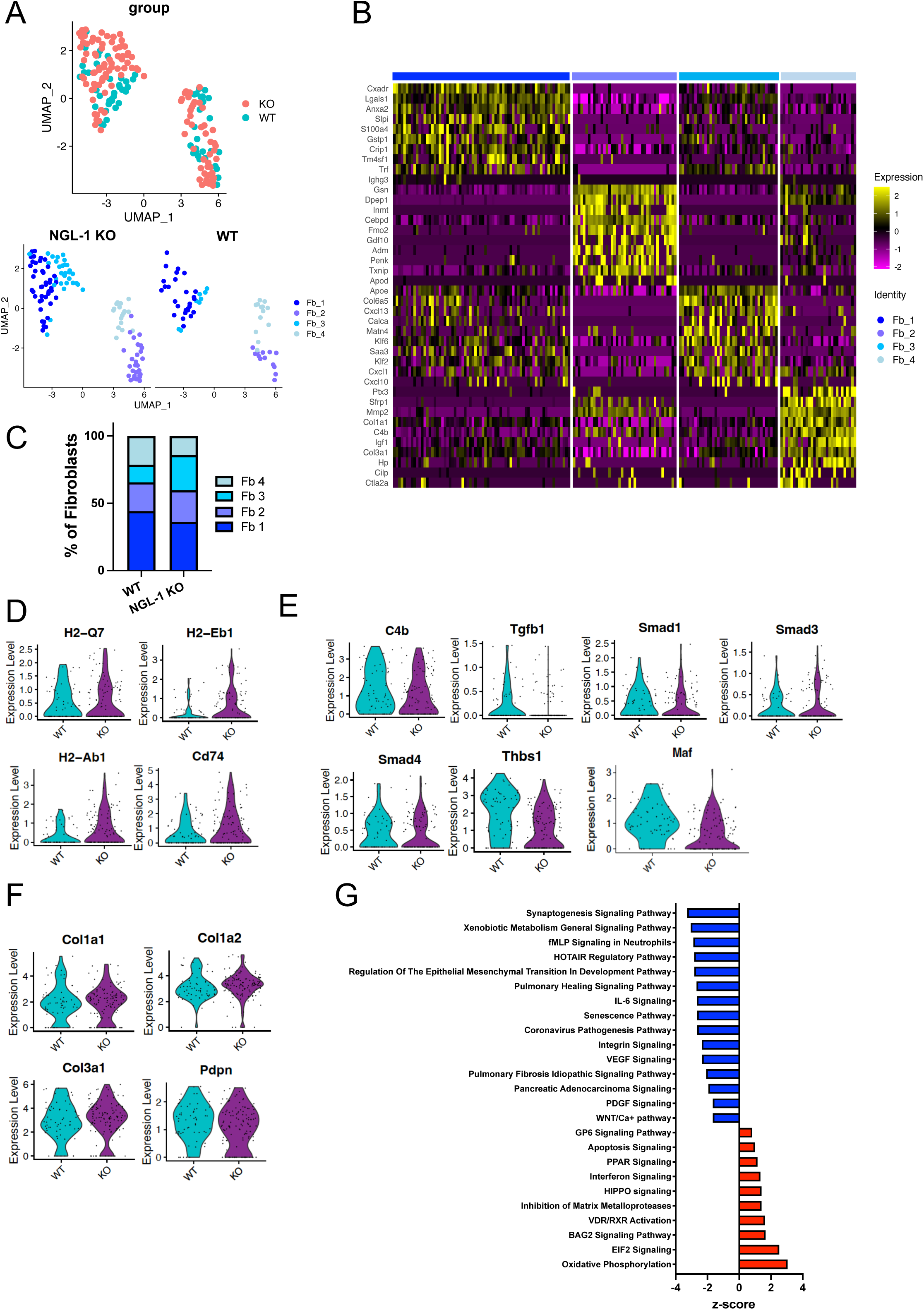
Fibroblasts from the NGL-1 KO tumors downregulate genes involved in TGF-β signaling and pro-tumoral pathways, while upregulating genes related to antigen presentation. **A)** UMAP analysis of the fibroblastic populations from WT and NGL-1 KO tumors (Top panel). There were four different subsets of fibroblastic cells identified (represented by different colors; Bottom panel). **B)** Heatmap analysis of the top genes in each identified subset. Upregulated genes are in yellow tones, while downregulated genes are in magenta tones. **C)** Graph showing the relative abundance of cells in the different fibroblastic cell clusters in the tumors of WT and NGL-1 KO mice. **D-F)** Violin plots depicting the expression of genes defining: **D)** mesothelial-like cells and immune response (antigen presentation) (*H2-Q7, H2-Eb1, H2-Ab1, Cd74)*, **E)** pro-tumoral and TGF-β signaling (*C4b, Tgfb1, Smad1, Smad3, Smad4, Thbs1, Maf*) and **F)** fibroblast/extracellular matrix components (*Col1a1, Col1a2, Col1a3, Pdpn*) at the single cell level, in WT and NGL-1 tumors. **G)** Selected canonical pathways predicted to be downregulated (in blue) or upregulated (in red), as determined by z-score, in fibroblastic cells from the NGL-1 KO tumor compared to WT tumor. Note the downregulation of important pro-tumoral pathways in PDAC. Pathways were filtered with p < 0.01.

**Sup. Figure 11:**
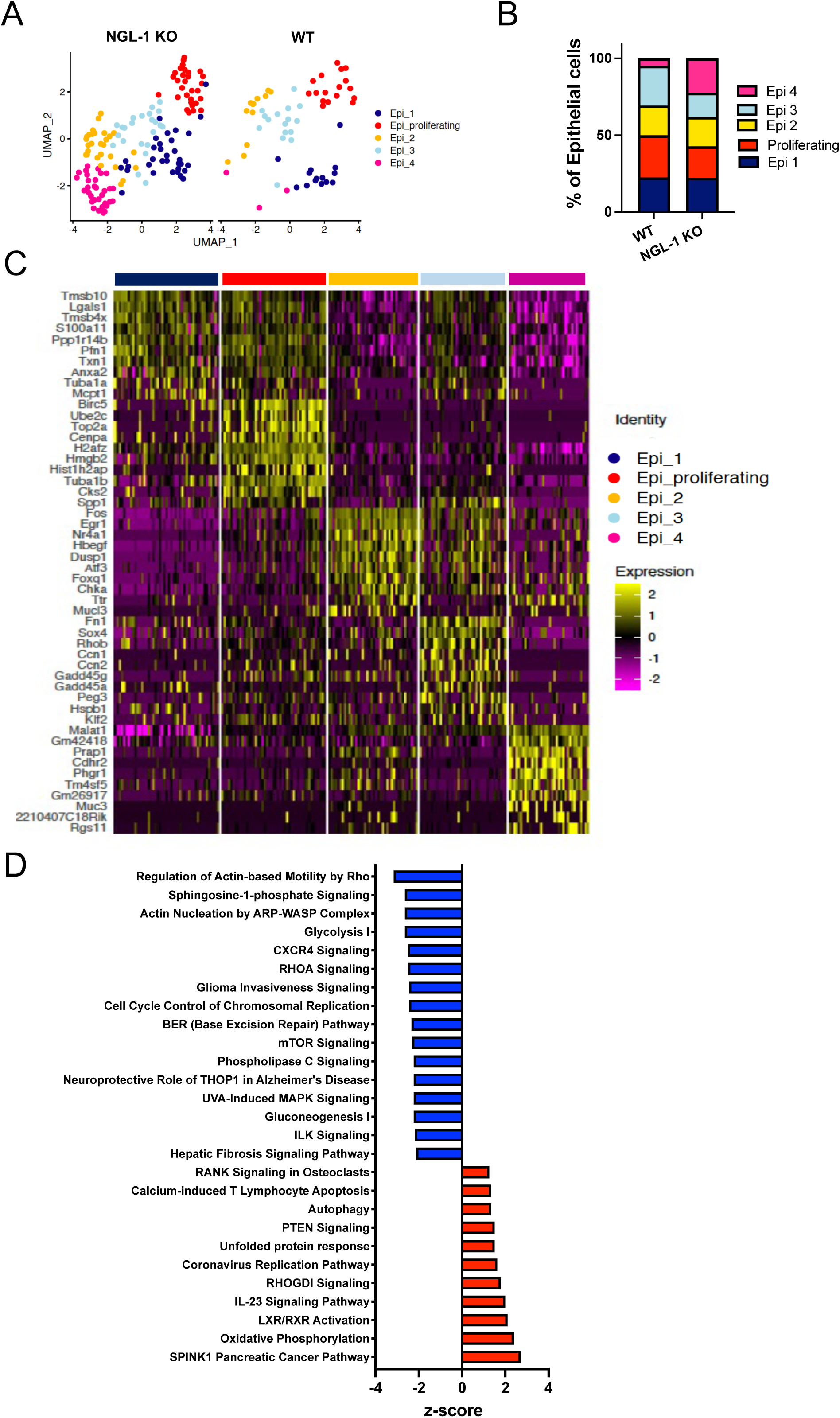
The transcriptome of epithelial cell populations differs between WT and NGL-1 KO tumors. **A)** UMAP analysis of the epithelial populations from WT and NGL-1 KO tumors. There were five different subsets of epithelial cells identified (represented by different colors). **B)** Graph showing the relative abundance of cells in the different epithelial cell clusters in the tumors of WT and NGL-1 KO mice. **C)** Heatmap analysis of the top genes in each identified subset. Upregulated genes are in yellow tones while downregulated genes are in magenta tones. **D)** Selected canonical pathways predicted to be downregulated (blue) or upregulated (red), as determined by z-score, in epithelial cells from the NGL-1 KO tumor compared with WT tumor. Pathways were filtered with p < 0.01.

**Sup. Figure 12:**
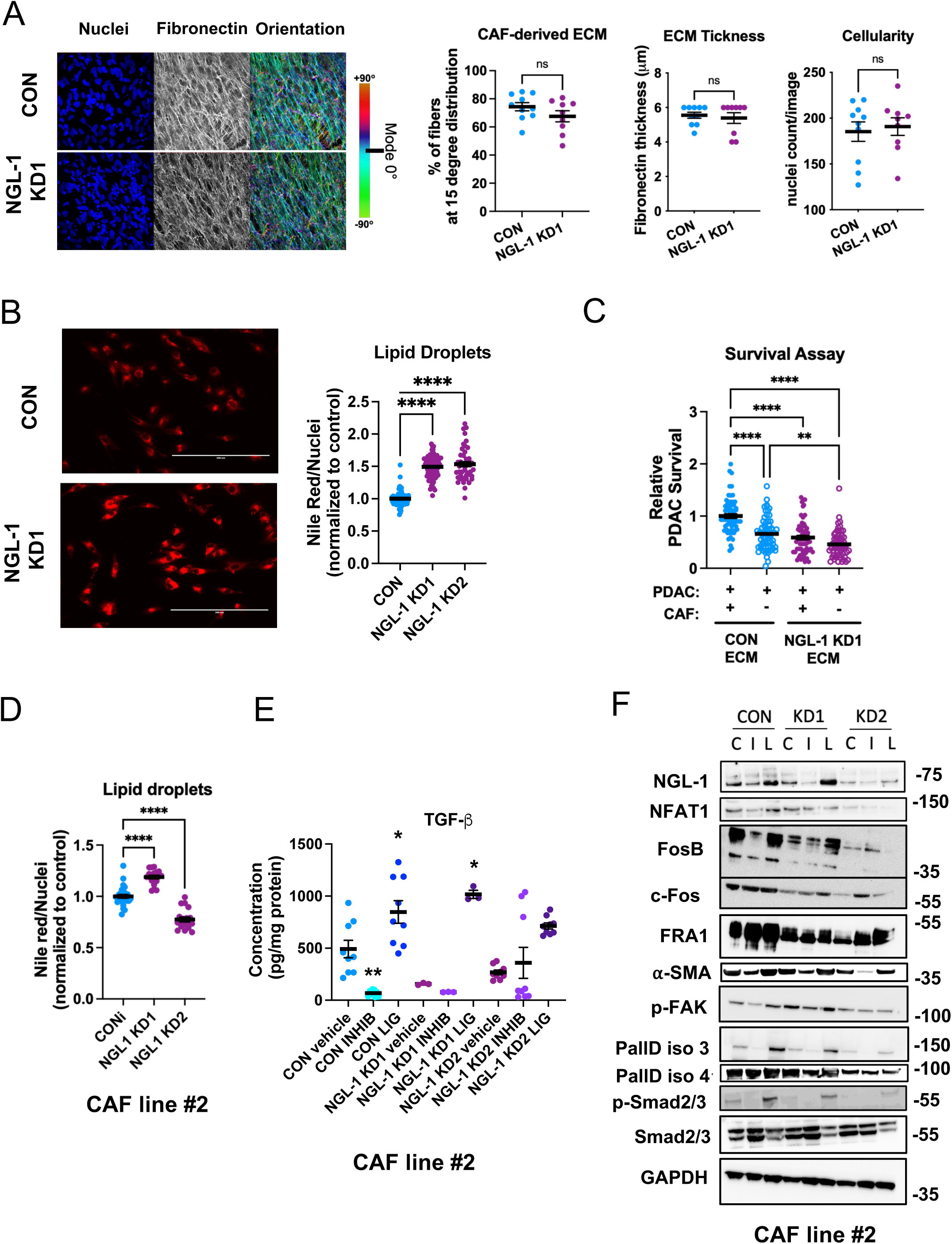
Knockdown of NGL-1 in CAFs results in decreased pro-tumor features, without disrupting ECM production. **A)** Representative images of Control (CON) and NGL-1 KD1 CAF-derived ECM produced over 5 days. Representative immunofluorescence images display cellularity (nuclei, DAPI, blue), fibronectin fibers (white) and pseudo-colored images of the fibronectin fibers depicting their orientation angles. The bar represents the distribution of angles, with the mode angle set as 0 **°** (cyan). The graph shows the level of fiber alignment, set as the percentage of fibers presenting angles within 15 degrees from the mode angle, fibronectin thickness and cellularity (nuclei counts/image). Only experiments with substantial ECM production were considered (ECM thickness above 4 µm). Unpaired t test. **B)** Representative images of Control and NGL-1 KD1 and KD2 CAFs stained with Nile Red (red) for the detection of lipid droplets retained intracellularly, as an index of CAF normalization. Cells were identified by nuclear staining (DAPI, not shown). The graph shows the quantification of Nile Red intensity normalized to DAPI^+^ nuclei. All values were normalized to the levels of the control group (CON), set as 1. Each experiment was repeated three times, at least 6 wells/cell line/experiment. One-Way ANOVA, Dunnett’s multiple comparisons test. Scale bar: 200 µm. **C)** Survival assay of RFP^+^ PDAC cells (2×10^4^ cells) co-cultured (filled circles) or not (empty circles) with GFP^+^ CON or NGL-1 KD1 CAFs (2×10^4^ cells) in the ECM produced by CON or NGL-1 KD1 CAFs, in the absence of serum and glutamine for 4 days. The graph shows the relative survival of PDAC cells normalized to the control condition (co-cultured with CON CAFs, within CON ECM). Any condition lacking NGL-1 is sufficient to impair PDAC survival. The experiment was conducted twice, with six replicates per condition. At least 4 images were acquired per replicate. One-Way ANOVA, Tukey’s multiple comparisons test. **D)** Quantification of Nile Red intensity normalized to DAPI^+^ nuclei in the CAF line #2, CON and NGL-1 KD1 and 2. All values were normalized to the levels of CON, set as 1. Each experiment was repeated two times, at least 9 wells/cell line/experiment. One-Way ANOVA, Dunnett’s multiple comparisons test. **E)** Graph depicting quantification of TGF-β ELISA detected in the conditioned media of CAF line #2, Control (CON), NGL-1 KD1 and NGL-1 KD2 CAFs growing in 3D, in the presence of vehicle (DMSO + citric acid), recombinant TGF-β or TGF-β receptor inhibitor. N = 3 independent experiments, with 3 replicates of each sample/experiment. One-Way ANOVA, Dunnett’s multiple comparisons test, compared to Control treated with vehicle. **F)** Representative western blotting for myofibroblastic markers (PallD iso 3, PallD iso 4, p-Smad2/3 and Smad2/3), NGL-1, NFAT1, AP1 transcription factor members c-Fos and FRA-1 in a second CAF line (CAF line #2). Control, NGL-1 KD1 and NGL-1 KD2 CAFs treated with recombinant TGF-β (ligand, L) and TGF-β receptor inhibitor (inhibitor, I), and their respective vehicles (control, C, Citric acid + DMSO). GAPDH was used as loading control, N = 3. All graphs depict mean ± SEM. * p < 0.05, ** p < 0.01, **** p < 0.0001. ns: not significant.

**Sup. Figure 13:**
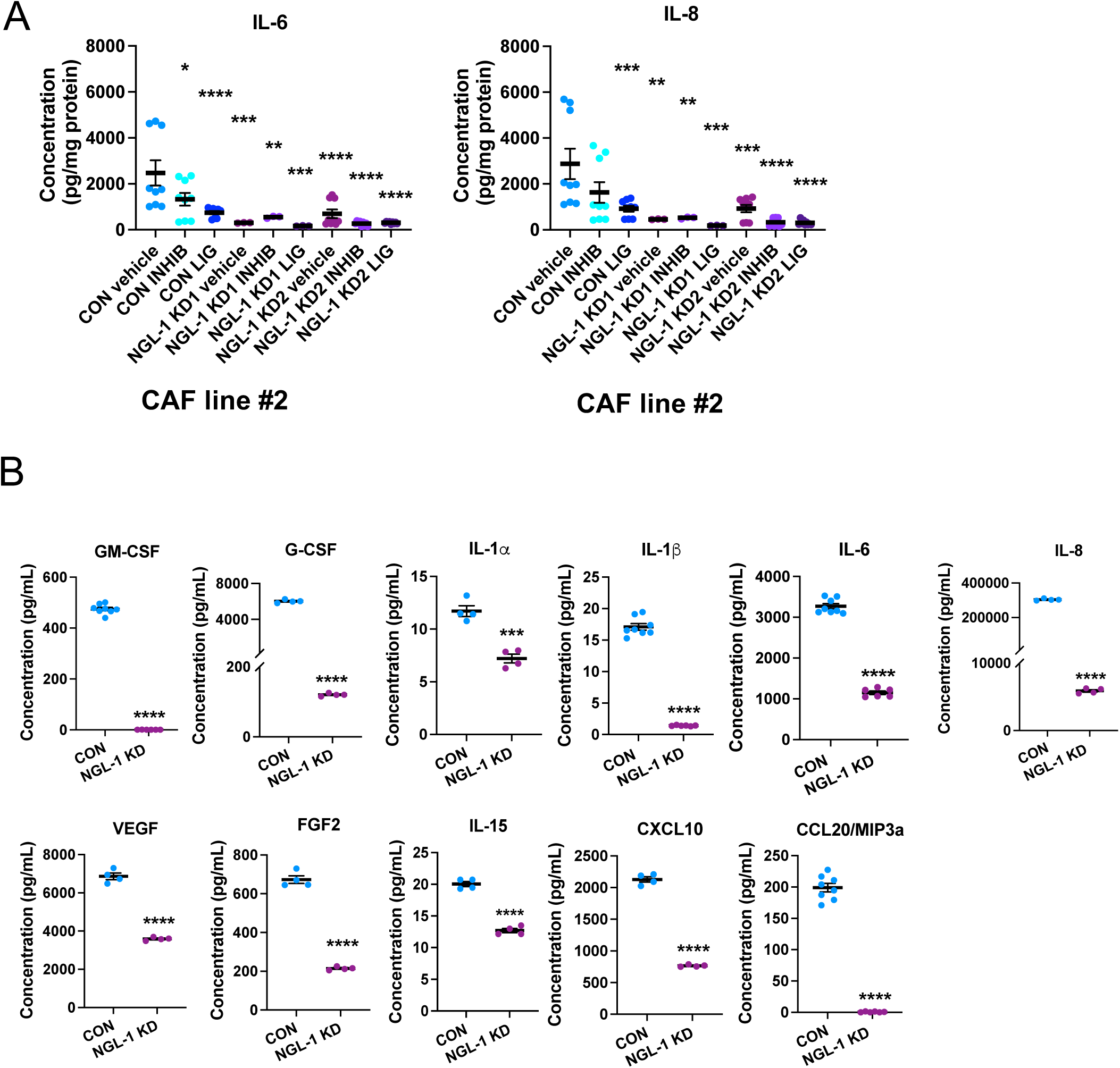
CAFs lacking NGL-1 produce less pro-tumor and immunosuppression factors. **A)** ELISA showing the decreased production of the immunosuppressive cytokines IL-6 and IL-8 by the NGL-1 KD CAFs compared to Control CAFs. The cytokines were detected in the conditioned media of Control (CON), NGL-1 KD1 and NGL-1 KD2 CAFs growing in 3D, in the presence of vehicle (DMSO + citric acid), recombinant TGF-β or TGF-β receptor inhibitor. N = 3 independent experiments, with 3 replicates of each sample/experiment. One-Way ANOVA, Dunnett’s multiple comparisons test, compared to Control treated with vehicle. **B)** Quantification of Milliplex (multiplex ELISA) for GM-CSF, G-CSF, IL1α, IL1β, IL6, IL8, VEGF, FGF2, IL-15, CXCL10 and CCL2/MIP3a) produced in the conditioned media of CON and NGL-1 KD1 CAFs, growing in 3D. N = 4 - 8 biological replicates. Unpaired t tests. All graphs depict mean ± SEM. * p < 0.05, ** p < 0.01, *** p < 0.001, **** p < 0.0001. ns: not significant.

**Sup. Figure 14:**
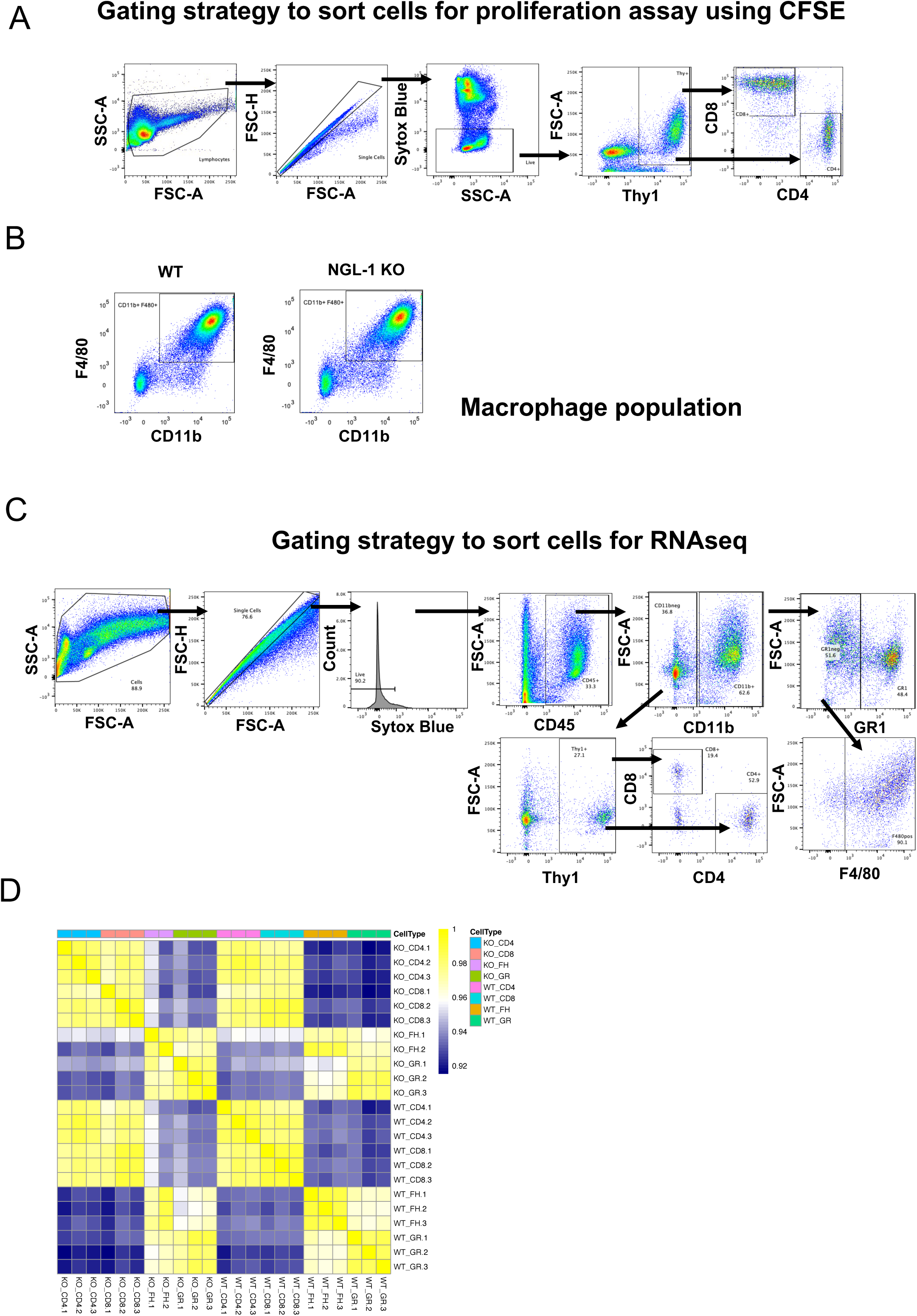
Gating strategy to obtain murine immune cell populations for functional experiments and bulk RNAseq. **A)** Gating strategy used to identify CD4^+^ and CD8^+^ T cells from total murine splenocytes by flow cytometry. **B)** Gating strategy to identify the enrichment in macrophage cells (CD11b^+^, F4/80^+^ cells) derived from the bone marrow of WT and NGL-1 KO mice. **C)** Gating strategy used to sort GR1^+^ cells (granulocytes), CD4^+^ and CD8^+^ T cells and F4/80^+^ cells (macrophages) from the PDAC tumors of WT and NGL-1 KO mice. The selected populations were sorted directly in Trizol for subsequent RNA extraction and RNA sequencing analysis. **D)** Heatmap of Pearson’s correlation to visualize the correlation between samples from the same group and from different groups, as a measurement of potential variability between samples. The bar shows the range of correlation coefficients, with the lowest correlation being 0.92 (dark blue) and the highest correlation being 1 (dark yellow). This matrix shows that there all the replicates in a group correlate, without outliers.

**Sup. Figure 15:**
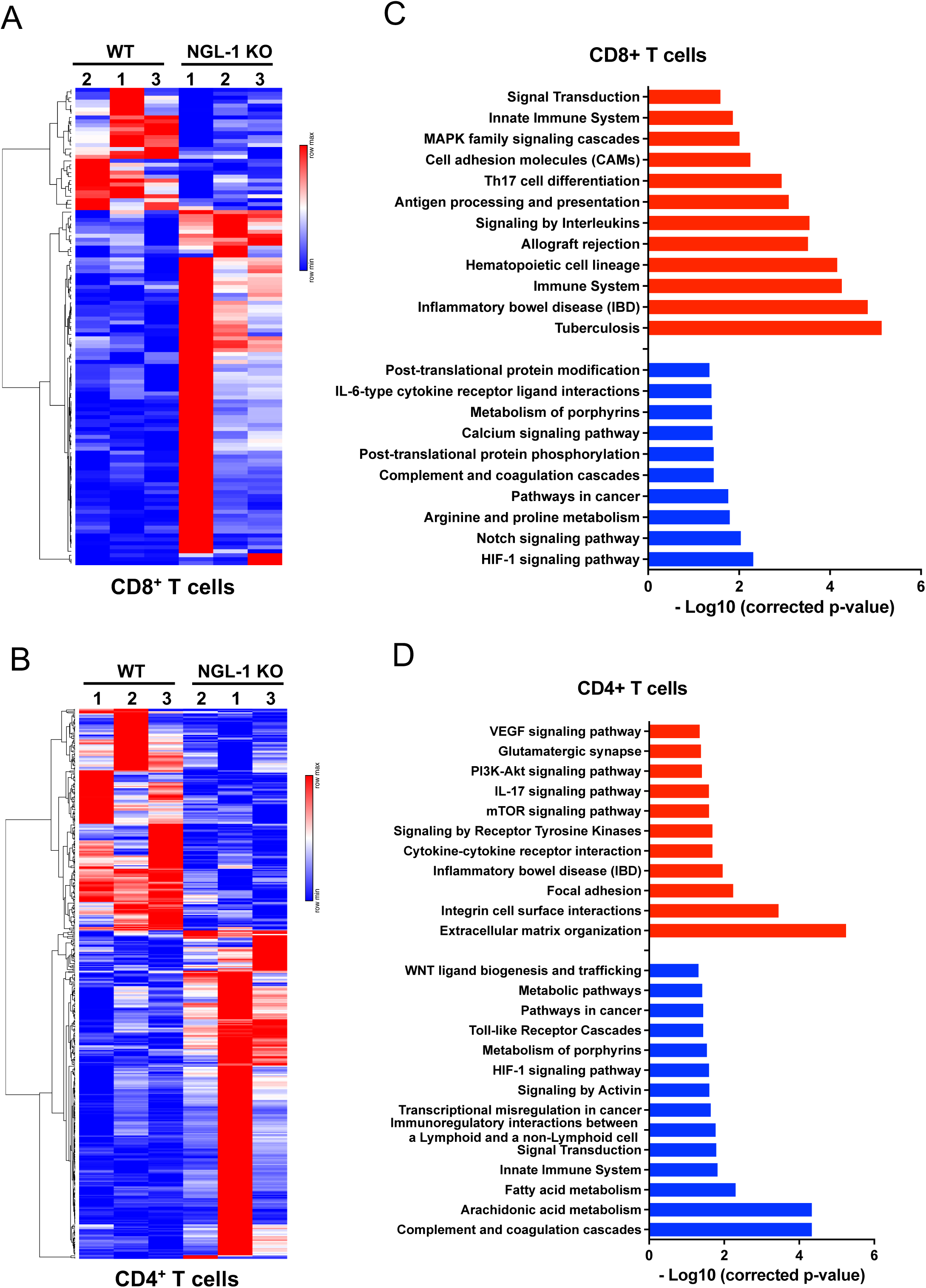
Bulk RNAseq analysis in sorted tumoral CD8^+^ and CD4^+^ T cells display an upregulation of immunostimulatory pathways in murine immune cells deficient in NGL-1. **A)** and **B)** Heatmap of the differentially expressed genes between WT and NGL-1 KO in **A)** CD8^+^ T and **B)** CD4^+^ T cells isolated from PDAC tumors, showing upregulated genes in red and downregulated genes in blue. All genes represented in the heatmap are statistically significantly (adjusted p-values < 0.05) up or downregulated in the NGL-1 KO samples compared to WT samples. **C)** and **D)** Bar graphs showing selected pathways from gene set enrichment analysis performed using the KEGG Orthology Based Annotation System (KOBAS) comparing WT and NGL-1 KO **C)** CD8^+^ T and **D)** CD4^+^ T cells isolated from PDAC tumors. Pathways predicted to be upregulated in the NGL-1 KO cells are shown in red, and those predicted to be downregulated are shown in blue. The data is represented in -Log10 scale, and all pathways were filtered with Padj < 0.05. N = 3 per group, where each sample is composed by sorted cells from 2 mice pooled together.

**Sup. Figure 16:**
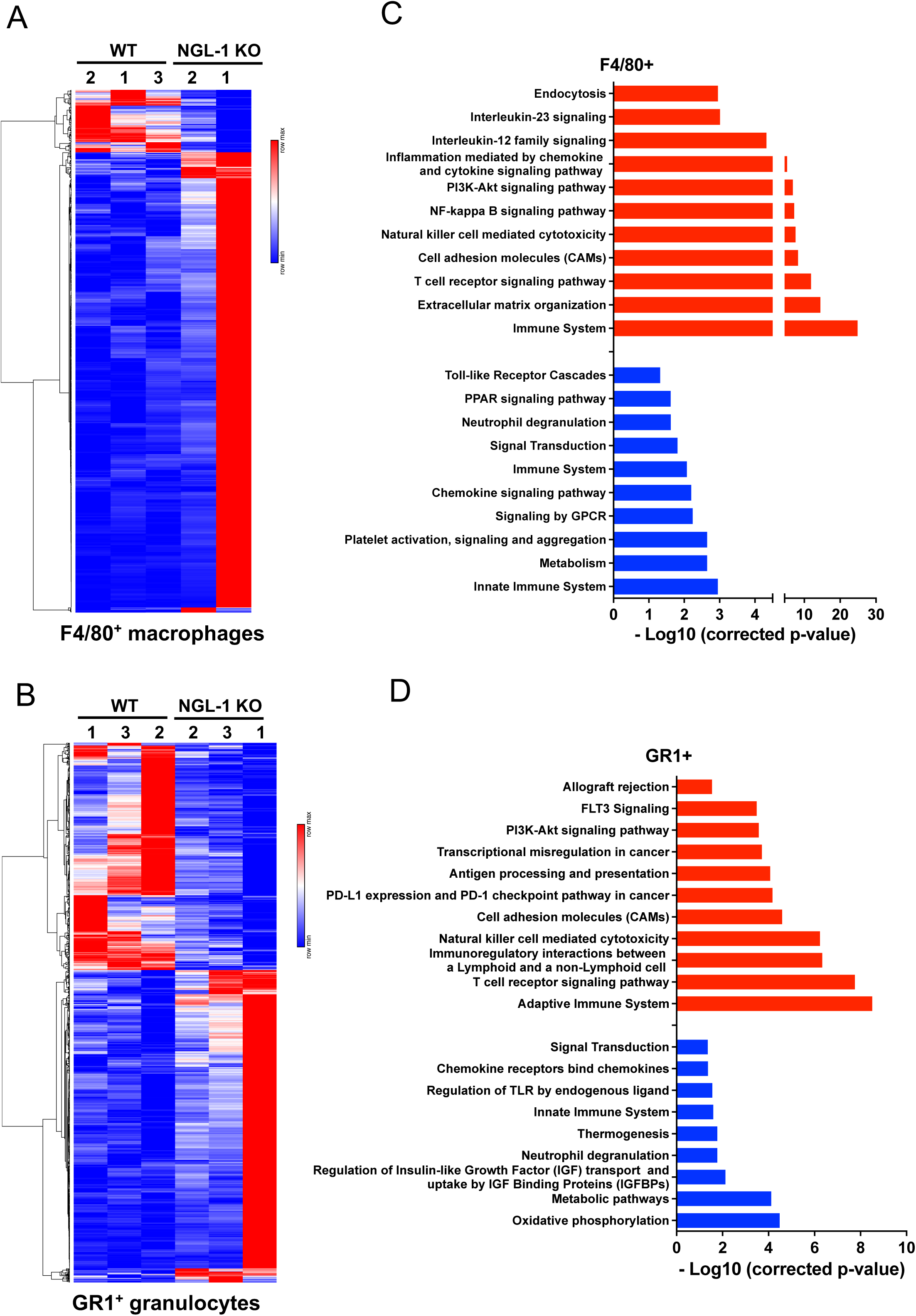
Bulk RNAseq analysis in tumoral myeloid cells (macrophages and granulocytes) shows an upregulation of immunostimulatory pathways in murine immune cells deficient in NGL-1. **A)** and **B)** Heatmap of the differentially expressed genes between WT and NGL-1 KO **A)** F4/80^+^ macrophages and **B)** GR1^+^ granulocytes isolated from PDAC tumors, showing upregulated genes in red and downregulated genes in blue. All genes represented in the heatmap are statistically significantly (adjusted p-values < 0.05) up or downregulated in the NGL-1 KO samples compared to WT samples. **C)** and **D)** Bar graphs showing selected pathways from gene set enrichment analysis performed using the KEGG Ontology Based Annotation System (KOBAS) comparing WT and NGL-1 KO **B)** F4/80^+^ macrophages and **D)** GR1^+^ granulocytes isolated from PDAC tumors. Pathways predicted to be upregulated in the NGL-1 KO cells are shown in red, and those predicted to be downregulated are shown in blue. The data is represented in -Log10 scale, and all pathways were filtered with Padj < 0.05. N = 2 - 3 per group, where each sample is composed of sorted cells from 2 mice pooled together.

**Sup. Figure 17:**
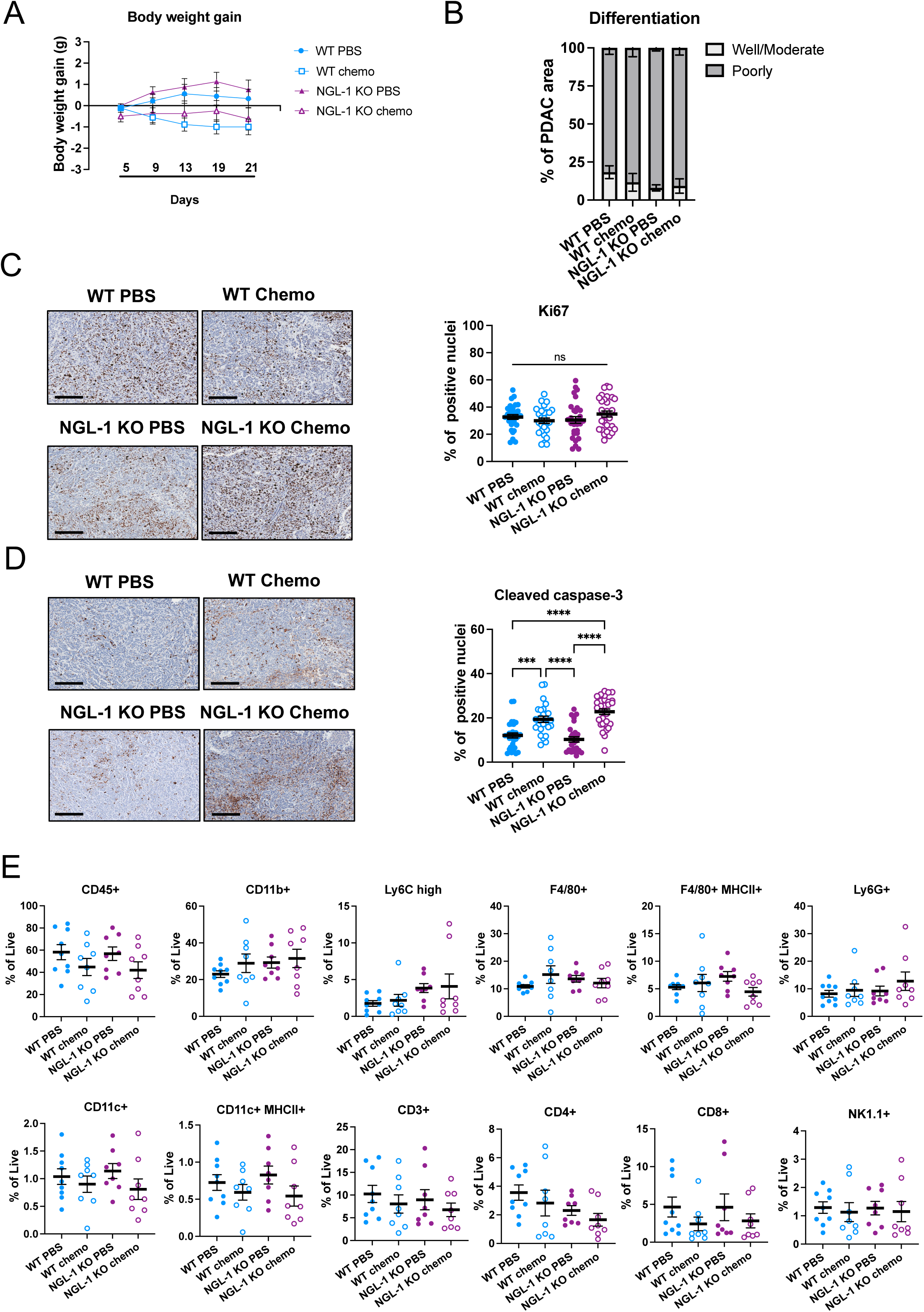
Treatment with chemotherapy leads to increased cell death and decreased infiltration of T cells in the tumors, independently of NGL-1 expression. **A)** Body weight gain in grams of the WT PBS, WT chemo, NGL-1 KO PBS and NGL-1 KO chemo groups throughout the experiment. There were no differences in body weight gain between groups in the selected dose and regimen for this experiment. **B)** Graph depicting the differentiation status of tumor areas from tumor tissue isolated from treated mice. No differences were identified between groups, as tumor tissue contained similar % areas with well/ moderate (light gray) or poorly (dark gray) differentiated tumor cells, scored by a blinded pathologist. **C)** Representative immunohistochemistry images and quantification of the cells positive for Ki67 (brown) in relation to total number of cells (stained with hematoxylin), as an index of cell proliferation. There were no differences in proliferation between groups. N = 3 mice/group, each dot represents the quantification of a single field. Scale bar: 300 µm. **D)** Representative immunohistochemistry images and quantification of the cells positive for cleaved caspase 3 (brown) in relation to total number of cells (stained with hematoxylin), as an index of cell death. There were more dead cells in the groups treated with FOLFIRINOX, independently of NGL-1 expression. N = 3 mice/group, each dot represents the quantification of a different field. Scale bar: 300 µm. One-Way ANOVA, Tukey’s multiple comparisons test. **E)** Immunophenotyping of the tumors from the different groups. The tumors were digested and the single cell suspensions were analyzed by flow cytometry. All cell populations were gated in singlets and then in live cells. Percentages of each cell subset is in relation to the % of live cells. There were tendencies for decreased percentages of T cells in the groups treated with chemotherapy, independently of NGL-1 expression, compared to the groups treated with PBS. One-Way ANOVA, Dunnett’s multiple comparisons test. All graphs depict mean ± SEM. *** p < 0.001, **** p < 0.0001. ns: not significant.

**Sup. Figure 18:**
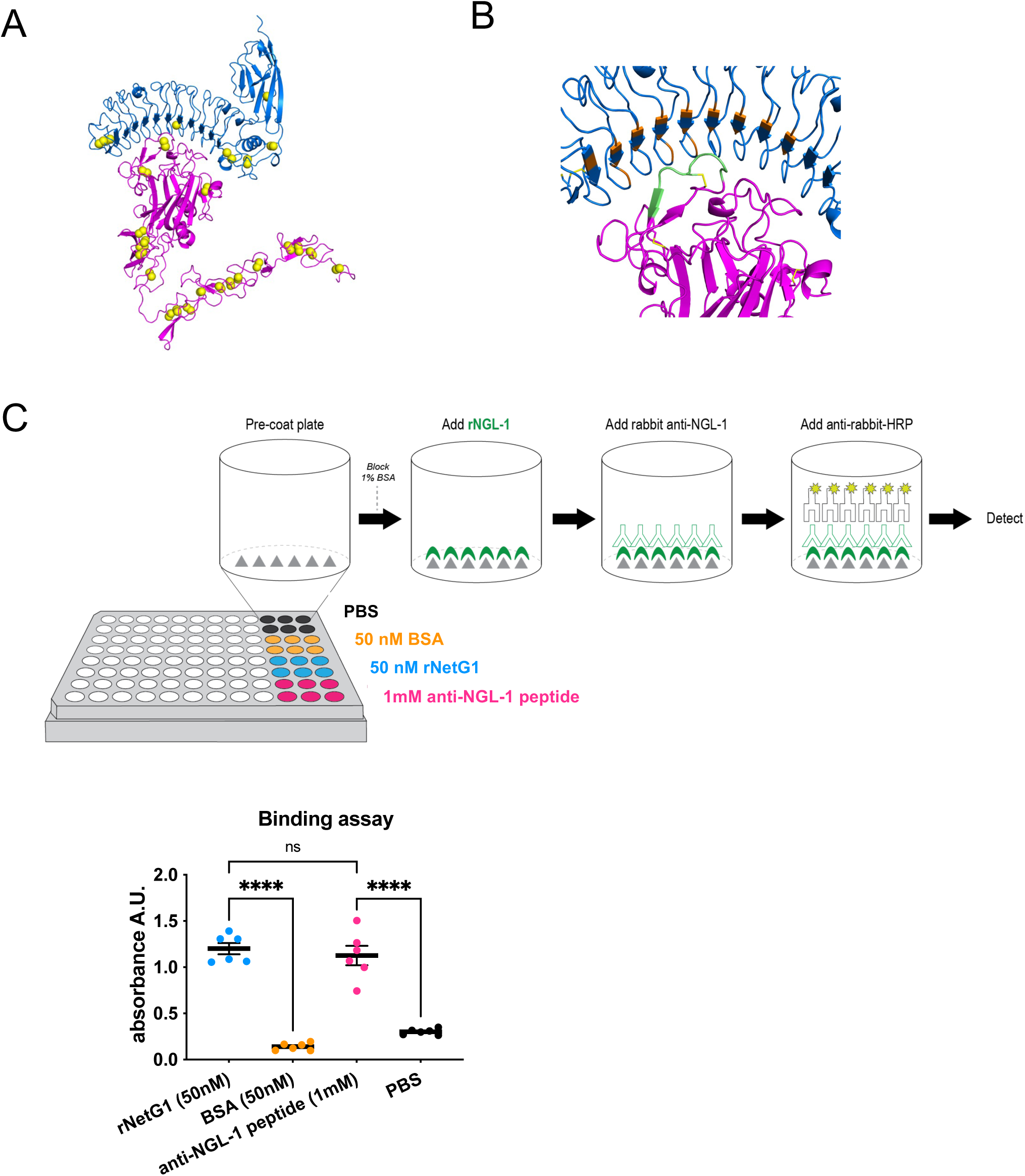
Design and validation of a small peptide targeting NGL-1. **A)** AlphaFold-Multimer model of the complex of the extracellular portions of NetG1 (magenta) and NGL-1 (blue), excluding the signal sequences and C-terminal disordered region of NGL-1 (residues 29-510 of NetG1, UniProt entry Q9Y2I2 and residues 45-448 of NGL-1, UniProt entry Q9HCJ2). The sulfur atoms of all cysteine residues appear in yellow spheres. All of them are in disulfide bonds, except C207 of NGL-1 which does not have a cysteine partner residue anywhere nearby. **B)** Closeup of NetG1/NGL-1 interface. The peptide chosen as an inhibitory molecule (residues 84-91 of NetG1, sequence NPYMCNNE with C->S mutation, NPYMSNNE) is shown in green. All 19 residues within 5 Å of any residue in this segment in NGL-1 are shown in orange (K59, L80, N82, H84, T104, Q106, T128, E130, F132, E152, W154, R156, R176, D178, Y201, N203, E223, K247, W248). **C)** Scheme of the Binding assay by ELISA, where wells in a 96 well plate were coated with either anti-NGL-1 Peptide (1 mM), recombinant NetG1 (50 nM), PBS or BSA as control (50 nM). The wells were then treated with recombinant NGL-1, and binding was allowed to occur. Next, after washing, wells were treated with an antibody against NGL-1 for 2h, and then an antibody against rabbit antibodies, linked to HRP, was added. Finally, TMB substrate was added and the absorbance signified the amount of NGL-1 bound to the well. Graph depicting the absorbance (a.u.), where only NGL-1 that bound to either peptide or to recombinant NGL-1 was detected by absorbance (450 nm). One-Way ANOVA, Tukey’s multiple comparisons test. All graphs depict mean ± SEM. **** p < 0.0001.

**Sup. Figure 19.**
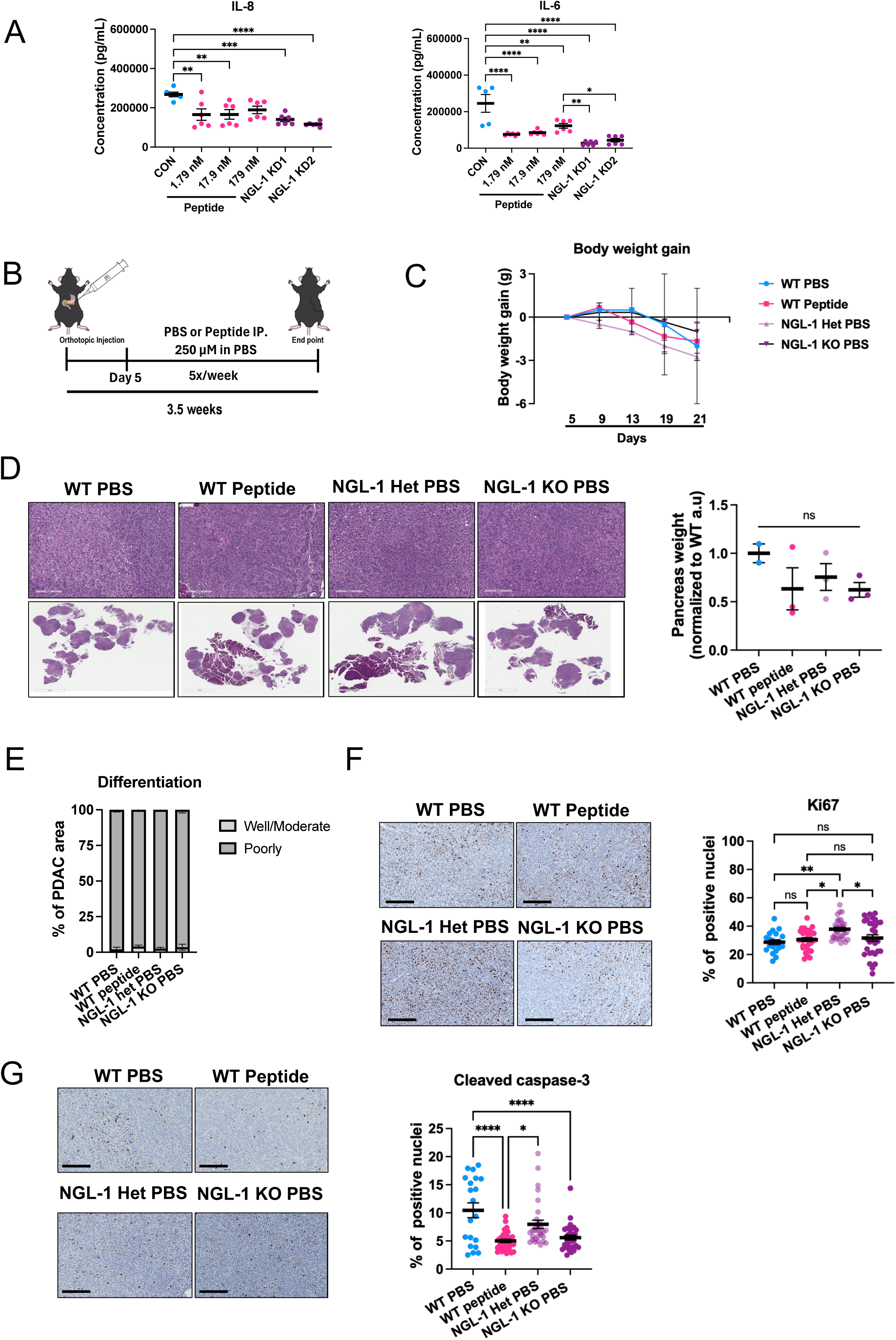
Anti-NGL-1 peptide phenocopies the genetic deletion of NGL-1 *in vitro* and *in vivo*, validating the therapeutical potential of targeting NGL-1. **A)** ELISA detecting the production of IL-8 and IL-6 in the conditioned media of CON CAFs treated with PBS (vehicle control) or peptide at different concentrations (1.79, 17.9 and 179 nM) and NGL-1 KD1 and KD2 CAFs treated with PBS. Note that the treatment with the peptide phenocopies the genetic knockdown of NGL-1, leading to decreased cytokine production compared to CON. One-Way ANOVA, Tukey’s multiple comparisons test. N = 2 independent experiments, each experiment conducted in duplicate or triplicate. **B)** Schematics of the orthotopic allografts using WT, NGL-1 Het and NGL-1 KO mice. Five days post injection with KPC3 cells, mice were treated intraperitoneally with either PBS (10mL/kg) or peptide (250 µM in PBS), 5 times per week until the end of the experiment (3.5 weeks post allografts). **C)** Body weight gain in grams of the different groups throughout the experiment. There were no differences in body weight gain between groups, suggesting that the peptide at this dose and regimen was not toxic to the mice. **D)** Representative images of the pancreas from WT mice treated with PBS or Peptide, NGL-1 Het and NGL-1 KO mice stained with H&E. Scale bars: 7mm and 300 µm. The pancreas weight is shown in the graph, normalized to the average of the WT PBS group, as a measurement of tumor burden. There were tendencies for less tumor burden in the WT peptide, NGL-1 Het and NGL-1 KO groups compared to WT PBS group. N = 2 - 4 mice per group. One-Way ANOVA, Tukey’s multi comparisons test. **E)** Graph showing that there were no differences between the different groups in terms of % of PDAC areas with well/ moderate (light gray) and poorly (dark gray) differentiated tumors, scored by a blinded pathologist. **F)** Representative immunohistochemistry images and quantification of the cells positive for Ki67 (brown) in relation to total number of cells (stained with hematoxylin), as an index of cell proliferation. N = 2 - 3 mice/group, each dot represents the quantification of a single field. Scale bar: 300 µm. One-Way ANOVA, Tukey’s multiple comparisons test. **G)** Representative immunohistochemistry images and quantification of the cells positive for cleaved caspase 3 (brown) in relation to total number of cells (stained with hematoxylin), as an index of cell death. There were more dead cells in the WT PBS compared to the other groups. N = 2-3 mice/group, each dot represents the quantification of a different field. Scale bar: 300 µm. One-Way ANOVA, Tukey’s multiple comparisons test. All graphs depict mean ± SEM.* p < 0.05, ** p < 0.01, *** p < 0.001, **** p < 0.0001. ns = not significant.

**Sup. Figure 20:**
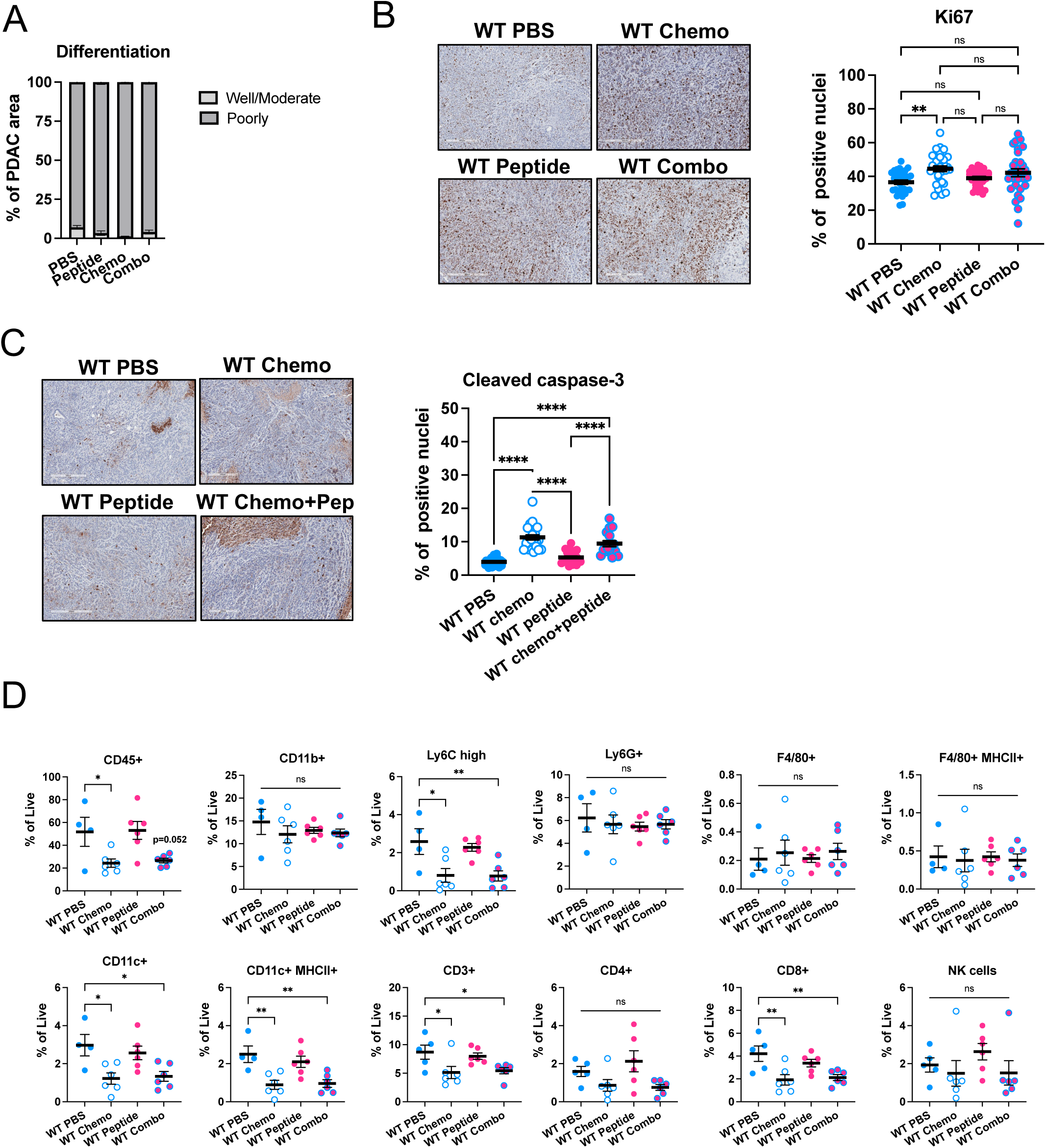
Treatment with chemotherapy leads to increased cell death and decreased infiltration of immune cells in the tumors, independently of the treatment with peptide. **A)** Graph depicting no differences between the different groups (WT mice treated with PBS, chemo, peptide or combo) in terms of % of PDAC areas with well/ moderate (light gray) and poorly (dark gray) differentiated tumors, scored by a blinded pathologist. **B)** Representative immunohistochemistry images and quantification of the cells positive for Ki67 (brown) in relation to total number of cells (stained with hematoxylin), as an index of cell proliferation. There was increased percentage of proliferation in the group treated with chemo, compared to the other groups. N = 3 mice/group, each dot represents the quantification of a single field. Scale bar: 300 µm. **C)** Representative immunohistochemistry images and quantification of the cells positive for cleaved caspase 3 (brown) in relation to total number of cells (stained with hematoxylin), as an index of cell death. There were more dead cells in the groups treated with chemo, independently of the treatment with peptide. N = 3 mice/group, each dot represents the quantification of a different field. Scale bar: 300 µm. One-Way ANOVA, Tukey’s multiple comparisons test. **D)** Immunophenotyping of the tumors from the different groups. The tumors were digested, and the single cell suspensions were analyzed by flow cytometry. All cell populations were gated in singlets and then in live cells. Percentages of each cell subset is in relation to the % of live cells. There were decreased percentages of immune cells in the tumors from the groups treated with chemotherapy, independently of the treatment with peptide. N = 4 – 6 mice/group. One-Way ANOVA, Dunnett’s multiple comparisons test. All graphs depict mean ± SEM. * p < 0.05, ** p < 0.01, **** p < 0.0001. ns = not significant.

