## Supplementary material for "Netrin G1 Ligand is a new stromal immunomodulator that promotes pancreatic cancer": Key resource table

### Key Resources Table

| ANTIBODIES AND KITS | | |
|---|---|---|
| **Immunofluorescence and Immunohistochemistry** | | |
| **Reagent or Resource** | **Source** | **Identifier** |
| NGL-1 Antibody (C1C3) (1:100) | Genetex | # GTX119612<br>RRID:AB_11176528 |
| NGL-1 Antibody (N1N3) (1:150) for IHC or labeled with Qd565 (1:35) for SMI | Genetex | # GTX121508<br>RRID:AB_10721992 |
| Netrin G1 Antibody (D-2) (1:100) for IHC | Santa Cruz | # sc-271774<br>RRID:AB_10707668 |
| Netrin G1 Antibody (N2C1)<br>(1:100) | Genetex | # GTX115637<br>RRID:AB_10625511 |
| Netrin G1 Antibody labeled with Qd655 for SMI (1:150) | SinoBiological | #12313-T16 |
| Pan-cytokeratin (AE1/AE3) Antibody (1:40) | Agilent | # M3515<br>RRID:AB_2132885 |
| Amylase (G10) Antibody (1:200) | Santa Cruz | # sc-46657<br>RRID:AB_626668 |
| Vimentin (EPR3776) antibody (1:200) | Abcam | # ab92547<br>RRID:AB_10562134 |
| ImmPRESS HRP Goat Anti-Rabbit IgG Polymer Detection Kit, Peroxidase | Vector Lab | # MP7451<br>RRID:AB 2631198 |
| Fibronectin antibody (1:200) | Sigma-Aldrich | # F3648<br>RRID:AB_476976 |
| p-FAK [Y397] Antibody, labeled with Qd625 for SMI (1:35) | GeneTex | # GTX24803<br>RRID:AB_ 380431 |
| Donkey anti-rabbit secondary antibody- Alexa Fluor 647 conjugation (1:100) | Jackson ImmunoResearch | # 711-606-152<br>RRID:AB_2340625 |
| Anti-Mouse IgG (whole molecule)–Peroxidase antibody (1:5,000) | Sigma-Aldrich | # A9044<br>RRID:AB_258431 |
| CD4 antibody (1:100) | Cell Signaling | # 25229<br>RRID:AB_2798898 |
| CD8 antibody (1:50) | Cell Signaling | # 98941<br>RRID:AB_2756376 |
| FoxP3 antibody (1:20) | Cell Signaling | # 12653<br>RRID:AB_2797979 |
| NK 1.1 antibody (1:50) | Invitrogen | # MA1-70100<br>RRID:AB_2296673 |
| F4/80 antibody (1:100) | BioRad | # MCA497G<br>RRID:AB_872005 |
| Rabbit monoclonal anti-pSMAD2 (Ser465+467)/SMAD3 (Ser423+425) (clone D27F4) for IHC (1:300) | Cell Signaling | # 8828<br>RRID:AB_2631089 |
| Rabbit monoclonal anti-pSMAD2 (Ser465+467) (clone A5S) for SMI (1:100) | Sigma-Aldrich | #ZRB04953 |
| Ki67 antibody, IHC (1:500) | Cell Signaling | #9662<br>RRID:AB_331439 |
| Cleaved-caspase 3 antibody (1:500) | Cell Signaling | # 9664 |

| | | |
|---|---|---|
| | | RRID:AB_2070042 |
| Ki67 antibody, IF (1:500) | Abcam | # ab15580 RRID:AB_443209 |
| **Flow Cytometry** | | |
| **Reagent or Resource** | **Source** | **Identifier** |
| IFNγ secretion assay detection kit-PE | Miltenyi Biotec | # 130-054-202 |
| Purified Rat Anti-Mouse CD16/CD32 Mouse BD Fc Block | BD Biosciences | RRID:AB_394657 |
| Sytox Blue | Thermo Fisher Scientific | # S11348 |
| Cell tracer CFSE | Invitrogen | # C34554 |
| Purified Hamster Anti Mouse CD28 Clone: 37.51 | BD Biosciences | # 553295<br><br>RRID:AB 394764 |
| Purified Hamster Anti Mouse CD3e Clone: 145-2C11 | BD Biosciences | # 553058 RRID:AB 394591 |
| Anti-mouse CD90.2 (Thy 1.2), FITC, Clone: 53-2.1 | BioLegend | # 140304<br><br>RRID:AB 10642812 |
| Rat Anti-CD45 Monoclonal Antibody, PerCP-Cy5.5, Clone 30-F11 | BD Biosciences | # 550994 RRID:AB 394003 |
| Rat Anti-CD11b, APC-CY7, Clone: M1/70 | BD Bioscience | # 557657 RRID:AB 396772 |
| Rat Anti-Mouse, PE-Cy7, Ly-6G Clone: 1A8 | BD Bioscience | # 560601 RRID:AB 1727562 |
| Rat Anti-Mouse Ly-6G, FITC, Clone: 1A8 | BD Bioscience | # 551460 RRID:AB 394207 |
| Rat Anti-Mouse Ly-6C, APC, Clone: AL-21 | BD Bioscience | # 560595 RRID:AB 1727554 |
| Rat Anti-Mouse F4/80, PE, Clone: T45-2342 | BD Bioscience | # 565410 RRID:AB 2687527 |
| Anti-Mo F4/80, PE-Cy7, Clone: BMB | Thermo Fisher | # 25-4801-82 RRID:AB 469653 |
| Hamster Anti-Mouse CD11c, APC, Clone: HL3 | BD Bioscience | # 553801 RRID:AB 395060 |
| Rat Anti-Mouse I-A/I-E, PE, Clone: M5/114 15.2 | BD Bioscience | # 557000 RRID:AB 396546 |

| | | |
|---|---|---|
| Rat Anti-Mouse CD3 Molecular Complex, FITC, Clone: 17A2 | BD Bioscience | # 555274<br>RRID:AB 395698 |
| Rat Anti-Mouse CD3 Molecular Complex, PE, Clone: 17A2 | BD Bioscience | # 555275<br>RRID:AB 395699 |
| Anti-mouse CD3, APC/Cy7, Clone: 17A2 | BD Bioscience | # 100222<br>RRID:AB 2057374 |
| Mouse Anti-Mouse NK1.1, PE, Clone: PK136 | BD Bioscience | # 557391<br>RRID:AB_396674 |
| Rat Anti-Mouse CD4, APC, Clone: RM4-5 | BD Bioscience | # 553051<br>RRID:AB_398528 |
| Rat Anti-Mouse CD4, PE, Clone: RM4-5 | BD Bioscience | # 553049<br>RRID:AB 394585 |
| Rat Anti-Mouse CD8a, PE-Cy7 Clone: 53-6.7 | BD Bioscience | # 552877<br>RRID:AB 394506 |
| Rat Anti-Mouse Ly-6G and Ly-6C, FITC, Clone: RB6-8C5 | BD Bioscience | # 553127<br>RRID:AB 394643 |
| Anti-mouse Podoplanin, APC/Cy7, Clone 8.1.1 | BioLegend | # 127418<br>RRID:AB 2629803 |
| Anti-mouse CD326 (Ep-CAM), PE, Clone: G8.8 | BioLegend | # 118205<br>RRID:AB 1134172 |
| Anti-mouse/human CD324 (E-Cadherin), PE, Clone: DECMA-1 | BioLegend | # 147304<br>RRID:AB 2563040 |
| Mouse Anti-Mouse CD45.1, APC, Clone: A20 | BD Bioscience | # 558701<br>RRID:AB 1645214 |
| Mouse Anti- Mouse CD45.2, FITC, Clone: 104 | BD Bioscience | # 561874<br>RRID:AB 10894189 |
| Anti-human CD3, FITC, Clone: SK7 | BioLegend | # 981002<br>RRID:AB 2616618 |

| | | |
|---|---|---|
| Anti-human CD8a, Pacific Blue, Clone: HIT8a | BioLegend | # 300928<br>RRID:AB 10612744 |
| Propidium iodide | Invitrogen | # P1304MP |
| **Western Blotting** | | |
| **Reagent or Resource** | **Source** | **Identifier** |
| Anti-Mouse IgG (whole molecule)–Peroxidase antibody (1:5,000) | Sigma-Aldrich | RRID:AB_258431 |
| Goat anti-Rabbit IgG (H+L) Secondary Antibody, HRP (1:10,000) | Thermo Fisher Scientific | RRID:AB_2533967 |
| p-FAK [Y397] (1:200) | Cell Signaling Technology | # 8556<br>RRID:AB_10891442 |
| NFAT1 (D43B1) (1:100) | Cell Signaling Technology | # 5861<br>RRID:AB_10834808 |
| FRA1 (1:2000) | Cell Signaling Technology | # 5281<br>RRID: AB_10557418 |
| FRA2 (1:2000) | Cell Signaling Technology | # 19967<br>RRID: AB_2722526 |
| FosB (1:1000) | Cell Signaling Technology | # 2251<br>RRID: AB_2106903 |
| cFOS (1:1000) | Cell Signaling Technology | # 2250<br>RRID: AB_2247211 |
| c-Jun (1:2500) | Cell Signaling Technology | # 9165<br>RRID: AB_2130165 |
| GAPDH antibody (1:10,000) | Abcam | # ab9485<br>RRID:AB_307275 |
| GAPDH (14C10) (1:10,000) | Cell Signaling Technology | # 2118L<br>RRID:AB_561053 |
| NGL-1 (LRRC4C) (1:200) | LS Bio | # LS-C308814 |
| Palladin iso3 (1:50) | Santa Cruz Biotechnology | # 166563<br>RRID:AB_2236596 |
| Palladin (1:50,000) | Proteintech | # 10853-1-AP<br>RRID:AB_2158782 |
| p-SMAD2/3 (1:100) | Cell Signaling Technology | # 8828S<br>RRID:AB_2631089 |
| SMAD2/3 (1:250) | Cell Signaling Technology | # 8685S<br>RRID:AB_10889933 |
| α-SMA (1:20,000) | Sigma Aldrich | # A2547<br>RRID:AB_476701 |
| β-Catenin (1:5,000) | Cell Signaling Technology | #8480S<br>RRID:AB_11127855 |
| Histone H3 (D1H2) XP antibody (1:10,000) | Cell Signaling Technology | # 4499L<br>RRID:AB_10544537 |
| SureLock™ Tandem Midi Blot Module | Invitrogen | # STM2001 |

| | | |
|---|---|---|
| NuPAGE™ MOPS SDS Running Buffer (20X) | Invitrogen | # NP000102 |
| NuPAGE™ Transfer Buffer (20X) | Invitrogen | # NP00061 |
| NuPAGE™ 4 to 12%, Bis-Tris, 1.0 mm, Midi Protein Gels | Invitrogen | # WG1402BOX |
| NuPAGE™ LDS Sample Buffer (4X) | Invitrogen | # NP0008 |
| SureLock™ Tandem Midi Pre-cut Membranes and Filters, 0.45 µm, PVDF | Invitrogen | # STM2006 |
| 2x Laemmli buffer | BioRad | #1610737 |
| Immobilon Western Chemiluminescent HRP Substrate | EMD Millipore | #WBKLS0500 |
| Immobilon-P PVDF Membrane | EMD Millipore | # IPVH00010 |
| SuperSignal™ West Pico PLUS Chemiluminescent Substrate | Thermo Fisher Scientific | # 34580X4 |
| **BACTERIAL STRAINS** | | |
| **Reagent or Resource** | **Source** | **Identifier** |
| One Shot Stbl3 Chemically Competent E. coli | Thermo Fisher Scientific | # C737303 |
| **CHEMICALS** | | |
| **Reagent or Resource** | **Source** | **Identifier** |
| L-Ascorbic acid | Sigma-Aldrich | #A92902-100G |
| 25% Glutaraldehyde Solution in water | Sigma-Aldrich | #G6257-1L |
| Ethanolamine | Sigma-Aldrich | #E9508-1L |
| Polybrene | Santa Cruz Biotechnology | # sc-134220 |
| TGF-β receptor I inhibitor (SB-431542) | Sigma Aldrich | #616461 |
| Recombinant TGF-β 1 | Sigma Aldrich | #T7039 |
| Blotting grade blocker | Bio-Rad | # 170604 |
| Gelatin | Sigma Aldrich | # G7041 |
| TRI reagent LS | Sigma Aldrich | # T3934 |
| **CELL CULTURE** | | |
| **Reagent or Resource** | **Source** | **Identifier** |
| Fetal Bovine Serum (FBS) | Peak Serum | # PS-FB3 |
| FBS | Sigma Aldrich | # F0926 |
| FBS heat inactivated | Sigma Aldrich | # 12306C |
| DMEM | Sigma Aldrich | #50-013-PB |
| DMEM without L-glutamine and phenol red | Sigma Aldrich | #1145 |
| L-glutamine | Corning | #25-005CI |
| Cytiva HyClone Penicillin Streptomycin 100X Solution | Fisher Scientific | # SV30010 |
| IL-2 | Fox Chase Cancer Center | N/A |
| RPMI | Sigma Aldrich | # R8758 |
| Mycoplasma Detection Kit 50 TESTS | Invivogen | # REP-MYS-50 |
| **CELL LINES** | | |

| Reagent or Resource | Source | Identifier |
|---|---|---|
| KPC3 | Gabitova-Cornell et al, 2020 and Francescone et al, 2021. | PMID: 32976774 and 33127842 |
| Panc-1 | ATCC | RRID:CVCL_0480 |
| Cancer associated fibroblasts 1 (patient derived) | Fox Chase Cancer Center | |
| Cancer associated fibroblasts 2 (patient derived) | Fox Chase Cancer Center | |
| Cancer associated fibroblasts 3 (patient derived) | Fox Chase Cancer Center | |
| Cancer associated fibroblasts 4 (patient derived) | Fox Chase Cancer Center | |
| 293T Cells | ATCC | RRID:CVCL_0063 |
| Phoenix-Amphotropic (φNX) | ATCC | RRID:CVCL_H716 |
| **ORGANISMS/STRAINS (MICE)** | | |
| **Reagent or Resource** | **Source** | **Identifier** |
| C57BL/6J | Jackson Laboratories | # 000664 RRID:IMSR_JAX:000664 |
| B6.SJL-*Ptprc*$^{a}$*Pepc*$^{b}$/BoyJ (B6 CD45.1) | Jackson Laboratories | # 002014 RRID:IMSR_JAX:002014 |
| KPC model: LSL-Kras(G12D/+); Trp53(flox/WT);Pdx-1-Cre | Dr. Kerry Campbell (FCCC) | Original Mice: PMID: 15894267 |
| KC Model: LSL-Kras(G12D/+); Pdx-1-Cre | Dr. Kerry Campbell (FCCC) | Original Mice: PMID: 14706336 |
| NGL-1 KO mice | Dr. Shigeyoshi Itohara (RIKEN Brain Science Institute, Japan) | Original Mice: PMID: 25411505 |
| **PLASMIDS** | | |
| **Reagent or Resource** | **Source** | **Identifier** |
| pLV-CMV-H4-puro vector | Dr. Alexey Ivanov, West Virginia University School of Medicine, Morgantown, WV | N/A |
| psPAX2 | Addgene | # 12260 |
| pCMV-VSV-G | Addgene | # 8454 |
| pBABE-neo-hTERT | Addgene | # 1774 |
| CRISPRi-Puro | Modified from Addgene; (Thakore et al., | # 71236 |

| | | |
|---|---|---|
| | 2015) Francescone et al, 2021, PMID: 33127842 | |
| pMIG-hNFATc2 | Addgene | # 74050 |
| **CRITICAL COMMERCIAL REAGENTS** | | |
| **Reagent or Resource** | **Source** | **Identifier** |
| PureLink™ RNA Mini Kit | Thermo Fisher Scientific | #12183025 |
| TURBO DNA-*free*™ Kit | Thermo Fisher Scientific | # AM1907 |
| Phusion High-Fidelity PCR Master Mix | Thermo Fisher Scientific | # F-531L |
| Odyssey Blocking Buffer (PBS) | LI-COR | # 927-40000 |
| DeadEnd™ Fluorometric TUNEL System | Promega | # G3250 |
| Dnase I | Thermo Fisher Scientific | # EN0525 |
| X-tremeGene9 | Sigma-Aldrich | # 6365787001 |
| U-PLEX Biomarker Group 1 (hu) Assays, SECTOR (1 PL) | Meso Scale Discoveries | # K15067L-1 |
| MILLIPLEX MAP Mouse 32-plex cytokine/chemokine panel | MilliporeSigma | # MCYTMAG-70K-PX32 |
| DuoSet ELISAs (TGF-β, IL-6, IL-8, GM-CSF, mTGF-β, mGM-CSF, mIL-6, mIL-10, mTNFα) | R&D Systems | #DY240, # DY206, # DY208, # DY215, # DY1679, # DY415, # DY406, # DY417, # DY410 |
| DuoSet ELISA Ancillary Reagent Kit 1 | R&D Systems | # DY007 |
| DuoSet ELISA Ancillary Reagent Kit 2 | R&D Systems | # DY008 |
| Sample Activation Kit 1 | R&D Systems | # DY010 |
| Bradford Reagent | Sigma Aldrich | # B6916 |
| Pierce™ BCA Protein Assay Kits | Thermo Fisher | # A55865 |
| DAPI | Sigma Aldrich | # D9542 |
| Worthington Biochemical Corporation Collagenase III | Fisher Scientific | # NC9405360 |
| Amphotericin B | Gibco | # 15290018 |
| Bovine Serum Albumin (BSA) | Sigma Aldrich | # A2153 |
| SiteClick Qdot labeling Kits (565-655) | Thermo Fisher Scientific | #S10450, S10452, S10453. |
| Chromium Single Cell 30 Library, Gel Bead & Multiplex Kit and Chip Kit V3 | 10X Genomics | # PN-1000092 |
| Direct-zol RNA mini prep plus | Zymo Research | # R2071 |
| NEBNext® Ultra™ Directional RNA Library Prep Kit for Illumina | NEB | # E4720L |
| Truseq stranded mRNA library kit | Illumina, Inc. | # 20020595 |
| Superscript II reverse transcriptase | Thermo Fisher Scientific | # 18064014 |

| | | |
|---|---|---|
| SPRIselect beads | Beckman Coulter | # B23318 |
| HiSeq rapid SBS kit v2 (50 cycle) | Illumina, Inc. | # FC-402-4022 |
| Qubit™ 1X dsDNA High Sensitivity (HS) and Broad Range (BR) Assay Kits | Invitrogen | # Q33230 |
| High sensitivity DNA kit | Agilent Technologies | # 5067-4626 |
| Nextseq 2000 high output reagent kit v2.5 | Illumina | # 20024907 |
| High-Capacity cDNA Reverse Transcription Kit | Thermo Fisher Scientific | #4368814 |
| PowerSYBR Green PCR Master Mix | Thermo Fisher Scientific | #4367659 |
| MicroAmp Fast Optical 96 well Reaction Plate with Barcode (0.1 mL) | Thermo Fisher Scientific | #4346906 |
| ProLong™ Gold Antifade Mountant | Thermo Fisher Scientific | #P10144 |
| ImmPACT DAB Substrate Kit, Peroxidase (HRP) | Vector Labs | # SK4105 |
| EZ Prep solution | Roche | # 950–102 |
| Cell conditioning solution (CC1) | Roche | # 950–224 |
| DISCOVERY OmniMap anti-mouse HRP | Roche | # 760-4310 |
| DISCOVERY OmniMap anti-Rb HRP | Roche | # 760-4311 |
| DISCOVERY OmniMap anti-Rt HRP | Roche | # 760-4457 |
| ChromMap DAB detection kit | Roche | # 760-159 |
| Hematoxylin II | Roche | # 790-2208 |
| Bluing reagent | Roche | # 760-2037 |
| Lympholyte M | (Cedarlane) | # CL5031 |
| ACK Lysing Buffer | KD Medical | # RGF-3015 |
| Heparin ammonium salt from porcine intestinal mucosa ≥140 USP units/mg | Sigma Aldrich | # H6279 |
| Gentle MACS Tumor Dissociation Kit | Miltenyi Biotec | #130–096-730 |
| Gentle MACS C tube | Miltenyi Biotec | # 130-093-237 |
| LS tubes | Miltenyi Biotec | # 130-042-401 |
| Dead Cell Removal Kit | Miltenyi Biotec | # 130–090-101 |
| Pan T cell isolation Kit II | Miltenyi Biotec | # 130-095-130 |
| IsoPlexis Single-Cell Adaptive Immune Chip - (M) | Bruker Cellular Analysis | # S-PANEL-1004-4 |
| Recombinant Mouse M-CSF Protein | R&D Systems | 416-ML-050 |
| Lipopolysaccharide (LPS) | Sigma Aldrich | #L2018 |
| Recombinant murine IFNγ | Peprotech | # 315-05 |
| Envision+ polymer system | Dako | #K4000 (anti-mouse) #K4002 (anti-rabbit) |
| Vacutainer Sodium Heparin plastic tubes | BD Biosciences | # 367874 |
| Lymphoprep Medium | StemCell Technologies | # 07801 |
| EasySep Human $CD8^+$ T Cell Isolation Kit | StemCell | # 17953 |

| | | |
|---|---|---|
| | Technologies | |
| EasySep Magnet | StemCell Technologies | # 18000 |
| Dynabeads™ Human T-Activator CD3/CD28 for T Cell Expansion and Activation | Gibco | # 11132D |
| 24 Well glass bottom plates | Cellvis | # P24-1.5H-N |
| Lipids Droplets Fluorescence Assay Kit | Cayman Chemicals | # 500001 |
| Proteinase K, Lyophilized | Millipore | # 70663 |
| GoTaq™ master mixes | Promega | # PRM7122 |
| **OLIGONUCLEOTIDES** | | |
| **Reagent or Resource** | **Source** | **Identifier** |
| mNGL1 qPCR FW TGAGCTGAAATGTCGGGCTT | This paper | https://pga.mgh.harvard.edu/primerbank/ |
| mNGL1 qPCR RV CGTGGGTCATGACTGTTCCA | This paper | https://pga.mgh.harvard.edu/primerbank/ |
| 18s qPCR FW: GGCCCTGTAATTGGAATGAGTC | Primer Bank | https://pga.mgh.harvard.edu/primerbank/ |
| 18s qPCR RV: CCAAGATCCAACTACGAGCTT | Primer Bank | https://pga.mgh.harvard.edu/primerbank/ |
| NGL-1 CRISPRi 1.1 CACCGAAATGGTAAGAGGAATGGG | Compact and highly active next-generation libraries for CRISPR-mediated gene repression and activation | https://doi.org/10.7554/eLife.19760 |
| NGL-1 CRISPRi 1.2 AAACCCCATTCCTCTTACCATTTC | Compact and highly active next-generation libraries for CRISPR-mediated gene repression and activation | https://doi.org/10.7554/eLife.19760 |
| NGL-1 CRISPRi 2.1 CACCGTAAGCCAAAAACTCATCAA | Compact and highly active next-generation libraries for CRISPR-mediated gene repression and activation | https://doi.org/10.7554/eLife.19760 |
| NGL-1 CRISPRi 2.2 AAACTTGATGAGTTTTTGGCTTAC | Compact and highly active next-generation libraries for CRISPR-mediated gene repression and activation | https://doi.org/10.7554/eLife.19760 |
| NFAT1 P1 CRISPRi 1.1 CACCGCGCGCCCGGGGAAGCTGAG | Compact and highly active next- | https://doi.org/10.7554/eLife.19760 |

| | | |
|---|---|---|
| | generation libraries for CRISPR-mediated gene repression and activation | |
| NFAT1 P1 CRISPRi 1.2<br>AAACCTCAGCTTCCCCGGGCGCGC | Compact and highly active next-generation libraries for CRISPR-mediated gene repression and activation | https://doi.org/10.7554/eLife.19760 |
| NFAT1 P2 CRISPRi 1.1<br>CACCGGCGATCCGGCTTACTCCAG | Compact and highly active next-generation libraries for CRISPR-mediated gene repression and activation | https://doi.org/10.7554/eLife.19760 |
| NFAT1 P2 CRISPRi 1.2<br>AAACCTGGAGTAAGCCGGATCGCC | Compact and highly active next-generation libraries for CRISPR-mediated gene repression and activation | https://doi.org/10.7554/eLife.19760 |
| NFAT1 FW Xba<br>TAGGTATCTAGAatgaacgcccccgagcg | This paper | N/A |
| NFAT1 RV Xho<br>TAGGTACTCGAGttacgtctgatttctggcaggagg | This paper | N/A |
| CMV FW<br>CGCAAATGGGCGGTAGGCGTG | Francescone et al. 2021 | PMID: 33127842 |
| U6 FW<br>GAGGGCCTATTTCCCATGATT | Francescone et al. 2021 | PMID: 33127842 |
| **INSTRUMENTS** | | |
| **Reagent or Resource** | **Source** | **Identifier** |
| NanoDrop 1000 or 2000 | Thermo Fisher Scientific | http://www.thermofisherscientific.com |
| Nikon Eclipse TE2000U | Nikon | http://www.nikon.com/ |
| MSD SECTOR Imager 2400 | Meso Scale Discoveries | https://www.mesoscale.com |
| BioPlex 200 biomarker analyzer | Bio-Rad | https://www.bio-rad.com/en-us/product/bio-plex-200-systems?ID=715b85f1-6a4e-41b3-b5d9-80202d779e13 |
| Spark® Multimode Microplate | Tecan | https://www.tecan.com |
| SpectraMax ID3 MicroPlate reader | Molecular Devices | https://www.molecular |

| | | |
|---|---|---|
| | | devices.com/sites/default/files/en/assets/user-guide/br/spectramax-id3-userguide-5054747h.pdf |
| HiSeq2500 System | Illumina, Inc. | www.illumina.com |
| Agilent 2100 bioanalyzer | Agilent Technologies | www.agilent.com<br># DE34903146 |
| Qubit 3.0 fluorometer | Invitrogen | # Q33216 |
| StepOnePlus Real Time-PCR System | Applied Biosystems/Thermo Fisher Scientific | https://www.thermofisher.com/order/catalog/product/4376600#/4376600 |
| Leica SP8 DIVE confocal/multiphoton microscope system | Leica | https://www.leica-microsystems.com/products/confocal-microscopes/p/dive/ |
| Multiphoton IR laser Chameleon Vision II | Coherent Inc | https://www.coherent.com/ |
| Nikon Eclipse Ti2-E Inverted Microscope Imaging System | Nikon | https://www.microscope.healthcare.nikon.com/products/inverted-microscopes/eclipse-ti2-series |
| Nikon A1 camera | Nikon | https://www.microscope.healthcare.nikon.com/products/confocal-microscopes/a1hd25-a1rhd25 |
| Nikon Eclipse 50/55i clinical microscope | Nikon | https://www.microscopyu.com/museum/eclipse-50-55i |
| Nikon DS-Fi1 camera | Nikon | https://www.microscope.healthcare.nikon.com/products/cameras |
| Roche Ventana Discovery Ultra automated autostainer | Roche | https://diagnostics.roche.com/us/en/products/product-category/immunohistochemistry--ihc-/discovery-ultra.html |
| GentleMACS Dissociator | Miltenyi Biotec | # 130-093-235 |
| QuadroMACS separator | Miltenyi Biotec | # 130-090-976 |
| LSRII flow cytometer | BD Biosciences | https://www.bd.com/resource.aspx?IDX=17868 |
| BD FACS Aria II | BD Biosciences | |
| Isospark | Bruker Cellular Analysis | https://brukercellularanalysis.com/products/instruments/isospark- |

| | | |
|---|---|---|
| | | system/ |
| PowerEase™ Touch Power Supply | Invitrogen | # PS0350 |
| ChemiDoc Imaging System | BioRad | # https://www.bio-rad.com/en-us/product/chemidoc-imaging-system?ID=OI91XQ15 |
| Aperio ScanScope CS2 Scanner | Leica | https://www.leicabiosystems.com/us/digital-pathology/scan/aperio-cs2/ |
| Chromium Controller | 10X Genomics | https://www.10xgenomics.com/instruments/chromium-controller |
| Nextseq 2000 | Illumina | https://www.illumina.com/systems/sequencing-platforms/nextseq-1000-2000.html |
| **SOFTWARE/ALGORITHMS** | | |
| **Reagent or Resource** | **Source** | **Identifier** |
| ImageJ | NIH | https://imagej.nih.gov/ij/ (RRID:SCR_003070) |
| Metamorph 7.8.1.0 Image Analysis Software | Molecular Devices | http://www.moleculardevices.com/Products/Software/Meta-Imaging-Series/MetaMorph.html (RRID:SCR_002368) |
| SMIA-CUKIE | Franco-Barraza et al., 2017 | https://github.com/cukie/SMIA (RRID:SCR_014795) |
| FlowJo | FlowJo LLC | https://www.flowjo.com/solutions/flowjo (RRID:SCR_008520) |
| BD FACS DiVa Software v 8.0.1. | BD Biosciences | https://www.bdbiosciences.com/en-us/products/software/instrument-software/bd-facsdiva-software |
| OrientationJ plugin for ImageJ | Rezakhaniha et al, 2012 | http://bigwww.epfl.ch/demo/orientation/ (RRID:SCR_014796) |
| Fiji | ImageJ | https://fiji.sc/ |
| MSD Workbench 4.0 software | Meso Scale Discoveries | https://www.mesoscale.com |
| BioPlex manager software | Bio-Rad | https://www.bio-rad.com/en-us/category/bio-plex-software?ID=45938d9 |

| | | |
|---|---|---|
| | | d-c2ec-4ae4-9ed3-e7358a98d30b |
| Graph Pad Prism 10.0 software | GraphPad Software | https://www.graphpad.com/scientific-software/prism/ (RRID:SCR_000306) |
| Adobe Photoshop CS6 13.0.1 | Adobe | http://www.adobe.com/fr/products/photoshop.html (RRID: SCR_014199) |
| Nuance 3.0.1 | Caliper Life Sciences/Perkin Elmer | https://www.moleculardevices.com/Products/Software/Meta-Imaging-Series/MetaMorph.html (RRID: SCR_002368) |
| Aperio Imagescope | Leica | https://www.leicabiosystems.com/sites/default/files/media_document-file/2021-02/MAN-0001-Rev-Q.12.4_pdf.pdf |
| Leica Application Suite X 3.5.5 | Leica | https://www.leica-microsystems.com/products/microscope-software/p/leica-las-x-ls/ |
| Illumina base space | Illumina, Inc. | (https://basespace.illumina.com |
| CellRanger algorithm | 10X Genomics | https://10xgenomics.com/support/software/cell-ranger/latest/algorithms-overview/cr-gex-algorithm |
| Loupe browser | 10X Genomics | https://www.10xgenomics.com/support/software/loupe-browser/latest |
| IsoSpeak | Bruker Cellular Analysis | https://brukercellularanalysis.com/resource/isospeak-manual-2/ |
| KOBAS | PMID: 21715386 | http://kobas.cbi.pku.edu.cn |
| Morpheus | Broad Institute | https://software.broadinstitute.org/morpheus/ |
| Ingenuity | Qiagen | https://digitalinsights.qiagen.com/products-overview/discovery-insights-portfolio/analysis-and- |

| | | |
|---|---|---|
| | | visualization/qiagen-ipa/?gad_source=1&gclid=CjwKCAjwtqmwBhBVEiwAL-WAYc6YvMXhRr5wnSUydcLmlmOQmOpVdHAOZDAuBwfb2zEyYWxj3O_GchoCYGwQAvD_BwE |
| Seurat v4 | | https://satijalab.org/seurat/articles/get_started.html |
| Word Cloud Generator | | https://www.freewordcloudgenerator.com/ |
